# An improved workflow for the quantification of orthohantavirus infection using automated imaging and flow cytometry

**DOI:** 10.1101/2023.12.21.572760

**Authors:** Laura Menke, Christian Sieben

**Affiliations:** Nanoscale Infection Biology, Helmholtz Centre for Infection Research, Braunschweig, Germany; Institute of Genetics, Technische Universität Braunschweig, Spielmannstr. 7, Braunschweig 38106, Germany

**Keywords:** orthohantavirus, virus quantification, imaging, bunyavirales

## Abstract

Determination of the infectious titer is a central requirement when working with pathogenic viruses. The plaque or focus assay is commonly used but a labor- and time-consuming approach to determine the infectious titer of orthohantavirus samples. We have developed an optimized virus quantification approach that relies on the fluorescence-based detection of the orthohantavirus nucleocapsid protein (N) in infected cells with high sensitivity. We present the use of flow cytometry but highlight fluorescence microscopy in combination with automated data analysis as an attractive alternative to increase the information retrieved from an infection experiment. Additionally, we offer an open-source software equipped with a user-friendly graphical interface, eliminating the necessity for advanced programming skills.

## Introduction

Orthohantaviruses are enveloped viruses that contain three circular single-stranded RNA genome segments of negative sense, namely the small (S-Segment), medium (M-Segment) and large segment (L-Segment) [1, 2]. Encoded on the S-segment is the nucleocapsid (N) protein [3], which is often used for virus quantification due to its early and abundant production [4-6]. Research on orthohantaviruses started with the discovery of Hantaan Virus (HTNV) in 1978 [7] and morphological and genetic insights on HTNV followed as soon as virus propagation and titration could be established in cell culture systems [8, 9]. Here, especially the viral adaptation to Vero E6 cells was an important milestone [10]. Since then, several protocols for orthohantavirus titration in cell culture have been developed. The widespread foci assay titrates the number of infectious particles by detecting the viral N protein in infected cells [9, 11, 12]. The foci assay is closely related to the plaque assay where plaques of dead cells are visualized as empty spots following a general cell staining procedure (i.e., with methylene blue). Since orthohantaviruses are non-lytic, infected cells are detected via the expression of viral proteins. Specifically, a semisolid overlay medium ensures that after infection viral spread can only occur to neighboring cells, which leads to a patch of infected cells that can be counted as a focus [9, 11, 12]. A derivative of the fluorescence-based foci quantification protocol is the viral antigen detection via chemiluminescence which potentially offers improved sensitivity [13]. One drawback of the foci assay is its labor and time-consuming protocol, which takes 5-11 days depending on the viral strain [13]. Recently, a flow cytometry-based titration assay was established using Andes orthohantavirus (ANDV). This assay has been shown to reduce the time required for the detection of viral replication during infection [14]. The authors state that the tested FOCI assays can be compared with their improved protocol, in which ANDV detection was possible as early as 6 hours post infection.

Here we provide an improved virus quantification protocol based on fluorescence imaging and flow cytometry that is compatible with other orthohantavirus strains such as PUUV and TULV. While earlier time points can also be detected by fluorescence microscopy, our data suggest that PUUV and TULV should best be quantified after 48 hours. We provide an automated imaging workflow together with a data analysis package, which allows to routinely quantify large amounts of single-cell data (5,000-10,000 cells) in a time-efficient way. As an additional advantage, titration by imaging allows to extract structural information at the same time without the burden of an extra experimental setting, a major advantage, which even allow discrimination between early and late stages of infection.

## Material and Methods

### Cells, Viruses and Reagents

Vero E6 cells were purchased from ATCC and grown in DMEM (Gibco: DMEM (1x) + GlutaMAX™-I, [+] 2.5g/l D-Glucose, [-] Pyruvat) containing 10% fetal calf serum (FCS) (Sigma) and 1% Penicillin/Streptomycin (Pen/Strep) (Gibco). Deionized water processed by an EMD Millipore™ Milli-Q™ advantage A10 ultrapure water purification system served as basis for all buffers and solutions. All disposable cell culture materials were purchased from Sarstedt, TPP, Gibco, Thermo Fisher Scientific and Greiner bio-one. Nucleocapsid (N) proteins were detected using primary antibodies 5E11 (Abcam; ab34757) [18], or monoclonal antibody anti-TULV-1 generously provided by Jens Peter Teifke and Sven Reiche (Friedrich-Loeffler-Institut, Greifswald, Germany). We used Alexa Fluor 568 and Alexa Fluor 488 secondary antibodies (Thermo).

### Orthohantavirus propagation

Hantaviruses were propagated on Vero E6 cells (ATCC) under strict standard operating procedures using biosafety level 2 (BSL2) facilities and practice. For virus propagation, T75 flasks with Vero E6 cells were seeded to reach a density of 80-90% confluence after overnight incubation at 37°C in 5% Carbon dioxide (CO_2_). The cells were infected using a multiplicity of infection (MOI) of 0,01-0,1 in DMEM + 2% FCS + 1% Pen/Strep. After 1 hour at 37°C (+ 5% CO_2_), the inoculum was removed and the medium replaced with fresh DMEM + 2% FCS + 1% Pen/Strep. The infected cells were further incubated for 7-10 days, whereby after day 4 the supernatant can be collected on a daily bases and later either quantified passage per passage or pooled by using concentration methods. Collected supernatants were centrifuged for 30 min at 1200-2000 g at 4°C to remove cell debris, then transferred into 50-100 kDa filtration units (Amicon) (Sartorius) and centrifuged at 4000 g until volume decrease to 1-3 ml. The retentates were mixed, resuspended and frozen at -80 °C until further use.

### Preparation of glass cover slips for microscopy

200 ml washing solution containing 60% of hydrochloric acid (HCl) and 40% ethanol (EtOH) were mixed and 120 ml of HCl were added and heated to 37°C. The solution was filtered through a pleated filter. Cover Slips are washed carefully in a 1 l glass bottle on a roll shaker for at least 30 min. Afterwards the HCl/EtOH mix was discarded. Cover Slips were washed 3 times with water and then dried on whatman-paper. Finally, the slides were sterilized at 180°C.

### Preparation of a fluorophore-conjugated primary antibodies

20 μl of the antibody 5E11 (Abcam; ab34757) [18], together with 20 μl phosphate-buffered saline (PBS; filtered) (Gibco), 10μl NaHCO_3_ (pH 8-8,5; 0,5 M) and 0,5μl dye conjugate (CF-488A Dye; Biotium) (4 μg; 64 μM final; dye overload ca. 1:50) were incubated for 30 min at room temperature in the dark. NAP-5 Sephadex columns (Amersham) were rinsed with Phosphate buffered saline (PBS), before the sample was applied and eluted with PBS. Run through was collected drop-wise in a 96-well plate (well after well) and the dye/protein content of each well was measured using a NanoDrop (Thermo scientific). The wells with the highest concentration were pooled and measured again to get the final concentration (protocol is based on [19]).

### Immunofluorescence assay (IFA) for flow cytometry

100,000 Vero E6 cells were seeded in a 24-well plate (Corning) and incubated with DMEM + 10% FCS + 1% Pen/Strep for 24h at 37°C in 5% CO_2_. Dependent on the type of experiment, either a MOI of 1 was used or a fixed volume (5 μl or 50 μl). For infections, the cell layer should be 80 % confluent. The used virus volume was filled up to 300 μl per well with DMEM + 2 % FCS + 1 % Pen/Strep and incubated for 1 hour at 37 °C. The cells were washed twice with PBS and further incubated for 48 h with DMEM + 2 % FCS + 1 % Pen/Strep. After washing with PBS, the cells were treated with Accutase (Sigma-Aldrich) for 5 min at 37°C. The detached cells were transferred into a tube, which was fill up with PBS and then centrifuged (all following centrifugation steps: 2000 g for 5 min at 4 °C). Near-IR fluorescent reactive dye (Thermo Fisher) was used 1:1000 in PBS, and added for 30 min at room temperature (RT). As a washing step, the tubes were filled up with PBS and centrifuged. 4 % Paraformaldehyd (PFA) was used for 30 min at RT to fixate the samples, followed by a dilution/ washing step and centrifugation. The cells were permeabilized for 15 min using 0.5 % Saponin in PBS at RT, followed by a dilution/ washing step and centrifugation. Antibodies were added in a buffer consisting of PBS, 2 % FCS, 0.4 % 0.5M Ethylenediaminetetraacetic acid (EDTA) and 0.5 % Saponin and incubated for 1 hour at RT. After each antibody-staining step two washing steps were performed. The samples were analyzed using a BD LSR-II flow cytometer using the software BD FACSDiva. Data were further processed using FlowJo_V10.9.0.

### Immunofluorescence assay (IFA) for microscopy

100,000 Vero E6 cells were seeded in a 24-well plate (Corning) containing 12 mm coverslips (Epredia). Alternatively, 50,000 cells were seeded in a 48-well plate containing 7 mm coverslips. The cells were incubated in DMEM + 10% FCS + 1% Pen/Strep for 24 h at 37°C in 5% CO_2_. Dependent on the type of experiment, either a MOI of 1 was used or a fixed volume (5 μl or 50 μl). For infections, the cell layer should be 80% confluent. The used virus volume was filled up to 300 μl per well with DMEM + 2% FCS + 1% Pen/Strep and incubated for 1 h at 37 °C. The cells were washed with PBS and further incubated for 48 h with DMEM + 2%FCS + 1% Pen/Strep. 4% PFA was used for 15 min at RT to fix the cells (Thermo Fisher). As a quenching solution we used 300 mM glycine (Serva) in PBS and then permeablized the cells in 0.1% TritonX100 (Sigma) in PBS for 30 min at RT. As a blocking solution, 1% bovine serum albumin (BSA) (Sigma Aldrich) was mixed with 0.1% Tween20 (Sigma Aldrich) in PBS and added for 30 min at RT. All following antibodies or dyes were diluted in the BSA-Tween-PBS-blocking solution. The first antibody, as well as the secondary antibody was incubated for 1 h at RT, followed by 20 min incubation with Phalloidin (1:400 dilution pf a 66 μM stock solution; manufacturer’s specification) and 10 min of Hoechst (1 μg/μl) staining. Between all steps, the samples were washed three times with PBS and additional once with water before the samples were mounted in ProLong TM Gold antifade reagent (Thermo Fisher). The coverslips were dried for at least 24 h. As a sealant CoverGripTM (biotium) was used and dried for at least 24 h.

### Fluorescence microscopy

Fluorescence microscopy was performed on a Nikon Ti2-Eclipse microscope equipped with the spinning disk confocal module CSU-W1 (Yokogawa) as well as a LED widefield illumination light source (*p*E-4000, CoolLED). Images were acquired with a S Plan Fluor 20x and Plan Fluor 60× oil objective (Nikon), a PCOedge 4.2LT or a Zyla 4.2 sCMOS camera (Andor, Belfast, UK) and 405/488/561/638 nm laser lines (Omicron, Rodgau-Dudenhofen, Germany) controlled by NIS-Elements software. Z-stack images were acquired with a step size of 300 nm.

### Data processing and statistical analyses

Image analysis was carried out using CellProfiler and NIS-Elements (Nikon). Further post-processing was done using our custom software Datalyze (**Supplementary Software**). Further data processing steps and statistical analyses were carried out in NIS-Elements, Inkscape, Excel 2010 and Graphpad Prism 9. Graphical illustrations were created with BioRender.com.

## Results

### PUUV and TULV grow to low titers requiring time-consuming quantifications

Recently, imaging-based virus quantification has become increasingly popular due to its ease of use and the additional amount of information contained in microscopy images [20, 21]. Pseudotyped or recombinant viruses encoding a cytosolic fluorescent reporter further enable quick detection of infected cells. However, some orthohantavirus strains grow to relatively low titers, compared with other enveloped RNA viruses and a recombinant system is not yet available. In our hands, PUUV reached a 100-fold lower titer compared to influenza virus A/Puerto Rico/8/34 and La Crosse encephalitis virus (LACV), another Bunyavirus. Compared to VSV, the titer was even 10^4^-fold lower (**Supplementary Figure 1**). The typical proliferation time is also considerably longer for orthohantaviruses, which typically require on average 10 days of incubation. Consequently, the small amount of virus material available, is typically used up quickly, leading to a constant amplifications process, followed by quantifications of viral titers for again up to 10 days. We thus set out to optimize the orthohantavirus quantification methods to reduce the amount of infectious material and save time compared to the standard method.

### An improved flow cytometry workflow to quantify PUUV and TULV with an optimized time point and reduced material requirement

Quantifying orthohantavirus infection using flow cytometry has been described as early as 6 hours post-infection for ANDV infection [14]. We thus first tested this early time point after PUUV and TULV infection using the described protocol but could not detect any infected cells (**Supplementary Figure 2**). Instead, 48 hours post-infection enabled a robust detection of infected cells (**Figure 1 B-E**) suggesting that the time point requires optimization depending on the used virus strain. During our optimization, we could further reduce the required cell number by 80%. While the previous protocol describes the use of a 6-well plate format (well surface area: 9.6 cm^2^), we could adjust the protocol to a 24-well plate format (well surface area: 1.9 cm^2^). For the small amount of required starting material, which still allowed a measurement of 10,000 cells, we want to highlight several critical steps in our protocol (**Figure 1 A**). First, the cells were detached from the well plate using accutase, which we found to be superior compared to trypsin or cell scrapers which are often used in flow cytometry assays. The cells were collected in a buffer containing EDTA to prevent the cells from sticking together. Then, a live-dead marker was included which enables selecting only intact cells for the final analysis. Following this staining step, the cells were fixed in 4% PFA. The subsequent antibody staining requires cell permeabilization, for which we recommend using saponin, rather than detergents such as Triton X-100. Orthohantavirus infection is then detected using primary antibodies against the viral nucleocapsid protein N diluted in a buffer containing saponin to retain the cells permeable for the required intracellular staining. This step is followed by two washing steps and incubation with a secondary antibody, again followed by another two washing steps. Here, the use of a secondary antibody amplifies the signal since one primary antibody can bind more than one secondary antibody. However, due to the many washing steps we observed a strong loss of cells, on average 3-10% per washing step (**Supplementary Figure 3**). The use of a direct fluorophore-conjugated antibody can help overcome this limitation while remaining to use small cell culture volume (24 well). As we describe in the methods section, the anti-N primary antibody was then conjugated to a selected bright dye in a separate process. Using a pre-conjugated primary antibody saved about 75 min during each of the quantification experiments, as well as at least two washing steps and the associated loss in cell number. As an additional benefit, the background signal was also reduced when compared to the use of a combined primary/secondary antibody in a mock infected control sample (**Supplementary Figure 4**). Finally, following a simple gating strategy to select only live single cells (**Supplementary Figure 5**), our optimized protocol allowed to count cells infected with PUUV and TULV after 48 h using a small number of input cells.

**Figure 1:**
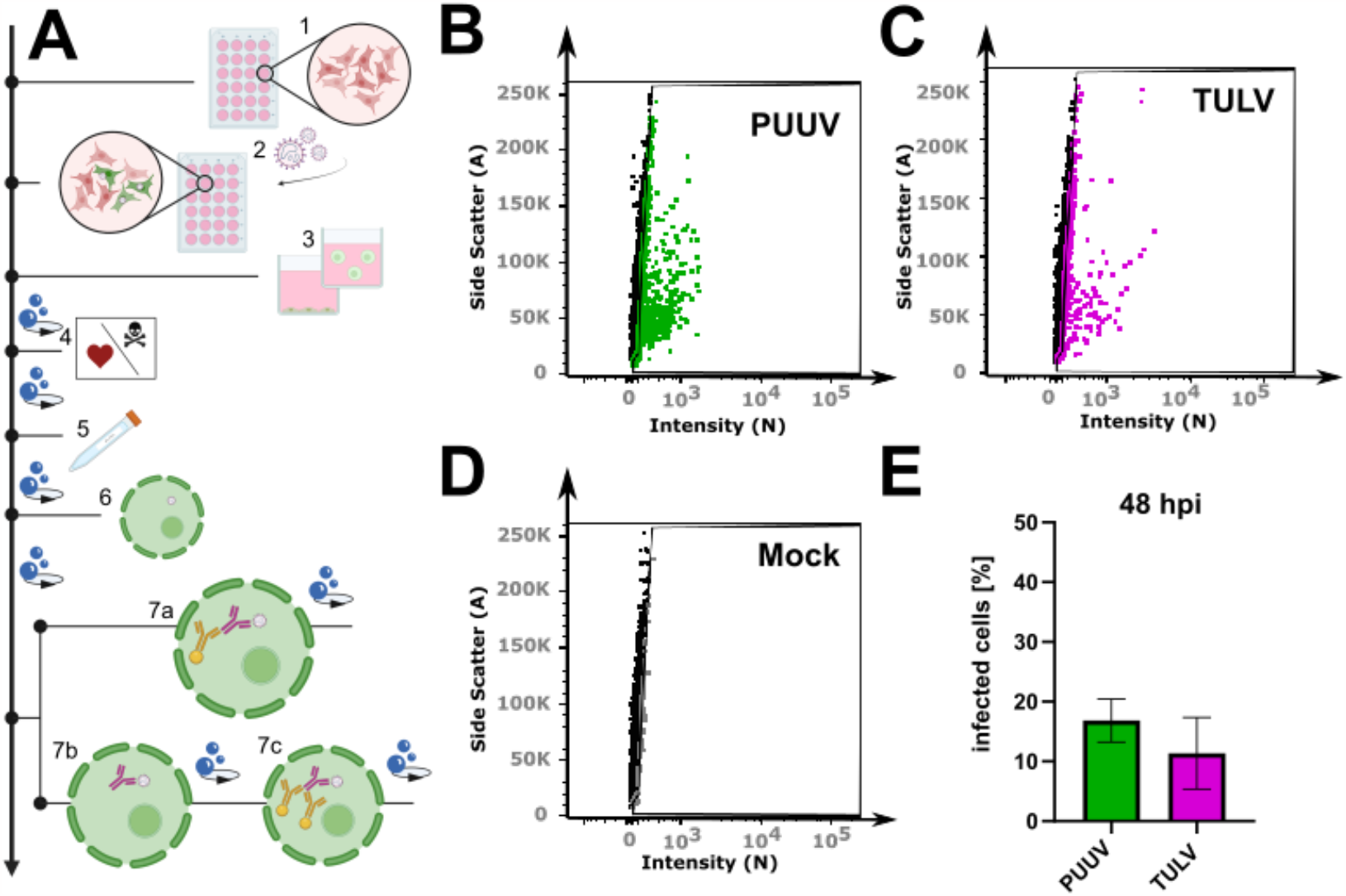
Flow cytometry workflow for quantifying orthohantavirus infection in cell culture: (A) Schematic overview of the protocol: (1) Cell seeding, (2) infection with orthohantavirus for 1 hour, (3) cell detachment with accutase, (4) Live-Dead staining, (5) fixation with paraformaldehyde (PFA), (6) cell permeabilisation with Saponin, (7) staining with antibodies: either with a conjugated antibody (7a) or with a 1^st^ (7b) and a 2^nd^ (7c) antibody. (B –D) Exemplary gate settings for quantification of newly propagated virus and their analysis in (E) as a bar graph displaying the percentage of infected cells (%). Shown are mean ± SD, N = 3.

### Microscopy in combination with automated image analysis as a highly sensitive approach for quantifying orthohantavirus infection phenotypes

The mere counting of infected cells based on the presence of a viral protein leaves out the opportunity to use the positional and structural information of the antibody-labelled proteins within the cellular context. Hence, we wondered if we could also use an imaging-based approach to quantify infection with PUUV and TULV in cell culture.

In general, quantification via microscopy, requires even lower input material due to the specific and exhaustive imaging of whole cell populations and the low loss during sample preparation. We routinely use either 12 mm coverslips in a 24-well or 7 mm coverslips in a 48-well plate (well surface area: 1.1 cm^2^). Compared to a 6-well format, this results in a reduction of 89 % with respect to the required reagents. Vero E6 cells were infected for 48 h, as described above, then fixed with 4% PFA directly on the glass slides. No detachment and centrifugation steps were required thus minimizing the cell loss compared to the flow cytometry workflow (**Supplementary Figure 5**). Nevertheless, on a 7 mm coverslip, it is still possible to acquire 15-25 non-overlapping images using a 20x objective, resulting in a cell count of 5,000-10,000 cells for the evaluation, which is comparable with the flow cytometry workflow described above (**Figure 2**). Following fixation, the cells were permeabilized with Triton X-100 and blocked with a BSA-containing buffer, which was also used for the antibody dilutions. Since there is no risk of losing cell material during the following washing steps, we recommend using primary/secondary antibodies to increase the signal to noise. To enable single cell segmentation later on, we then stained the cell body using phalloidin as well as the nuclear DNA using Hoechst (**Figure 2 D-K**). Images were taken at 20x magnification, whereby one image corresponds to approximately 200-500 Vero E6 cells (**Figure 2 B**) that could still be segmented adequately as shown by the corresponding software-based mask in **Figure 2** C. For image processing, we used the licensed program NIS-Elements AR, as well as the open-source tool CellProfiler [22]. Both programs first segment the cell nuclei (**Figure 2** F), followed by detection of the cell boundaries (**Figure 2** G and J), whereby not only epithelial cells can be recognized, but also fibroblast-like ones (**Supplementary Figure 6**). Following single-cell and nuclear recognition, segmentation of the N protein (**Figure 2** H and K) and association of the different masks can be used to distinguish infected from non-infected cells. Due to the mask association (per single-cell), virus particles non-specifically bound to the glass slide can easily be ignored. We provide a detailed walkthrough of the image analysis and refer to our short tutorial (**Supplementary Software**, sample datasets can be downloaded from 10.5281/zenodo.10417810). Using this protocol, we then analyzed cells infected with PUUV and TULV at different time points post infection (6h, 12h, 24h and 48h). Interestingly, a positive infection with PUUV as well as with TULV could already be detected at 6h post infection (**Figure 3** C), which was not possible using flow cytometry **(Supplementary Figure 2**). After 48h we found similar infection rates as measured by flow cytometry when using the same input virus (**Supplementary Figure 7**). Over the course of infection, we observed a correlation between the number of infected cells and the intensity of the segmented signal (**Figure 3A, B**). This observation implies that within the initial 48 hours of infection, the two parameters are linked and the quantification mainly monitors the primary infection. To test this conclusion, we used the supernatant of all time points to infect new Vero E6 cells for a subsequent 48h period. Interestingly, we observed infection rates at background levels for all supernatant samples except the 48h time point (**Supplementary Figure 8**) supporting our conclusion that we are effectively detecting the ongoing primary infection process (**Figure 3**). Based on these findings, we recommend the 48h time point post-infection, characterized by the highest signal intensity, as the optimal time frame for quantifying PUUV and TULV stocks.

**Figure 2:**
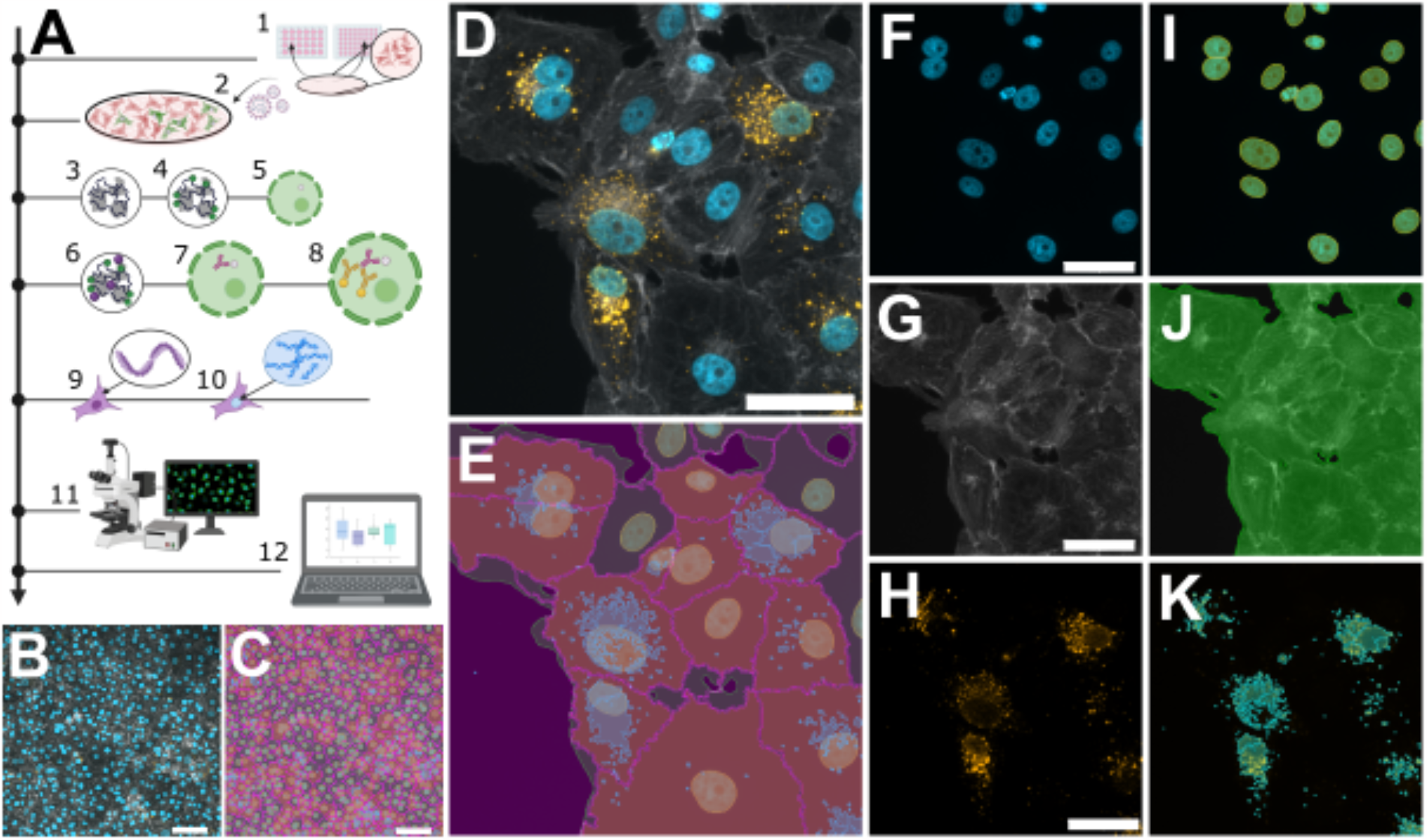
Microscopic workflow for quantifying orthohantavirus infection in cell culture. (A) Schematic overview of the protocol: Cell seeding (1), infection with orthohantavirus for 1 hour (2), cell fixation with paraformaldehyde (PFA) (3), blocking of unreacted aldehydes with glycine solution (4), permeabilisation with Triton-X100 (5), blocking of unspecific binding sites using a BSA-containing buffer (6), staining with 1^st^ antibody against viral nucleocapsid protein (N) (7) and staining with a 2^nd^ antibody (8), Phalloidin staining to label filamentous actin (9) and Hoechst for labelling dsDNA (10), imaging via fluorescence microscopy (11) and analysis via custom software tools (12). (B) and (C) show exemplary images recorded with 20x magnification showing ∼500 cells; (D-K) show pairwise original images together with their masks resulting from software-based segmentation: (D) shows a merged image of three channels, individual channels in (F-H): Hoechst (F) and it’s mask for nuclei recognition (I), Phalloidin staining (G) for recognizing cell bodies or cell boundaries, Anti-N staining (H) to reveal viral particles (K). The overlay of I, J and K is shown in E. Scale bars = 50 μm.

**Figure 3:**
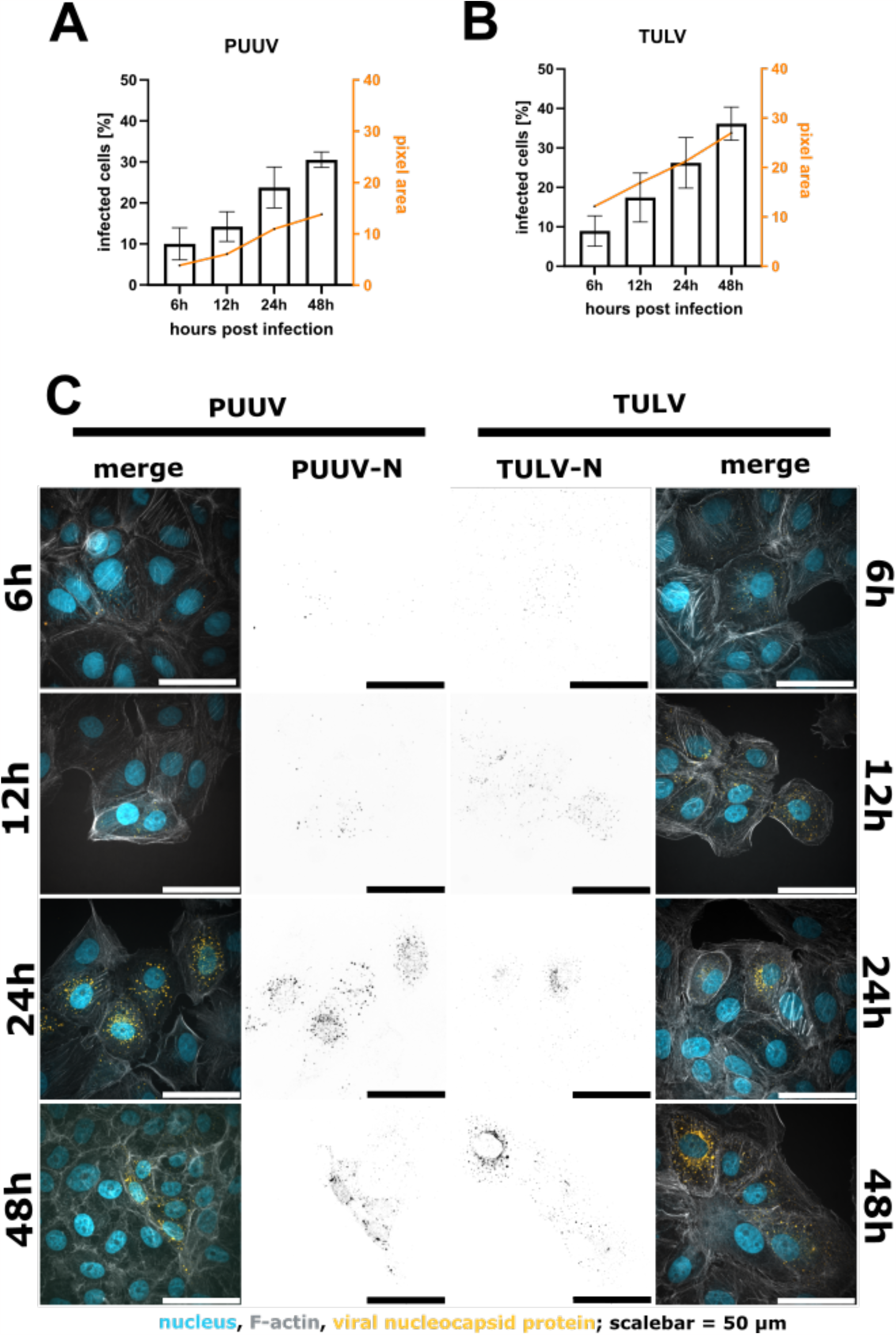
Quantification of infected Vero E6 cells 6h, 12h, 24h and 48h post infection with either PUUV (A) or TULV (B). A comparison of the percentage of infected cells shows an increase in detected cells after 48h compared to 6h. We observed a correlation of the number of infected cells and the intensity per pixel. Shown are mean ± SD, n = 3. Exemplary images in (C). Merged image show nuclei in blue, F-actin in grey and the viral N protein in orange. Viral N is also shown in black as a separated image in the middle of the image panel.

### Custom analysis software tools allow to quickly visualize the image data and calculate infection rates

The image analysis pipeline discussed above produces signal intensity datasets in tabular format, which require further processing. To ease this step, we provide two version of our open-source software equipped with a graphical user interface (**Figure 4** B-D), so that no programming skills are required. After download, the software to process CellProfiler or Nis Elements data is ready to use without further installation. The raw data can be visualized as a scatterplot (**Figure 4** C), whereby data sets are color-coded and can be further summarized as average percentage of infected cells (**Figure 4** D). For more detailed information, please refer to the provided software manual packaged together with the **Supplementary Software**. Finally, to convert the fraction of infected cells into infectious particles per milliliter, three parameters are necessary: 1) the percentage of infected cells as given by our analysis tool (e.g. 50.9 % converted as a fraction 0.509), 2) the used cell numbers (e.g. 100,000 cells per well), and 3) the amount of virus suspension (e.g. 5 μl per well) used to include the used dilution (1:200) to finally calculate infectious particles per ml.

**Figure 4:**
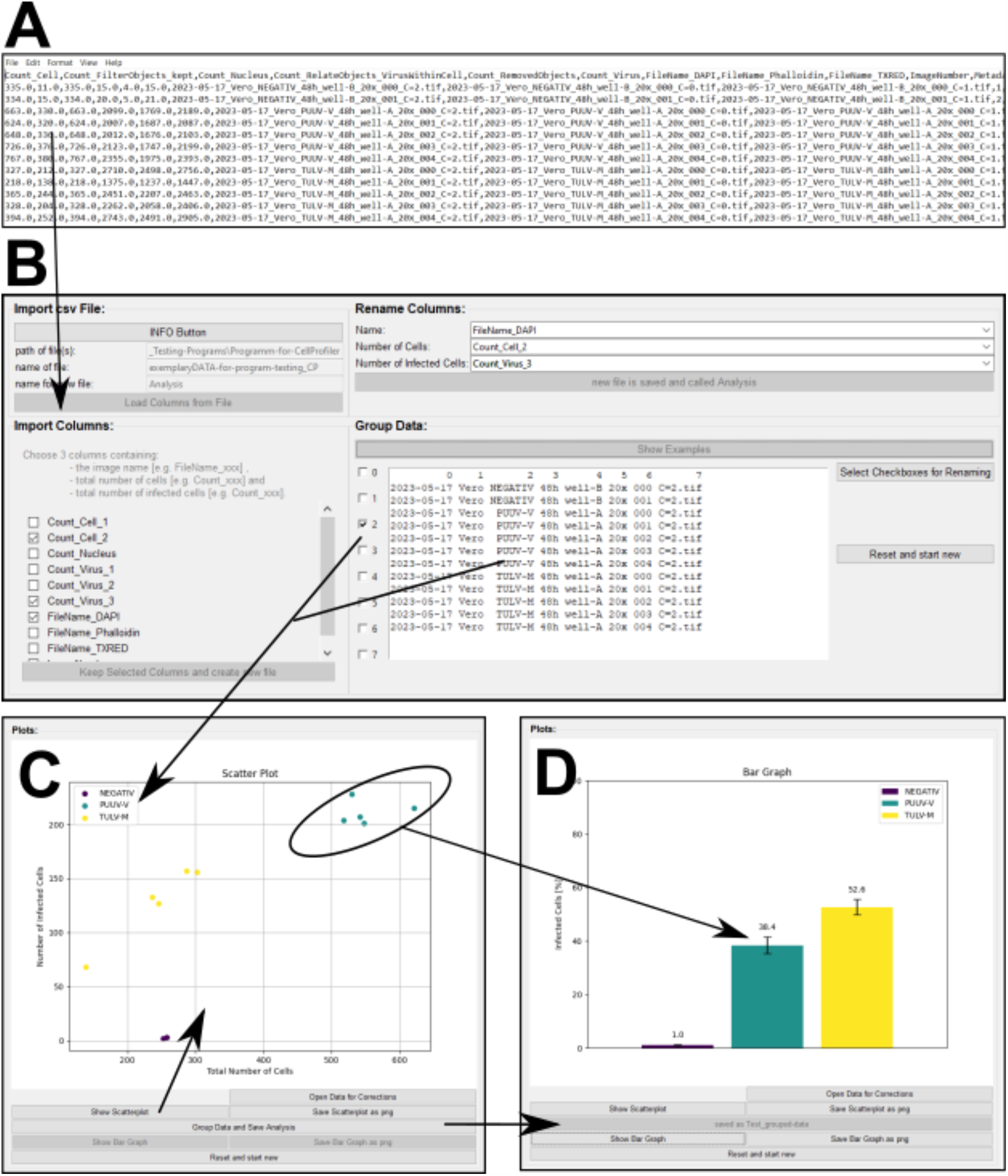
Custom software tool for processing CellProfiler output. (A) shows an exemplary csv file as a result using CellProfiler to quantify PUUV and/or TULV infection after 48 h; (B) Graphical User Interface (GUI) the software that allows processing of csv files. The software tool summarizes and groups important data columns, based on the image name, to get visual output in form of scatterplots (C) or bar graphs (D).

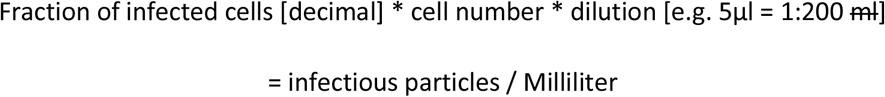

Example: 0.509 * 100,000 * 200 = 10,180,000 or 10.18 * 10^6^ infectious particles per ml

### Automated image analysis allows to extract structural parameters to describe the infection status

During the analysis of our acquired images, we noticed that the size of N protein structures per cell changes over the course of infection. This corresponds to different levels of N protein expression per cell presumably corresponding to different stages of viral replication. We thus wanted to test if such changes can be visualized using our image analysis pipeline. Indeed, in addition to the percentage of infected cells, other parameters such as the size of N protein segments can easily be extracted and displayed over time (**Figure 5 E** and **Supplementary Figure 9**). Both PUUV-N and TULV-N particles are detected after 6 h as small and mostly round-shaped structures. While the number of particles increases over time, also their shape changes, which can be measured using the area of the segmented N protein signal. TULV-N forms, apart from bigger round particles, also long fiber-like structures which can then in detail be investigated using higher magnification (**Figure 5** F, G).

**Figure 5:**
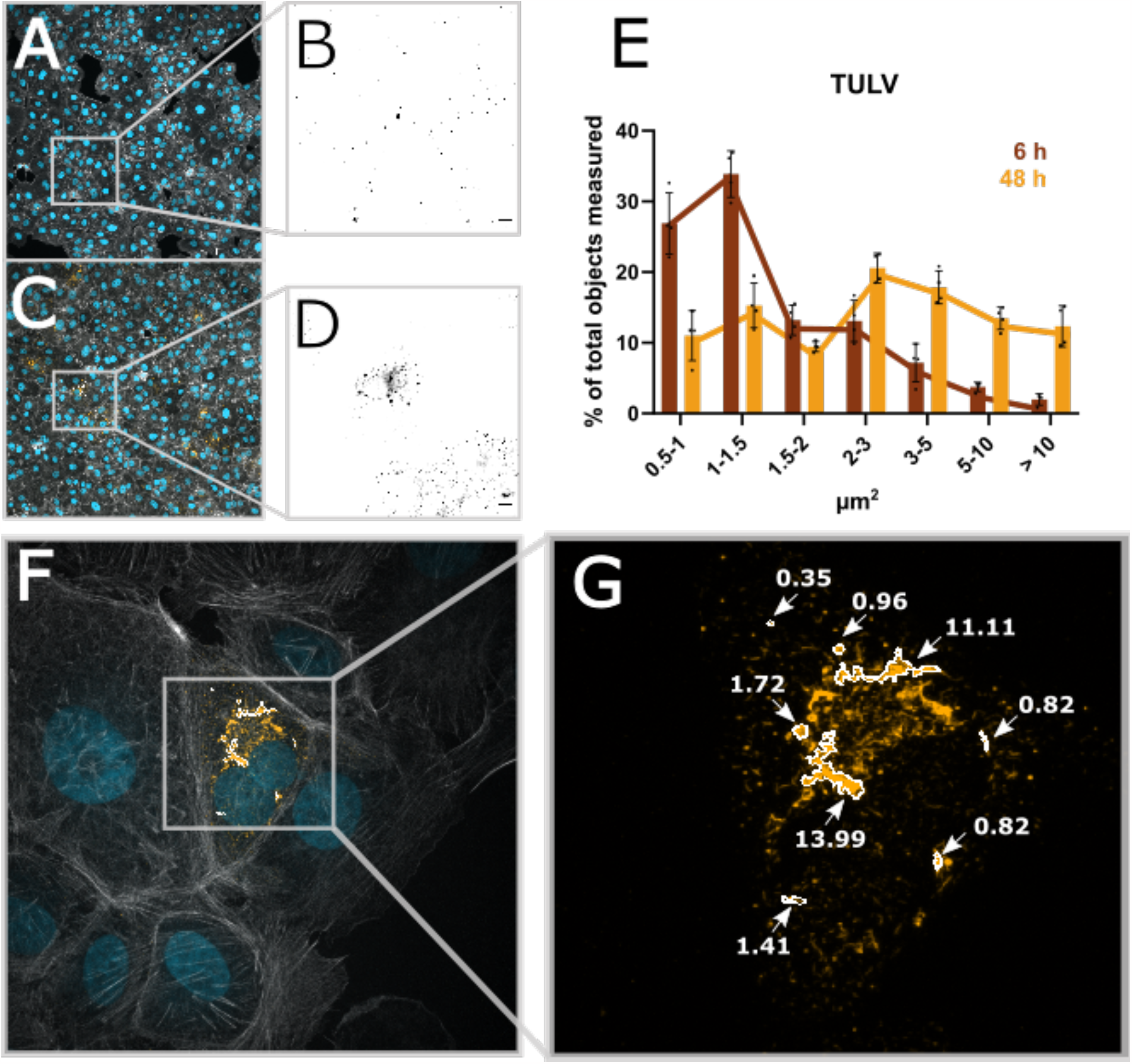
Automated image analysis allows to extract structural parameters. (A) shows a micrograph taken with 20x magnification of Vero E6 cells infected with TULV 6 h post infection. (B) shows the viral N protein; scale bar = 10 μm. (C) and (D) represent the 48 h post infection time point of TULV infection. (E) displays the size of TULV-N segmented particles at either 6 h or 48 h post infection. (F) displays a 100x magnification of 48 hpi TULV infected Vero E6 cells, (G) show exemplary TULV-N structures and the correlating size.

## Conclusion

We developed an optimized workflow to determine the infectious titer of two orthohantavirus strains PUUV and TULV using flow cytometry and fluorescence imaging. Our improved workflow allows to identify infected cells as early as 6 h post infection but we recommend using the 48h time point for robust recognition of infected cells. We provide a detailed protocol of the sample preparation procedure and include new user-friendly and open-source software tools to quickly process results from automated image analysis. In conclusion, our microscopic assay allows fast identification of infected cells with low cell loss and minimized material requirements. Imaging-based quantification furthermore allows to extract structural and positional features of the detected signal which can later be used for the correlation of other phenotypes such as replication status, as we show, or cell cycle for example.

## Supporting information

Supplementary Software

## Acknowledgements

MGLU-2-R (CCLV-RIE 1304) cells were kindly provided by Rainer G. Ulrich, Friedrich-Loeffler-Institute, Greifswald – Insel Riems, Germany. Puumala Virus (PUUV Vranica) was kindly provided by Sandra Essbauer, Bundeswehr Institute of Microbiology, Munich, Germany. Tulavirus (TULV Moravia/Ma5302V) was kindly provided by Rainer G. Ulrich (Friedrich-Loeffler-Institut, Greifswald -Insel Riems, Germany) and Detlev H. Krüger (Institut für Virologie an der Medizinischen Fakultät (Charité) der Humboldt-Universität, Berlin, Germany). C.S. acknowledges support from the Helmholtz Association and the Initiative and Networking Fund for Infection Research Greifswald (Project HANTadapt). Illustrations were generated with BioRender.com.

## Notes

### Competing Interest Statement

The authors have declared no competing interest.

