## Supplementary Figures for "An improved workflow for the quantification of orthohantavirus infection using automated imaging and flow cytometry"

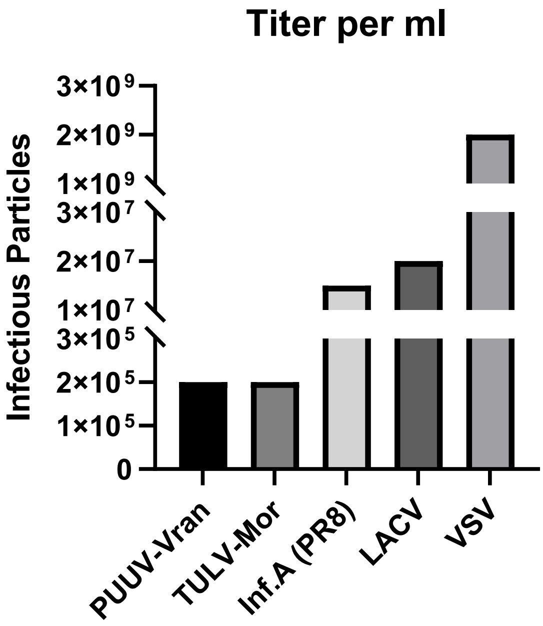


**Supplementary Figure 1: Comparison of the infectious titer of different enveloped RNA virus.** Titers of PUUV-Vranica, TULV-Moravia, Influenza A/Puerto Rico/8/34 (H1N1), LACV and VSV are measure as infectious particles per ml.

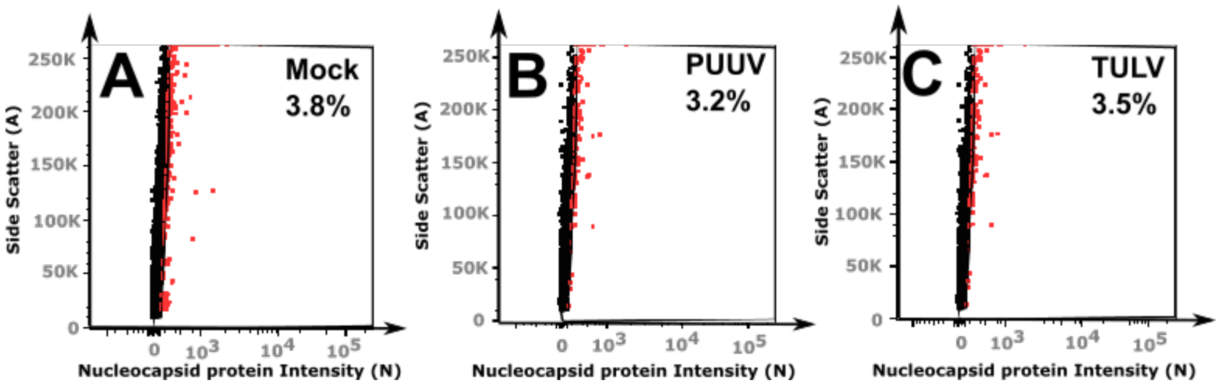


**Supplementary Figure 2: Flow cytometry measurements of Vero E6 cells 6 h post infection with PUUV and TULV.** (A) Mock population and highlighted false positive cells in red. (B) PUUV-Vranica infection and highlighted infected cells in red. (C) TULV-Moravia infection and highlighted infected cells in red. Percentage of infected cells are in the same range as false-positive cells in the Mock control.

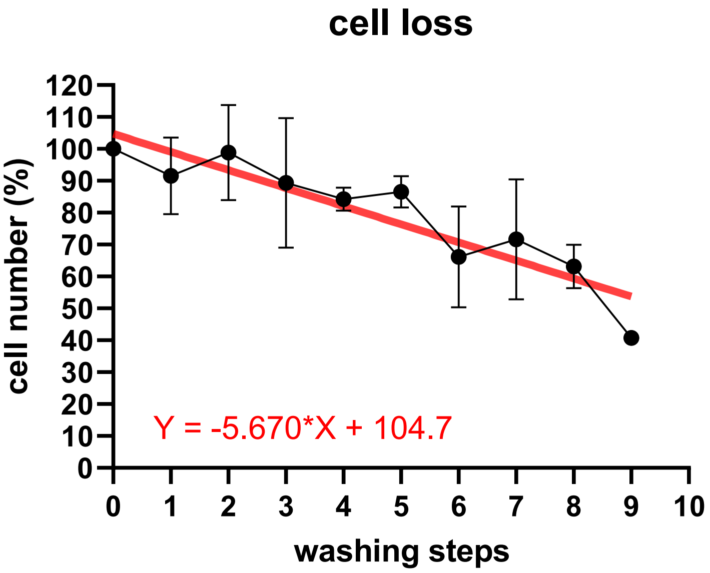


**Supplementary Figure 3: Percentual decrease of Vero E6 cells due to consecutively washing steps in PBS during the flow cytometry staining protocol**.

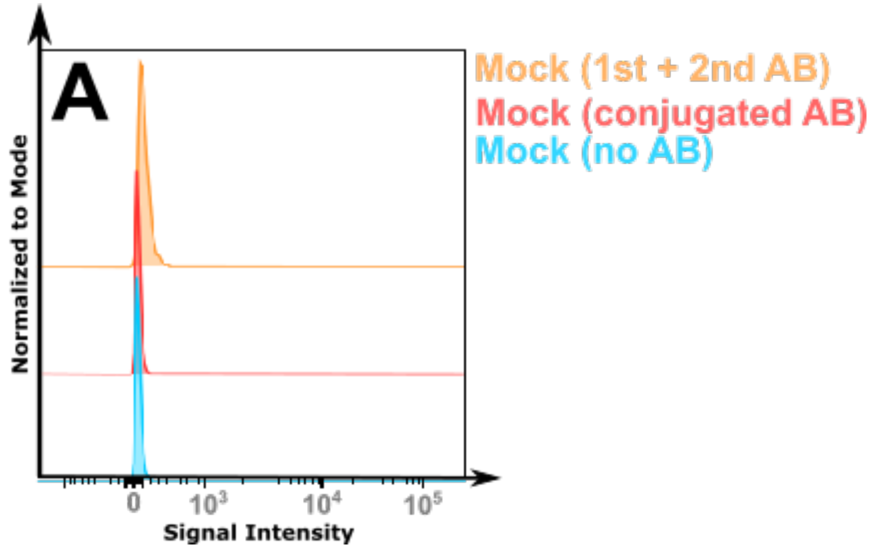


**Supplementary Figure 4: Staining background created by different usage of antibodies (AB)**. Blue: Mock-infected control, without any antibody treatment. In red: Mock-infected cell were stained for 2 h with a conjugated antibody against viral nucleocapsid protein (N). In orange: Mock-infected cells were stained for 1 h with a first antibody against N and afterwards stained for 1 h with a secondary antibody. Cells were washed twice with PBS after each treatment and then measured via a flow cytometer.

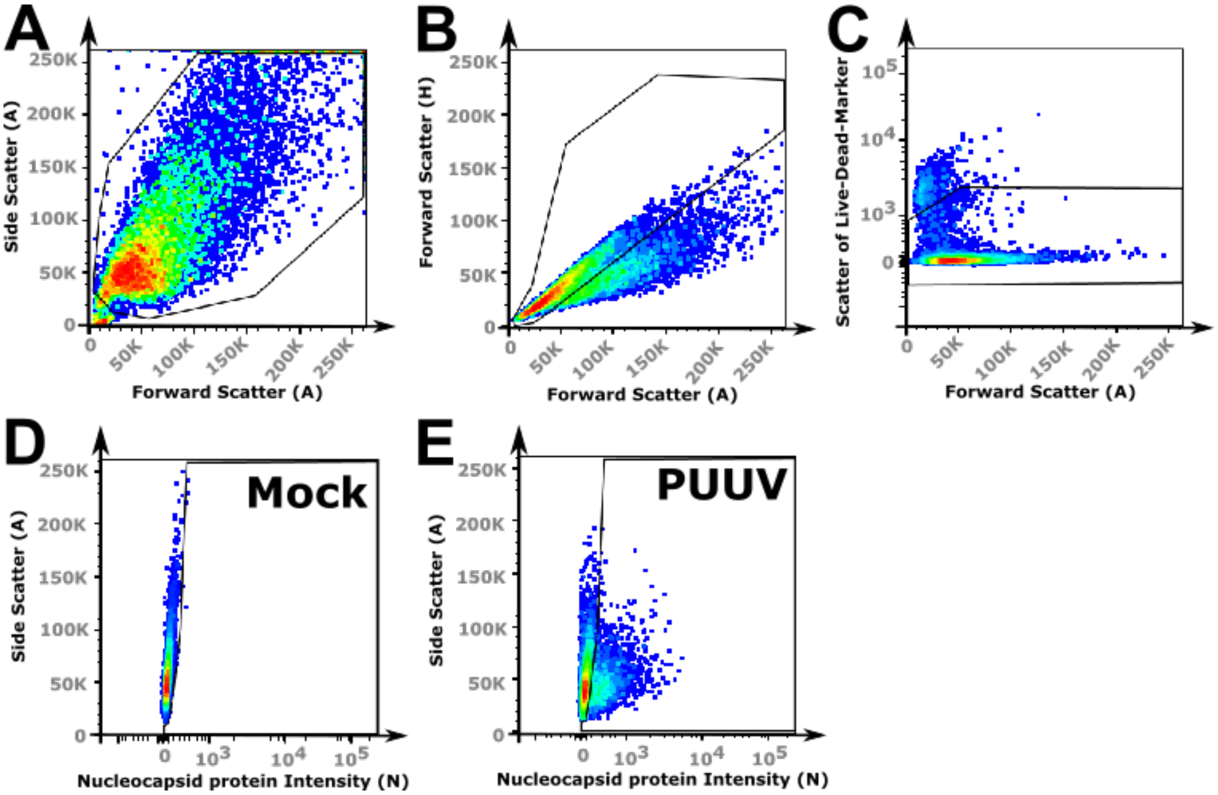


**Supplementary Figure 5: Exemplary gating settings for flow cytometric analysis of infected cells.** (A) Gate settings for Vero E6 cells. (B) Gate setting for single cells, excluding doublets. (C) Gate setting live cells, excluding dead cells. (D) Gate setting to further count infected cells (E).

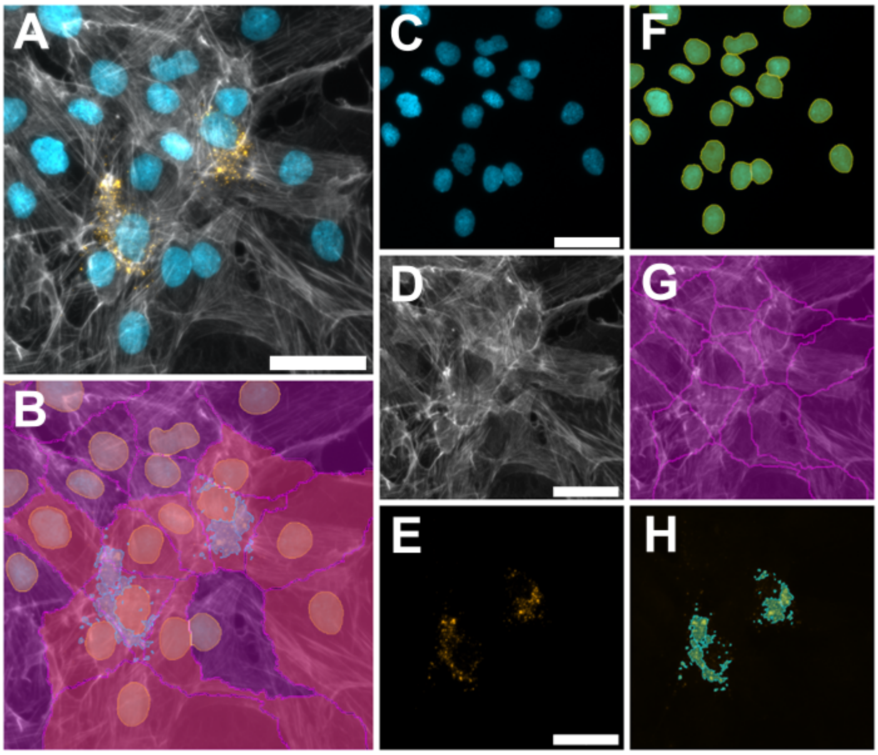


**Supplementary Figure 6: Software recognition of infected cells with a fibroblastic cell-type.** Shown are MGLU-2-R cells infected with PUUV-Vranica. (A-H) shows confronted original images vs. their recognition via software-based masks. (A) shows a merged image of three channels: Hoechst (C) and it´s mask for nuclei recognition (F), Phalloidin staining (D) for recognizing cell bodies or cell boundaries (G), Anti-N staining (E) to reveal infected cells (H). The overlay of F, G and H is shown in B. Scale bars = 50 µm.

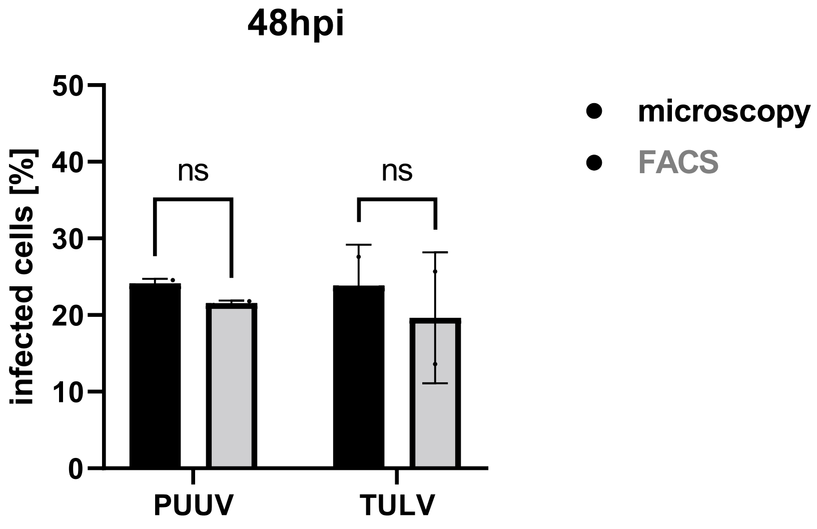


**Supplementary Figure 7: Quantification of PUUV and TULV infectious titer using microscopy or flow cytometry.** A comparison of the results gained with flow cytometry vs. microscopy/imaging shows no significant difference. Significance was analyzed using a unpaired t-test. ****, $p \leq 0.0001$; ***, $p \leq 0.001$; **, $p \leq 0.01$; *, $p \leq 0.05$; ns, $p > 0.05$ .

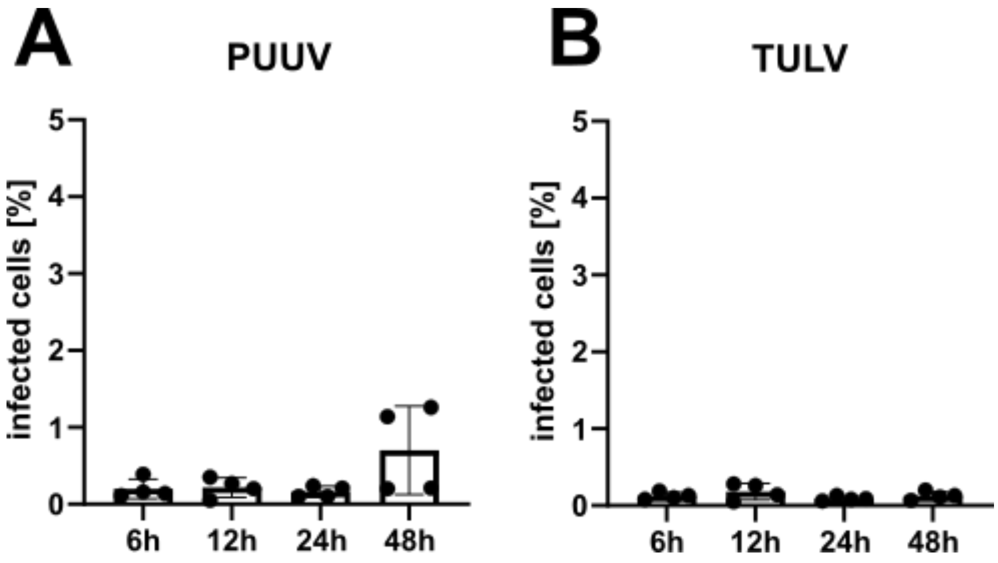


**Supplementary Figure 6:** Vero E6 cells infected with collected supernatant, which was harvested 6h, 12h, 24h and 48h post infection with either PUUV (A) or TULV (B). Reinfection was done for 1 hour, followed by washing and further incubation for 48h. Quantification was done via microscopy.

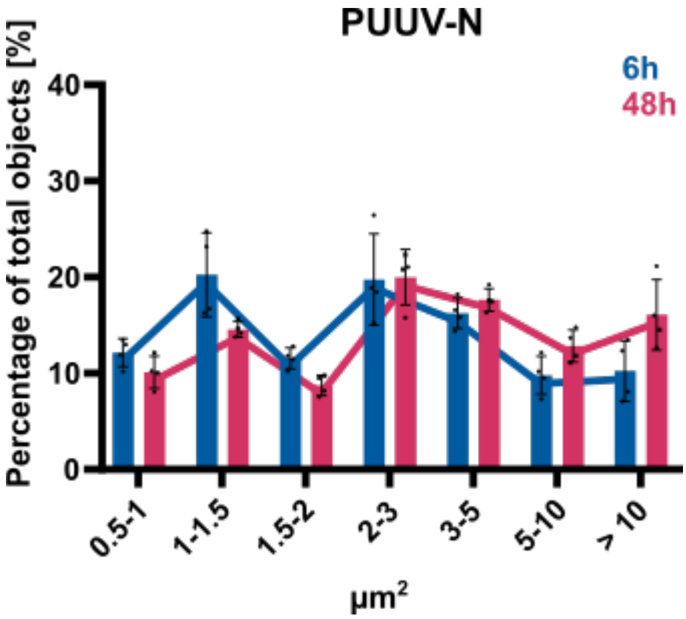


**Supplementary Figure 7:** Size distribution of identified segments of PUUV-N identified in infected cells at different time points (6 h vs. 48 h) in Vero E6 cells.
