## Supplementary Software for "An improved workflow for the quantification of orthohantavirus infection using automated imaging and flow cytometry": Tutorial-CellProfiler_Datalyze.pdf

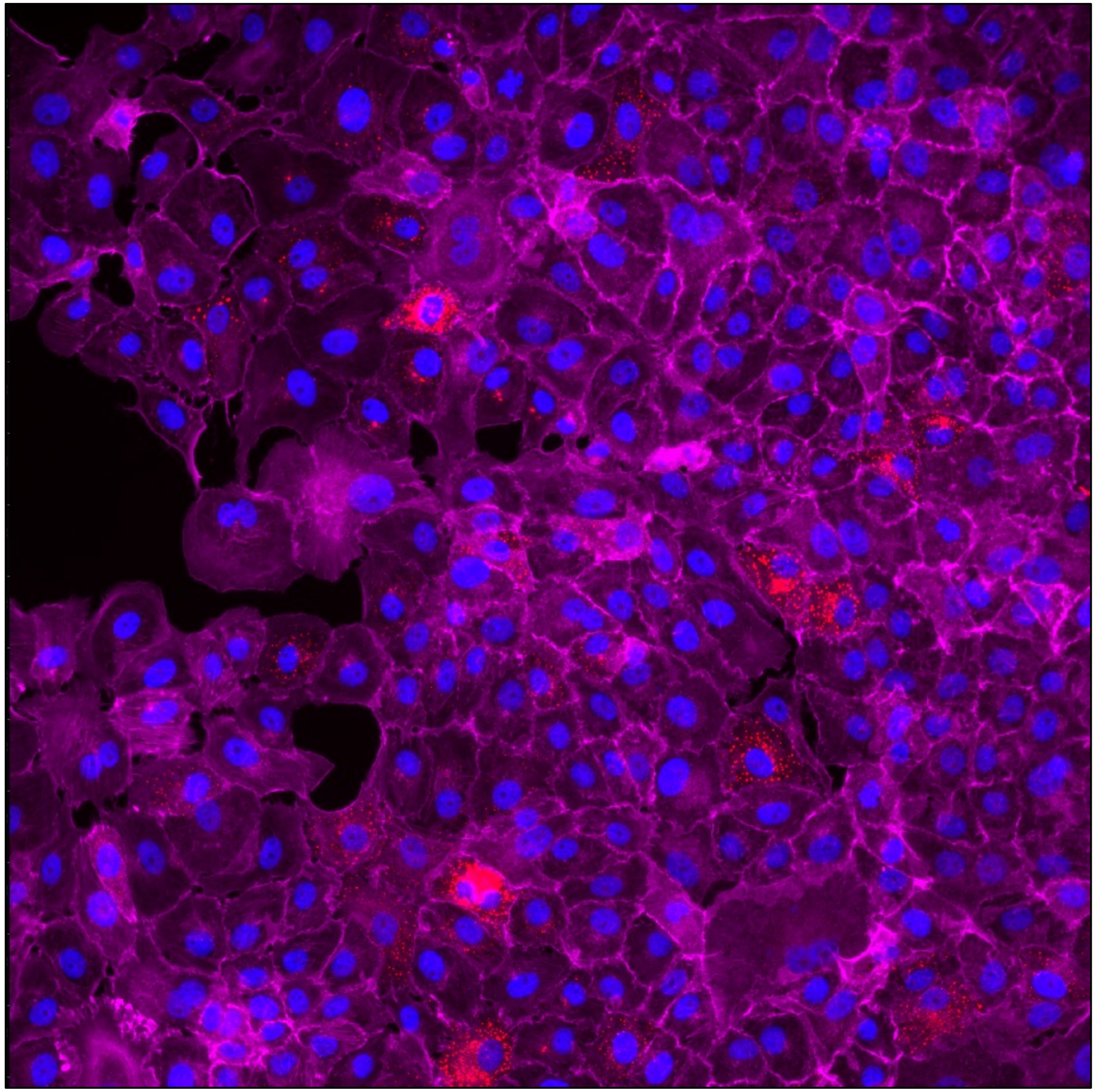

2023-05-17_Vero_TULV-M_48h_well-A_20x_002

### Tutorial for image analysis and virus quantification using CellProfiler

### Necessary staining:

Infobox:
- Tips for using the published python script:
  - Choose your image name by using "_" as a separator
  - e.g. "2023-08-20_VeroE6_PUUV"
  - By recognizing "_" the script will give you the opportunity to get rid of different name parts

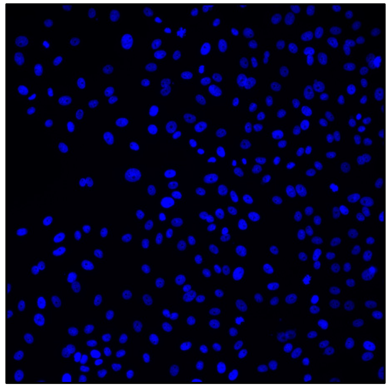

Nucleus staining

e.g Hoechst or DAPI staining

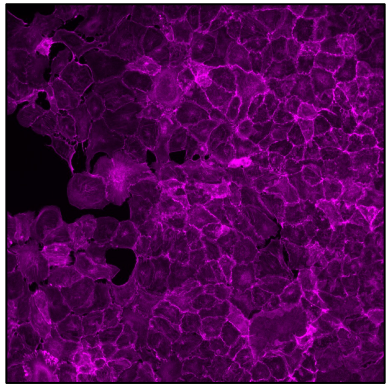

Cell borders

e.g F-Actin staining with Phalloidin

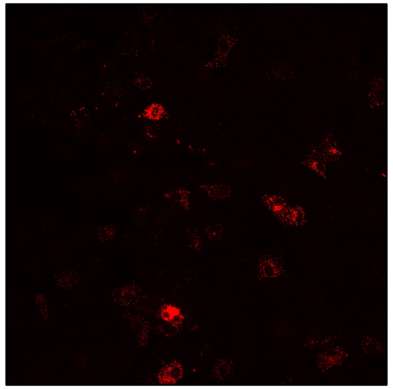

Virus staining

e.g. staining of viral nucleoprotein or glycoprotein

**Empty Pipeline:**

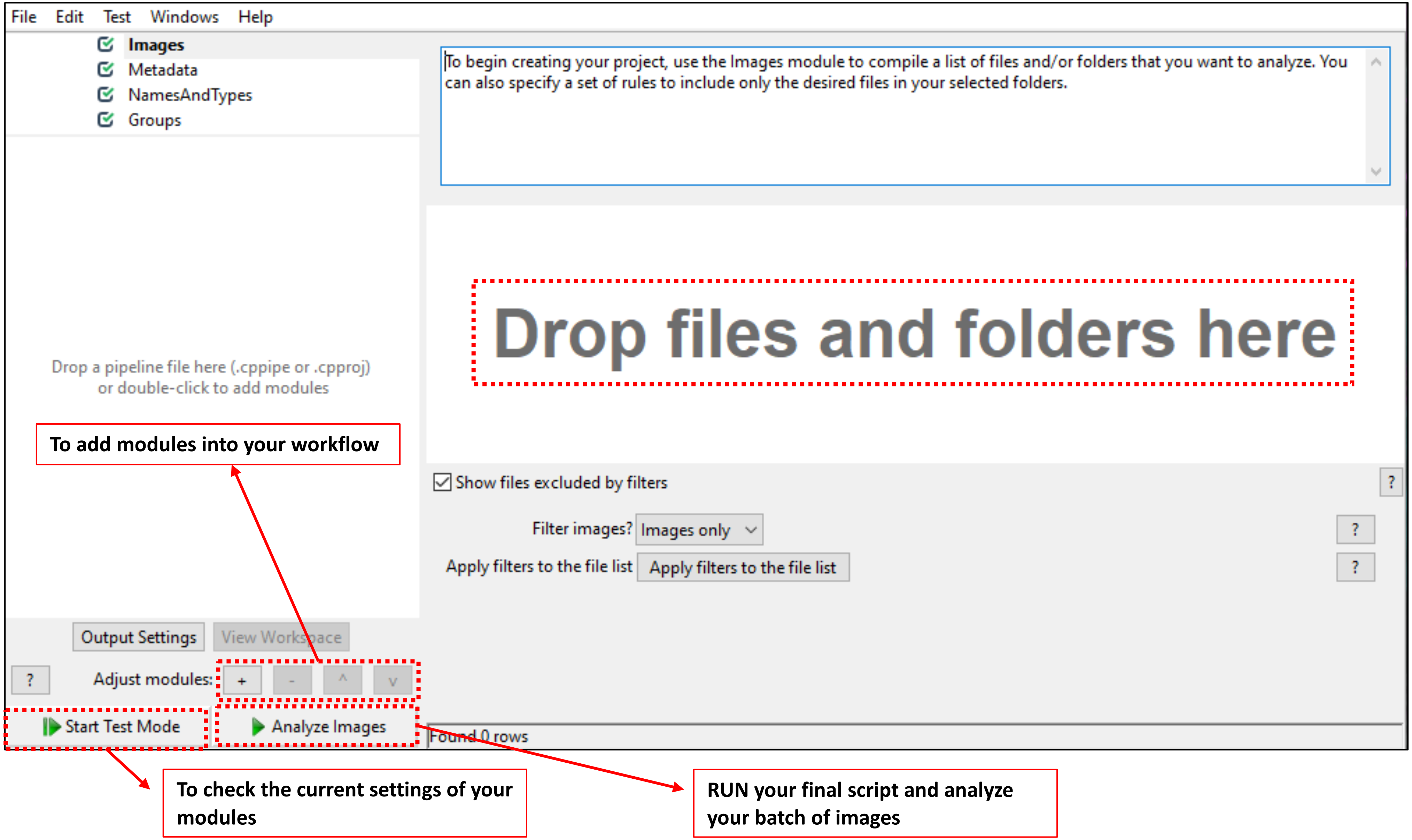

#### < Images >

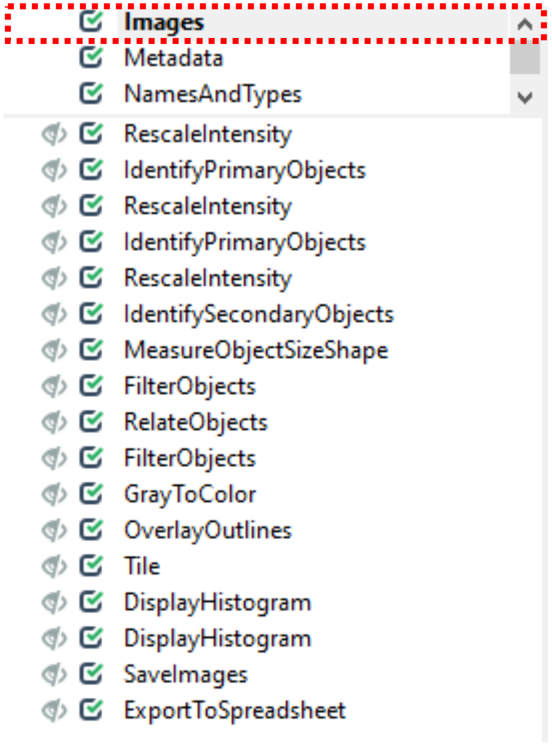


**Folder, which stores images. Drag and Drop images into CellProfiler.**

**One image divided into three channels**

- **Image name:**
  **2023-05-17_VeroE6_NEGATIV_48h_well-B_20x_000**

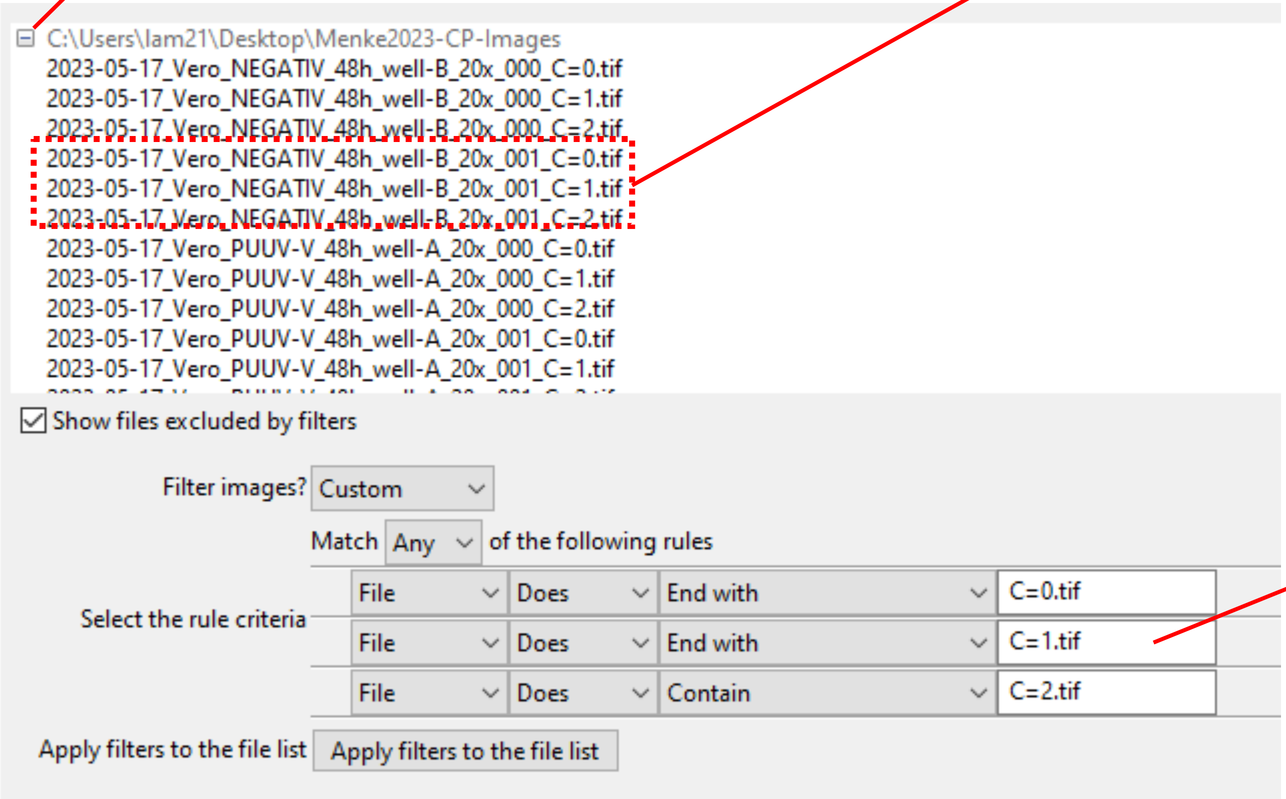


**To exclude other contents**
**(e.g. channels in the selected folder)**

- **Channel-1 e.g. Phalloidin: _C=0**
- **Channel-2 e.g. TXRed: _C=1**
- **Channel-3 e.g. Hoechst: _C=2**

#### < Metadata >

^(?P< xy >.*)_(?P< yz >.*)_...).tif

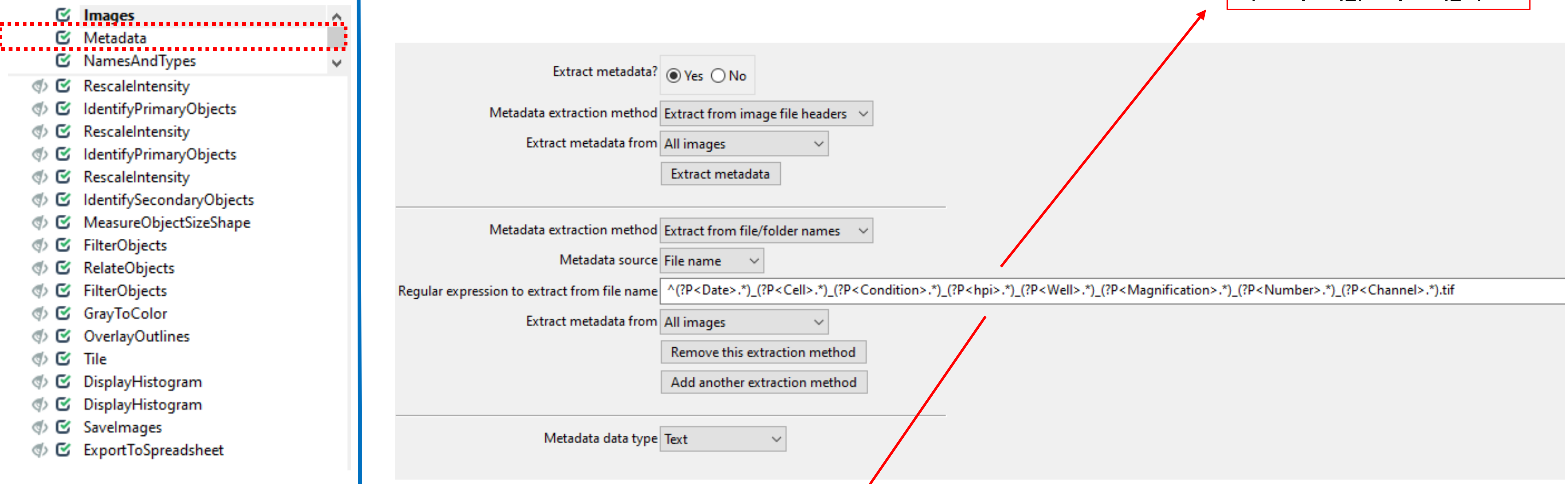


| Update | Path / URL | Series | Frame | Cell | Channel | Condition | Date | FileLocation | Magnification | Number | SizeC | SizeT | SizeZ | Well | hpi |
|---|---|---|---|---|---|---|---|---|---|---|---|---|---|---|---|
| 1 | C:\Users\lam2 | 0 | 0 | Vero | C=0 | NEGATIV | 2023-05-17 | file:///C:/Us...( | 20x | 000 | 1 | 1 | 1 | well-B | 48h |
| 2 | C:\Users\lam2 | 0 | 0 | Vero | C=1 | NEGATIV | 2023-05-17 | file:///C:/Us...( | 20x | 000 | 1 | 1 | 1 | well-B | 48h |
| 3 | C:\Users\lam2 | 0 | 0 | Vero | C=2 | NEGATIV | 2023-05-17 | file:///C:/Us...( | 20x | 000 | 1 | 1 | 1 | well-B | 48h |
| 4 | C:\Users\lam2 | 0 | 0 | Vero | C=0 | NEGATIV | 2023-05-17 | file:///C:/Us...( | 20x | 001 | 1 | 1 | 1 | well-B | 48h |
| 5 | C:\Users\lam2 | 0 | 0 | Vero | C=1 | NEGATIV | 2023-05-17 | file:///C:/Us...( | 20x | 001 | 1 | 1 | 1 | well-B | 48h |
| 6 | C:\Users\lam2 | 0 | 0 | Vero | C=2 | NEGATIV | 2023-05-17 | file:///C:/Us...( | 20x | 001 | 1 | 1 | 1 | well-B | 48h |
| 7 | C:\Users\lam2 | 0 | 0 | Vero | C=0 | PUUV-V | 2023-05-17 | file:///C:/Us...( | 20x | 000 | 1 | 1 | 1 | well-A | 48h |
| 8 | C:\Users\lam2 | 0 | 0 | Vero | C=1 | PUUV-V | 2023-05-17 | file:///C:/Us...( | 20x | 000 | 1 | 1 | 1 | well-A | 48h |
| 9 | C:\Users\lam2 | 0 | 0 | Vero | C=2 | PUUV-V | 2023-05-17 | file:///C:/Us...( | 20x | 000 | 1 | 1 | 1 | well-A | 48h |

### < NamesAndTypes >

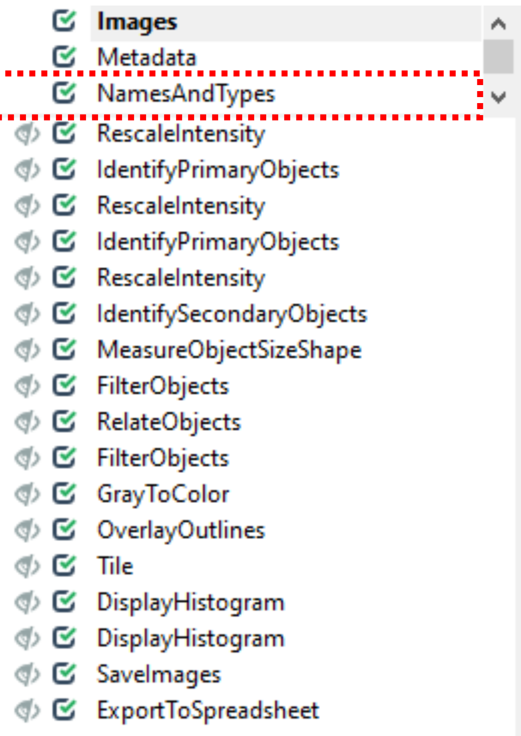


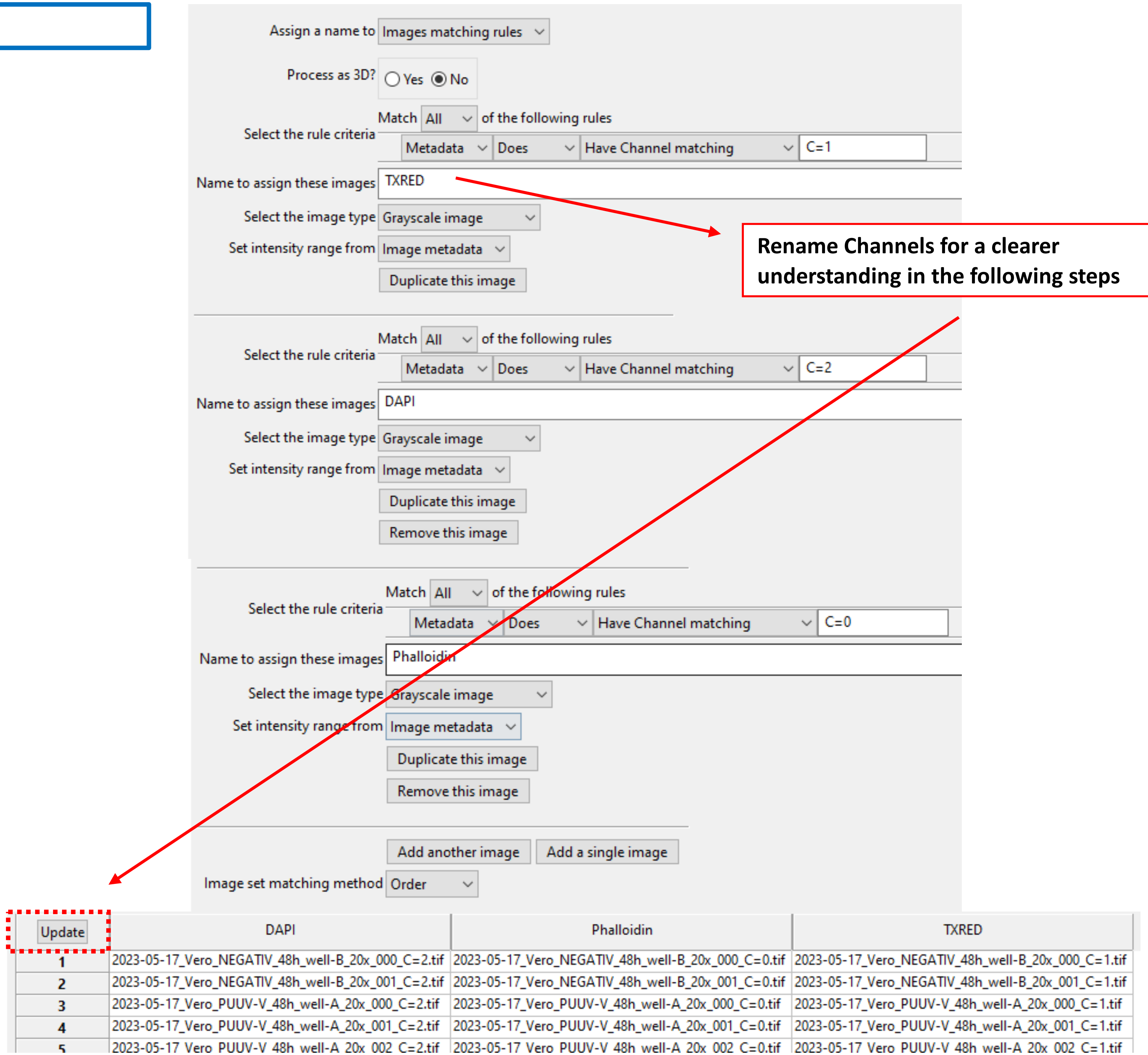


**Rename Channels for a clearer understanding in the following steps**

#### < Groups >

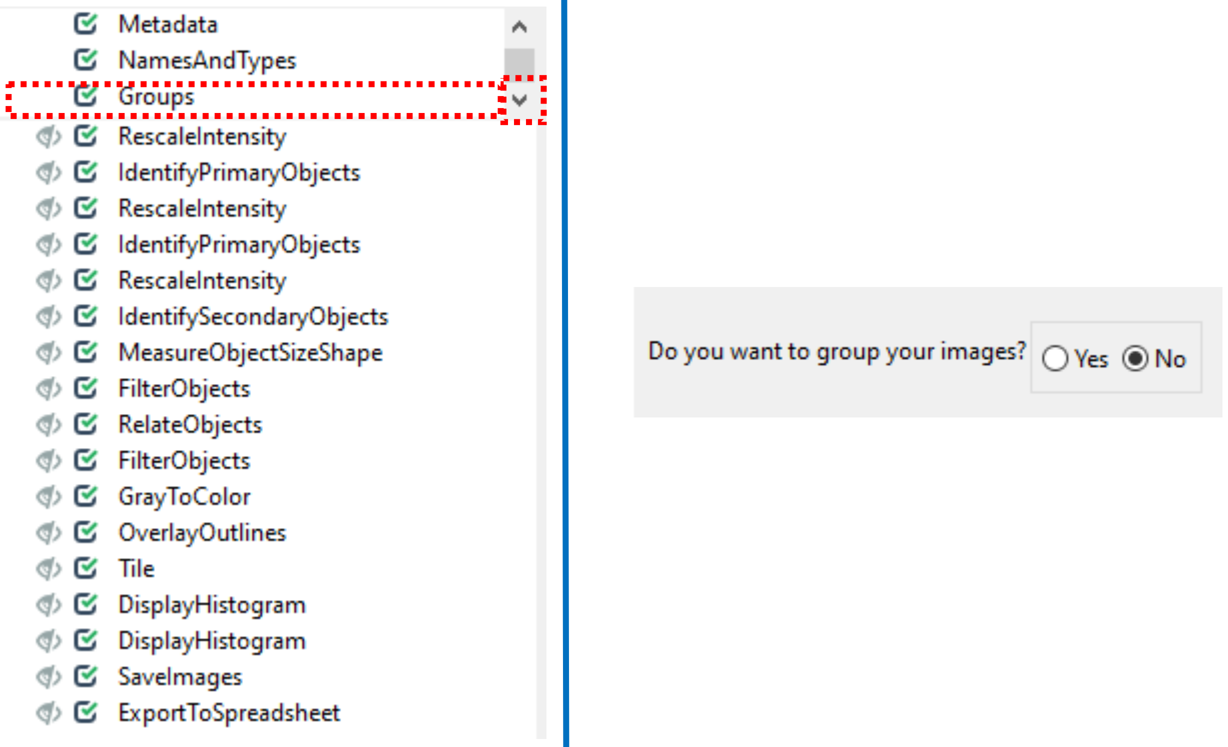
Metadata
NamesAndTypes
Groups
RescaleIntensity
IdentifyPrimaryObjects
RescaleIntensity
IdentifyPrimaryObjects
RescaleIntensity
IdentifySecondaryObjects
MeasureObjectSizeShape
FilterObjects
RelateObjects
FilterObjects
GrayToColor
OverlayOutlines
Tile
DisplayHistogram
DisplayHistogram
SaveImages
ExportToSpreadsheet
Do you want to group your images?
Yes
No

#### < RescaleIntensity >

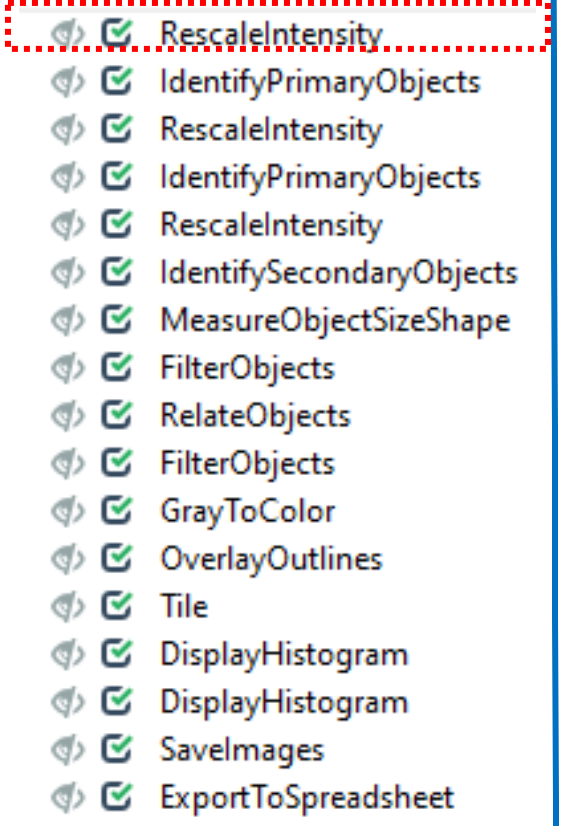


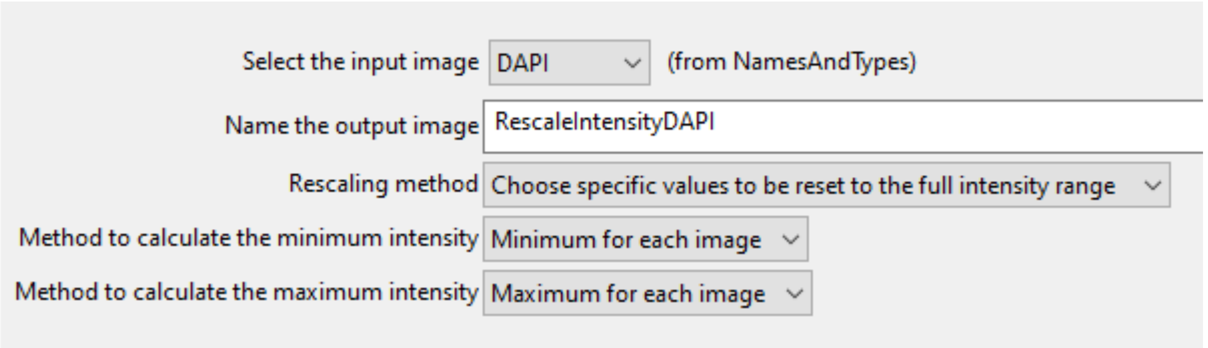


Necessary in case of less signal intensity of e.g. Hoechst staining

Settings can be "for each image" individually: -> it is important to get the most intense signal to get good masks per image. But we do not compare nuclei between different pictures, therefore it do not have to be a global value.

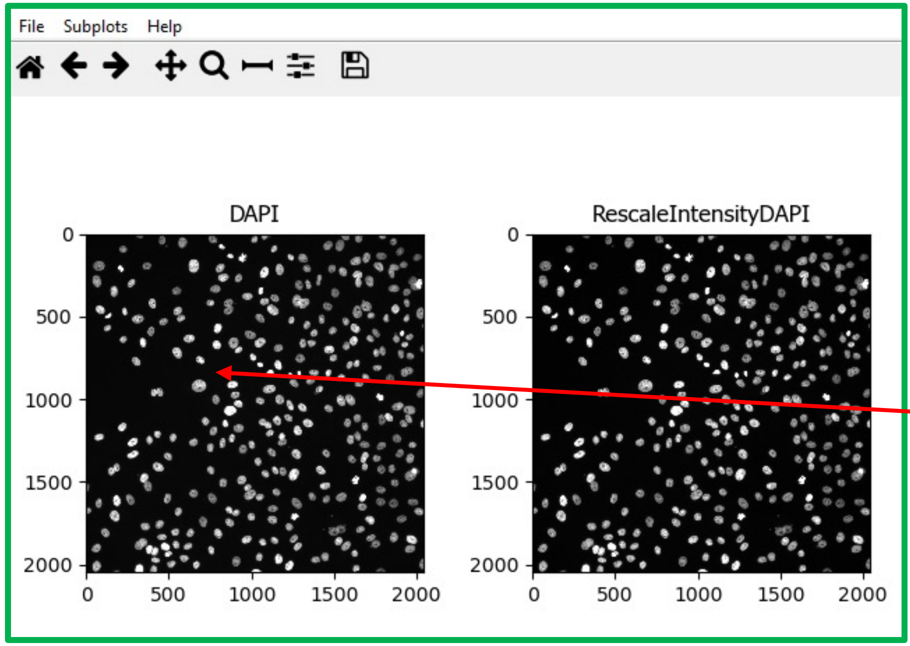


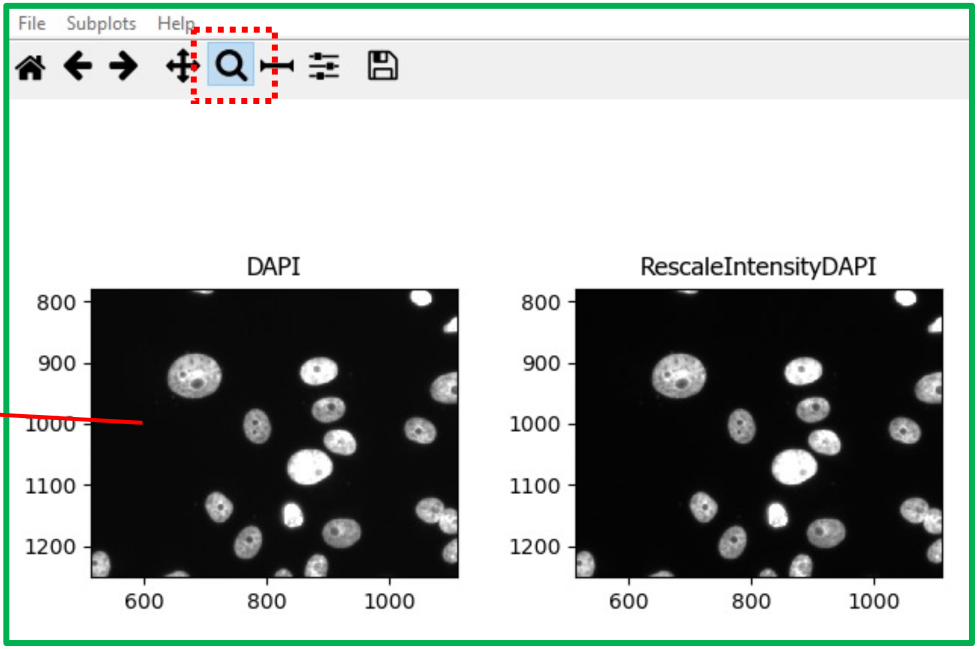

### < IdentifyPrimaryObjects >

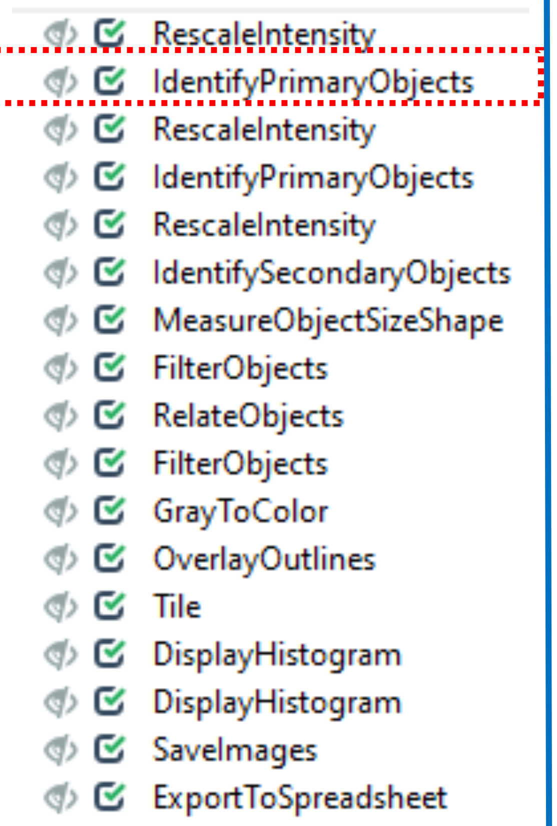


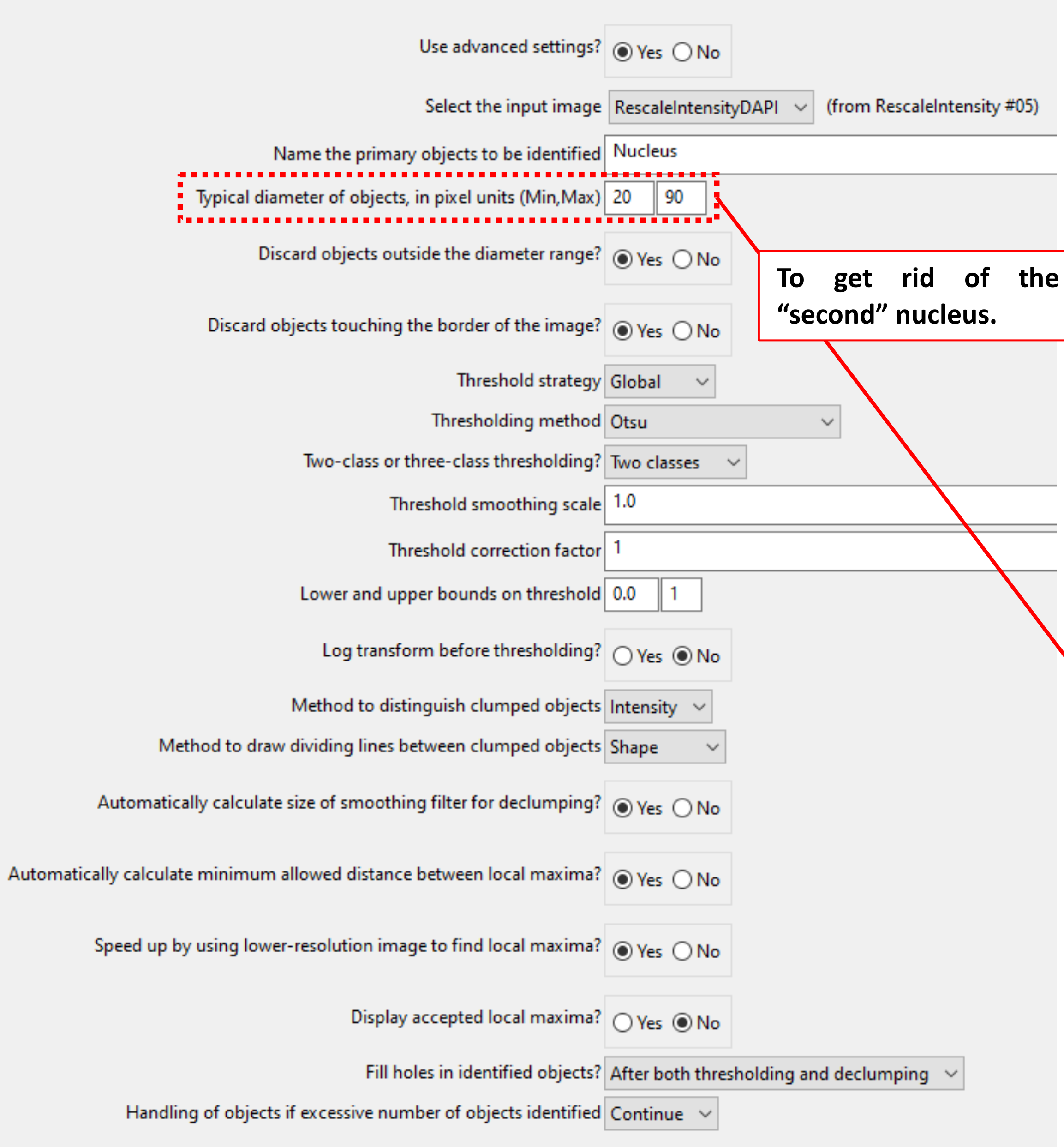


To get rid of the "second" nucleus.

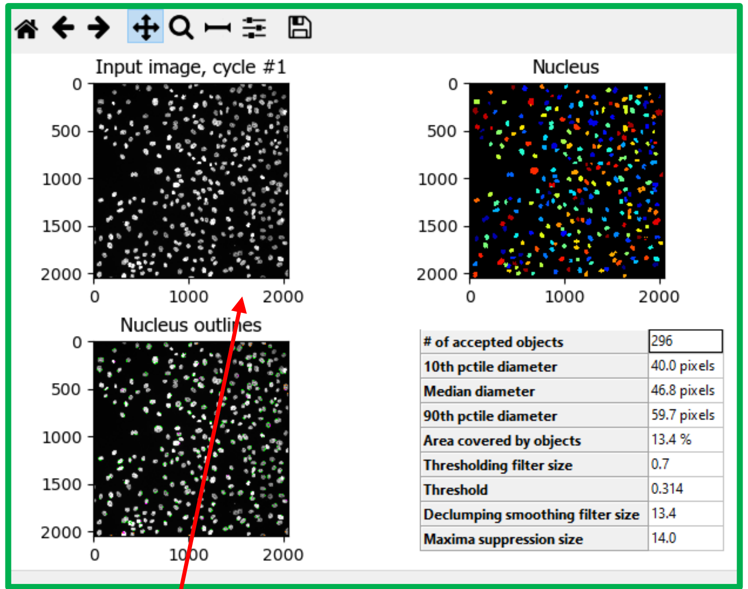


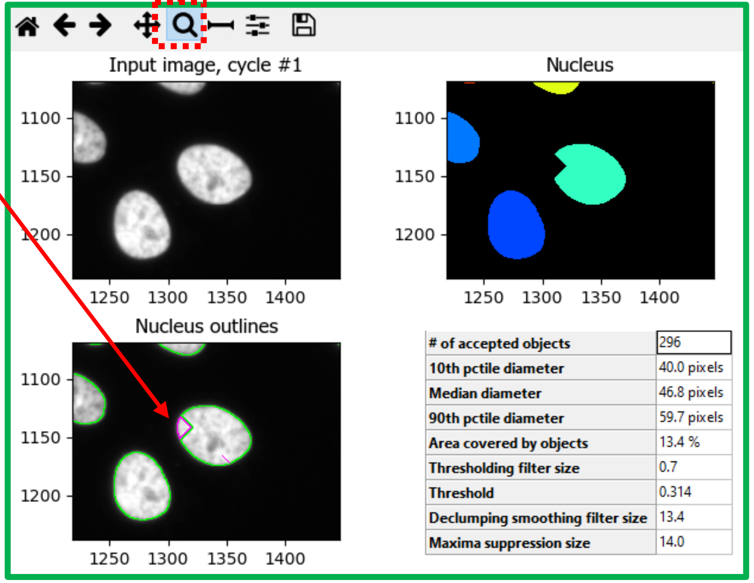


Zoom in to see how efficient and successful the software recognizes the nuclei.
Need to be adapted for each new analysis / new images.

#### < RescaleIntensity >

**! Critical step !**

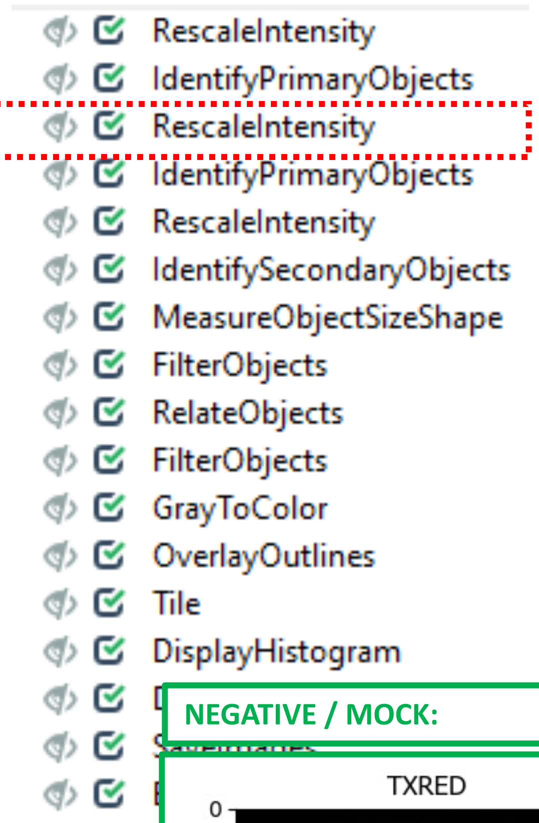


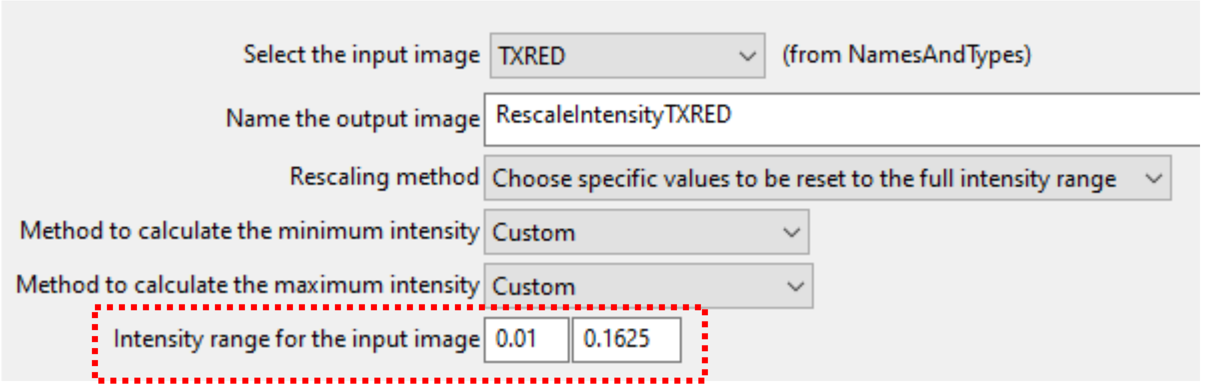


**A global Intensity range is important to measure all images with the same conditions.**

**We recommend to use a software tool (e.g. Fiji) to check within the original image how much intensity for the viral N still shows a real virus staining and at which point CellProfiler would also count background noise**

**NEGATIVE / MOCK:**

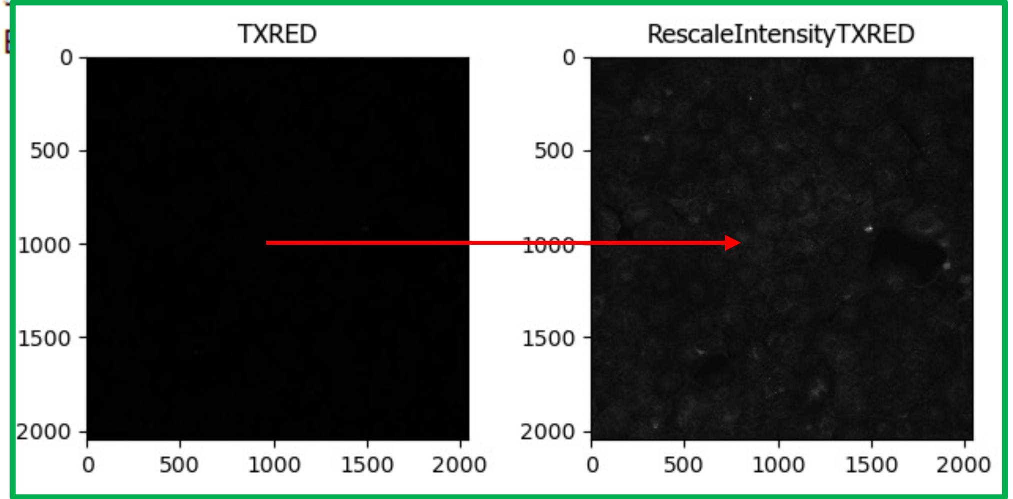


**Infected Cells:**

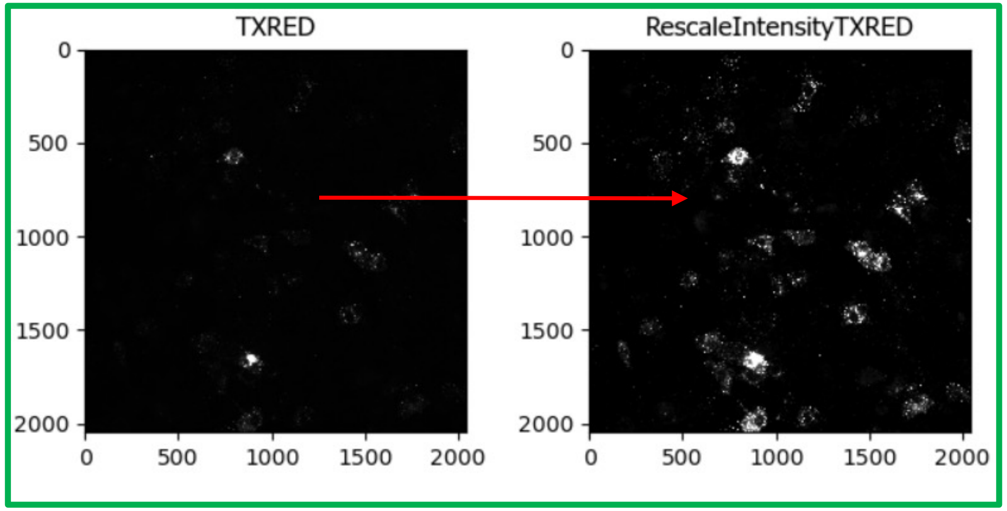

### < IdentifyPrimaryObjects >

- RescaleIntensity
- IdentifyPrimaryObjects
- RescaleIntensity
- IdentifyPrimaryObjects
- RescaleIntensity
- IdentifySecondaryObjects
- MeasureObjectSizeShape
- FilterObjects
- RelateObjects
- FilterObjects
- GrayToColor
- OverlayOutlines
- Tile
- DisplayHistogram
- DisplayHistogram
- SaveImages
- ExportToSpreadsheet

| | |
|---|---|
| Use advanced settings? | (•) Yes ( ) No |
| Select the input image | RescaleIntensityTXRED (from RescaleIntensity #07) |
| Name the primary objects to be identified | Virus_1 |
| Typical diameter of objects, in pixel units (Min,Max) | 10 / 500 |
| Discard objects outside the diameter range? | ( ) Yes (•) No |
| Discard objects touching the border of the image? | ( ) Yes (•) No |
| Threshold strategy | Global |
| Thresholding method | Minimum Cross-Entropy |
| Threshold smoothing scale | 1.3488 |
| Threshold correction factor | 1.0 |
| Lower and upper bounds on threshold | 0.3 / 1.0 |
| Log transform before thresholding? | ( ) Yes (•) No |
| Method to distinguish clumped objects | Intensity |
| Method to draw dividing lines between clumped objects | Intensity |
| Automatically calculate size of smoothing filter for declumping? | (•) Yes ( ) No |
| Automatically calculate minimum allowed distance between local maxima? | (•) Yes ( ) No |
| Speed up by using lower-resolution image to find local maxima? | (•) Yes ( ) No |
| Display accepted local maxima? | ( ) Yes (•) No |
| Fill holes in identified objects? | After both thresholding and declumping |
| Handling of objects if excessive number of objects identified | Continue |

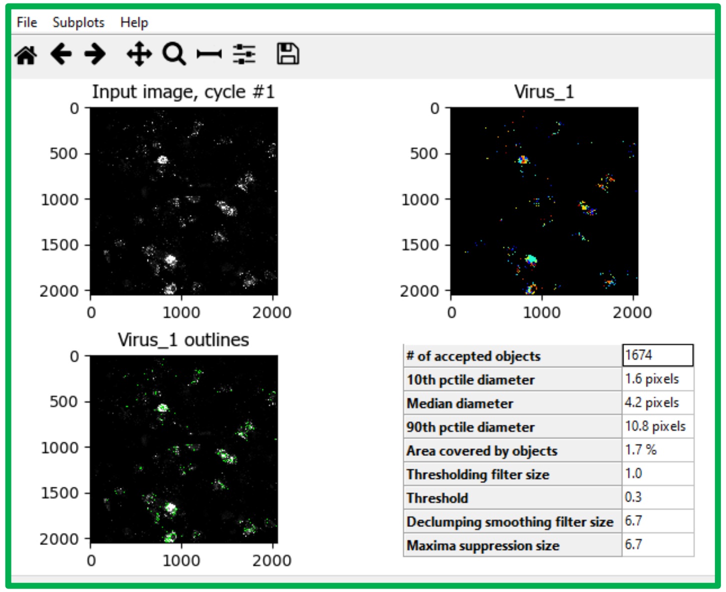


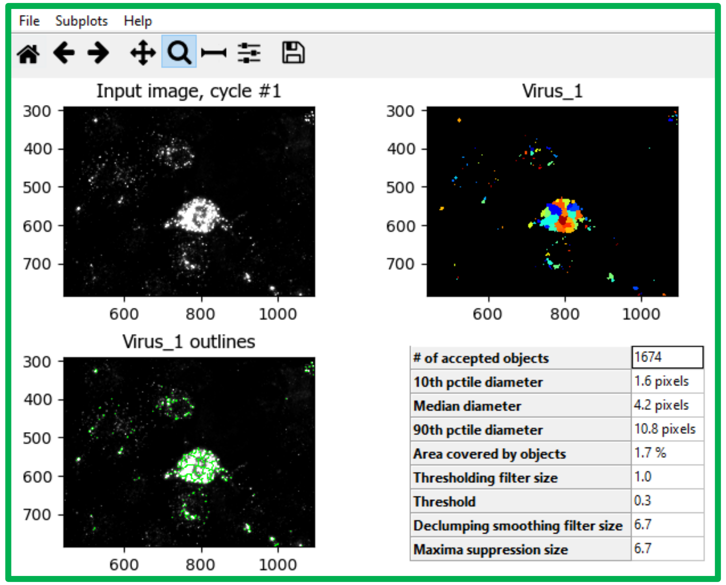

**< RescaleIntensity >**

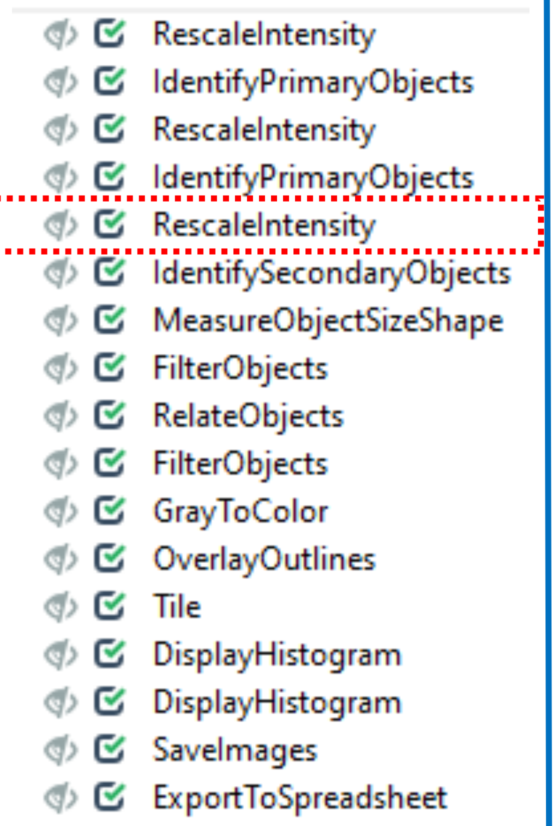


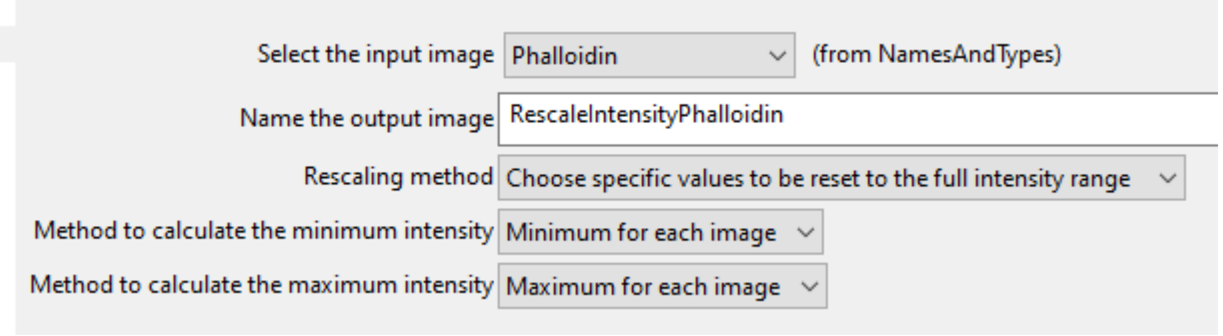


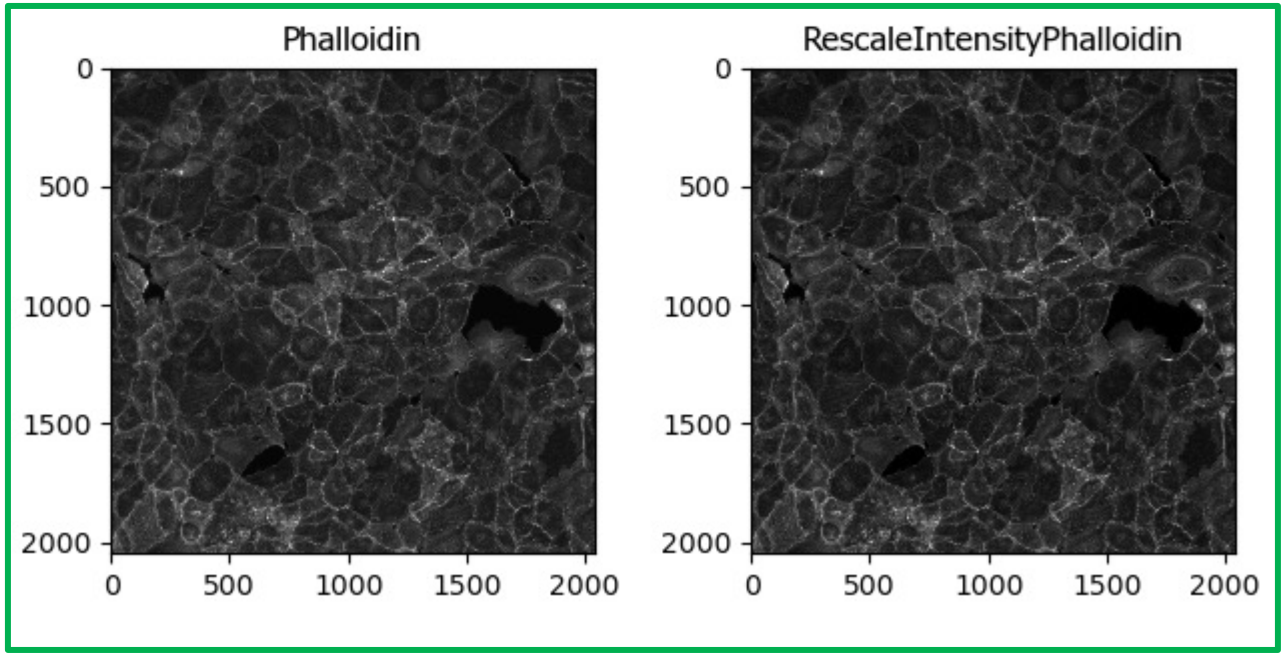

#### < IndentifySecondaryObjects >

- RescaleIntensity
- IdentifyPrimaryObjects
- RescaleIntensity
- IdentifyPrimaryObjects
- RescaleIntensity
- **IdentifySecondaryObjects**
- MeasureObjectSizeShape
- FilterObjects
- RelateObjects
- FilterObjects
- GrayToColor
- OverlayOutlines
- Tile
- DisplayHistogram
- DisplayHistogram
- SaveImages
- ExportToSpreadsheet

| | | |
|---|---|---|
| Select the input image | RescaleIntensityPhalloidin | (from RescaleIntensity #09) |
| Select the input objects | Nucleus | (from IdentifyPrimaryObjects #06) |
| Name the objects to be identified | Cell_1 | |
| Select the method to identify the secondary objects | Propagation | |
| Threshold strategy | Global | |
| Thresholding method | Minimum Cross-Entropy | |
| Threshold smoothing scale | 0 | |
| Threshold correction factor | 1.0 | |
| Lower and upper bounds on threshold | 0.0 / 0.003 | |
| Log transform before thresholding? | ○ Yes ◉ No | |
| Regularization factor | 0.5 | |
| Fill holes in identified objects? | ◉ Yes ○ No | |
| Discard secondary objects touching the border of the image? | ○ Yes ◉ No | |

**Check if nuclei and cell border are properly detected. Need to be adapted for each new analysis / new images**

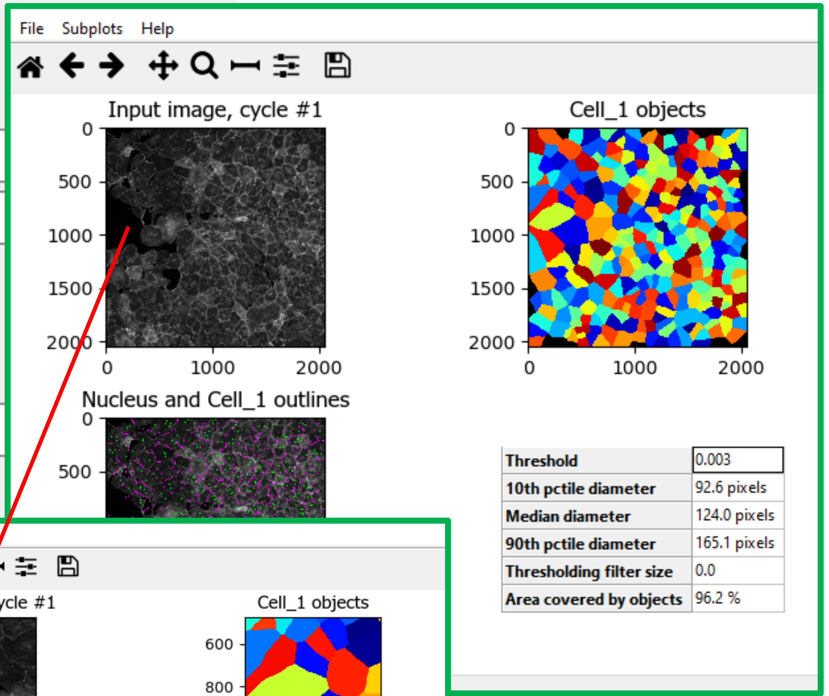


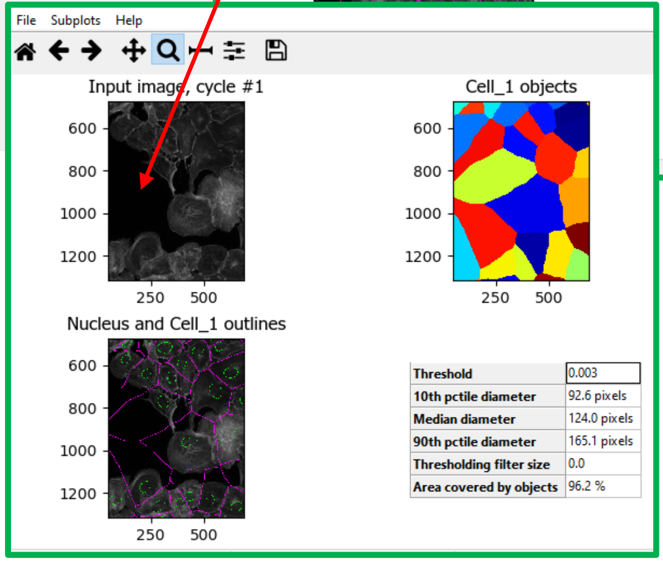


**In next step the software need to exclude big "cells", which are empty spaces in the original image**

< MeasureObjectSizeShape >

- RescaleIntensity
- IdentifyPrimaryObjects
- RescaleIntensity
- IdentifyPrimaryObjects
- RescaleIntensity
- IdentifySecondaryObjects
- MeasureObjectSizeShape
- FilterObjects
- RelateObjects
- FilterObjects
- GrayToColor
- OverlayOutlines
- Tile
- DisplayHistogram
- DisplayHistogram
- SaveImages
- ExportToSpreadsheet

Select object sets to measure

- [x] Cell_1 (from IdentifySecondaryObjects #10)
- [ ] Nucleus (from IdentifyPrimaryObjects #06)
- [ ] Virus_1 (from IdentifyPrimaryObjects #08)

Calculate the Zernike features? ○ Yes ◉ No

Calculate the advanced features? ○ Yes ◉ No

| Object | Feature | Mean | Median | STD |
|---|---|---|---|---|
| Cell_1 | Area | 13600.99 | 12059.00 | 8082.43 |
| Cell_1 | Perimeter | 484.63 | 475.82 | 131.53 |
| Cell_1 | MajorAxisLength | 161.27 | 156.06 | 48.65 |
| Cell_1 | MinorAxisLength | 108.28 | 105.02 | 29.24 |
| Cell_1 | Eccentricity | 0.70 | 0.71 | 0.14 |
| Cell_1 | Orientation | 0.71 | 6.42 | 50.87 |
| Cell_1 | Center_X | 1155.72 | 1181.43 | 548.12 |
| Cell_1 | Center_Y | 1041.83 | 1051.01 | 582.68 |
| Cell_1 | BoundingBoxArea | 22270.39 | 18879.00 | 14519.33 |
| Cell_1 | BoundingBoxMinimum_X | 1083.28 | 1115.50 | 557.05 |
| Cell_1 | BoundingBoxMaximum_X | 1228.77 | 1248.00 | 538.91 |
| Cell_1 | BoundingBoxMinimum_Y | 969.61 | 967.00 | 584.76 |
| Cell_1 | BoundingBoxMaximum_Y | 1114.92 | 1111.50 | 580.06 |
| Cell_1 | FormFactor | 0.69 | 0.69 | 0.06 |
| Cell_1 | Extent | 0.62 | 0.61 | 0.08 |
| Cell_1 | Solidity | 0.95 | 0.96 | 0.02 |
| Cell_1 | Compactness | 1.47 | 1.44 | 0.16 |
| Cell_1 | EulerNumber | 1.00 | 1.00 | 0.00 |
| Cell_1 | MaximumRadius | 49.59 | 48.00 | 13.64 |
| Cell_1 | MeanRadius | 18.01 | 17.48 | 4.74 |
| Cell_1 | MedianRadius | 16.12 | 15.81 | 4.20 |
| Cell_1 | ConvexArea | 14325.33 | 12728.00 | 8575.24 |
| Cell_1 | MinFeretDiameter | 110.72 | 107.25 | 30.72 |
| Cell_1 | MaxFeretDiameter | 172.92 | 167.64 | 50.91 |
| Cell_1 | EquivalentDiameter | 127.38 | 123.91 | 33.04 |

**Orientation for setting in the next module.**

#### < FilterObjects >

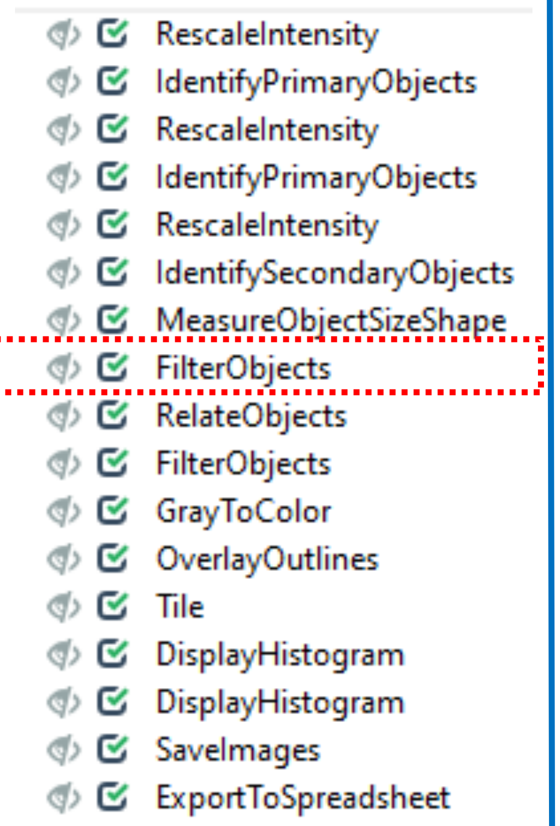


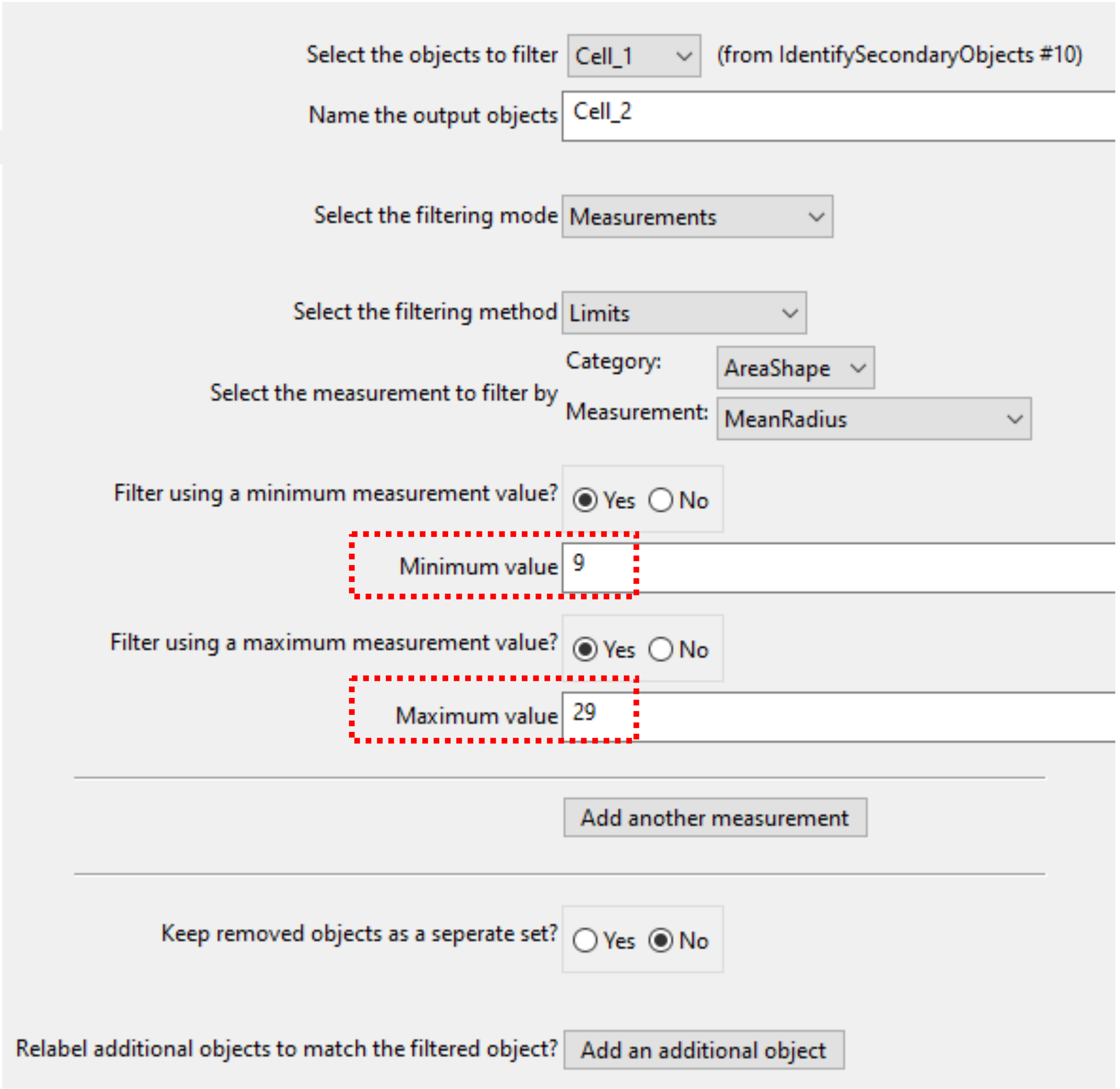


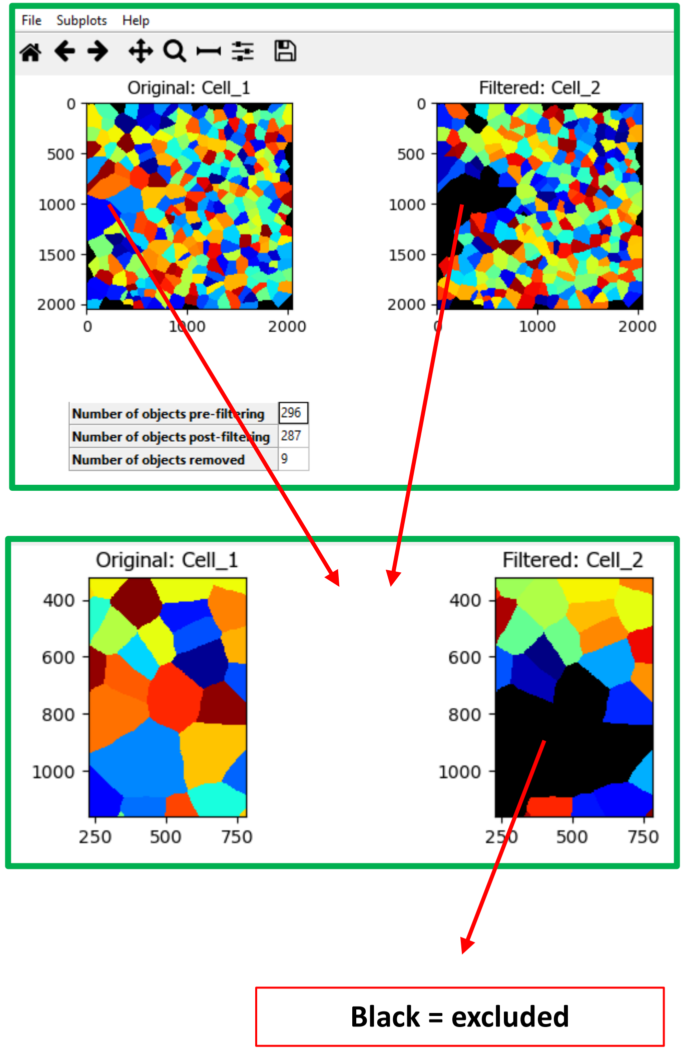

#### < RelateObjects >

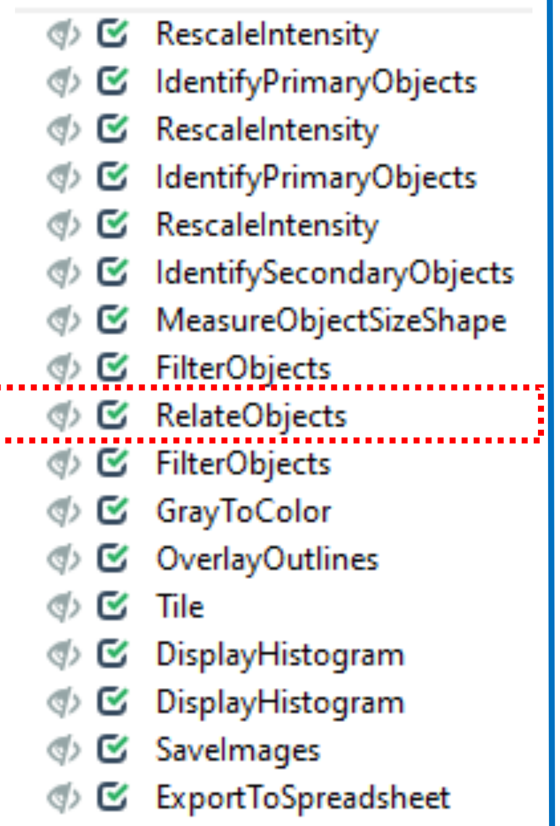


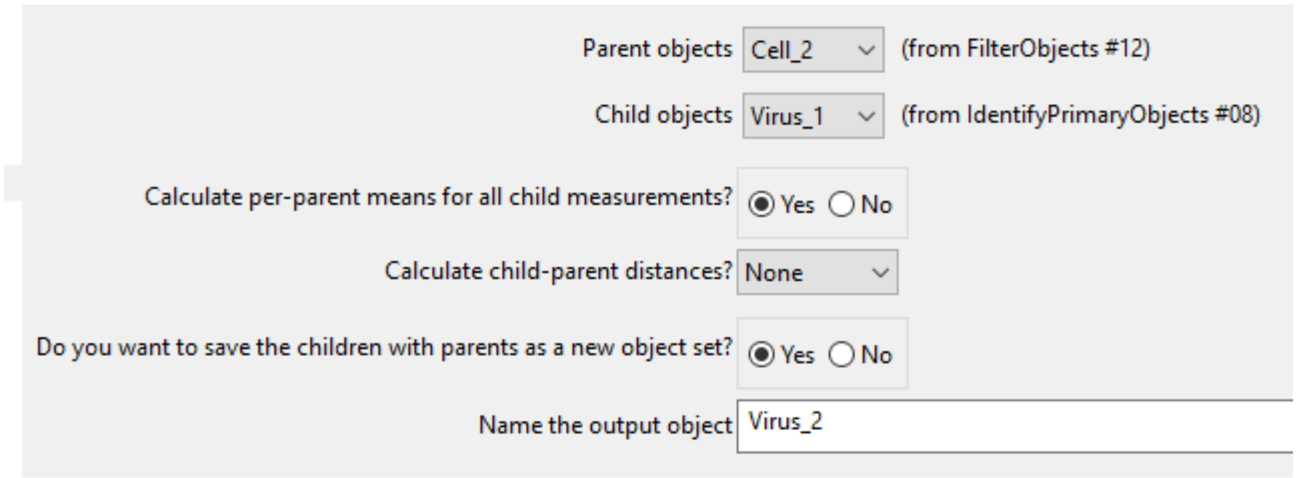


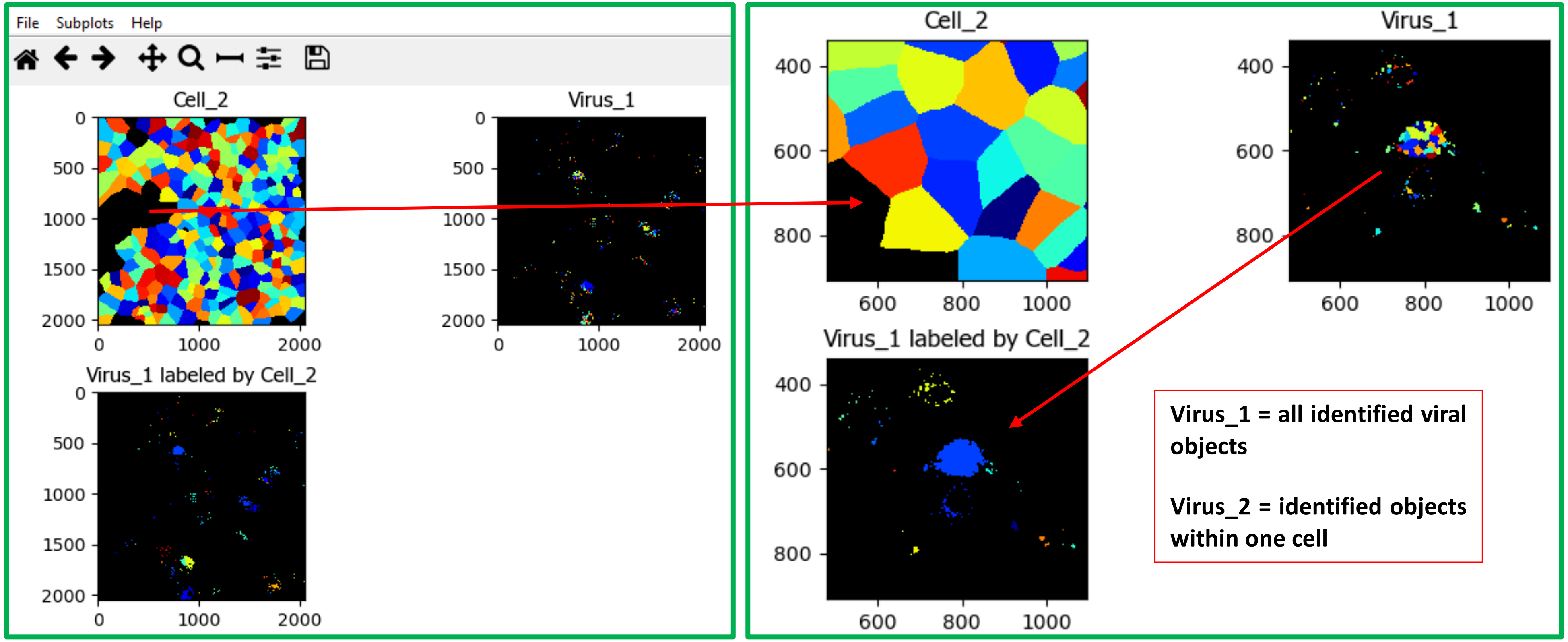

#### < FilterObjects >

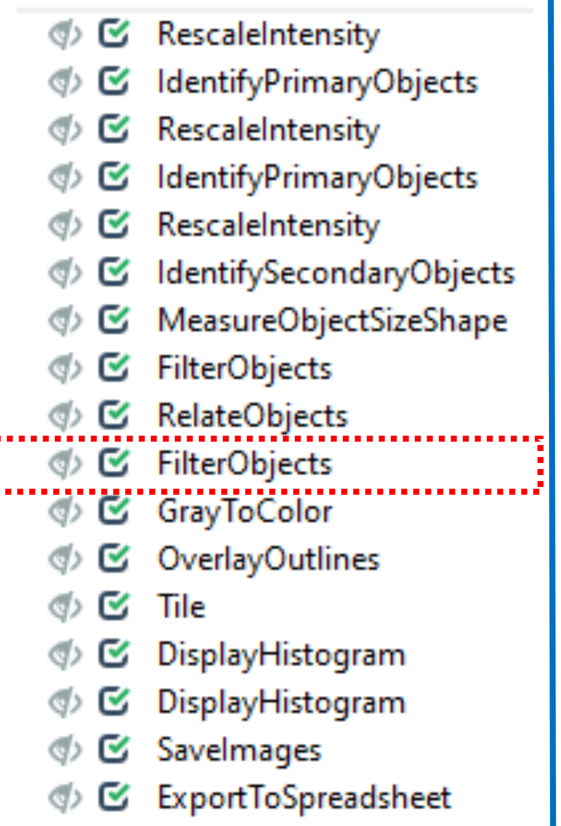


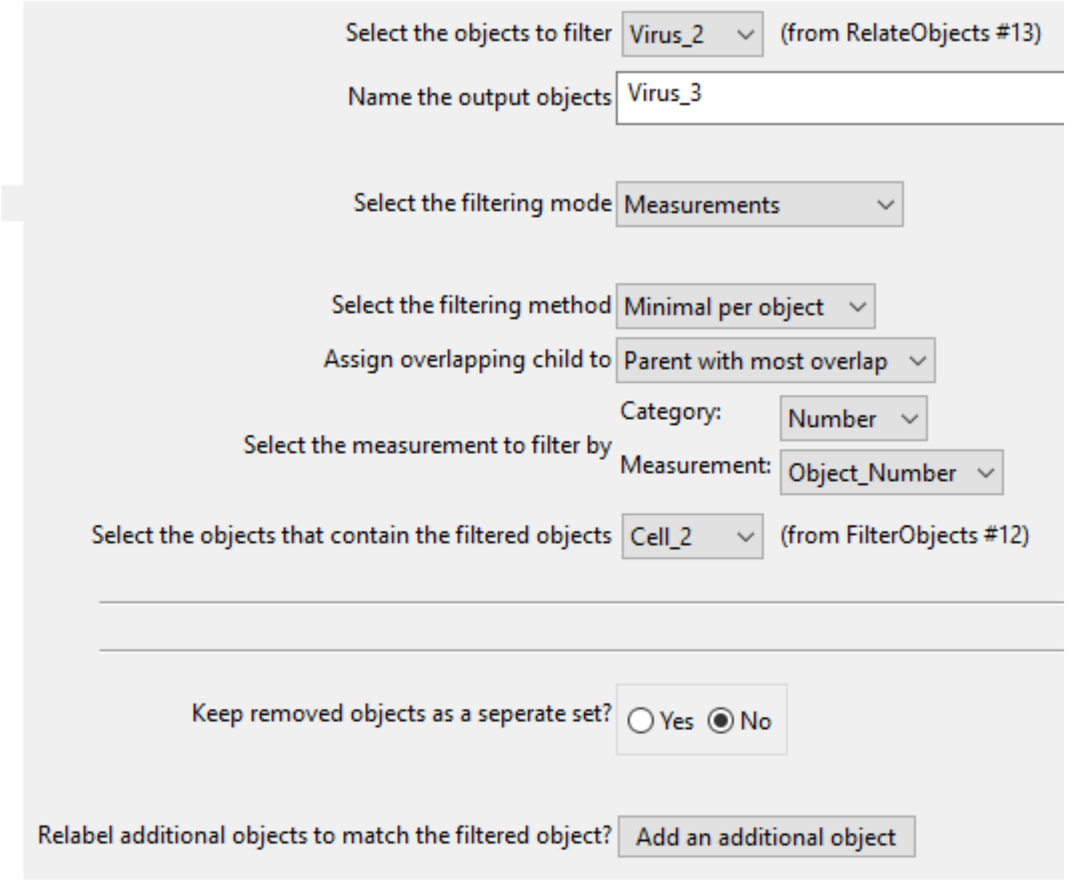


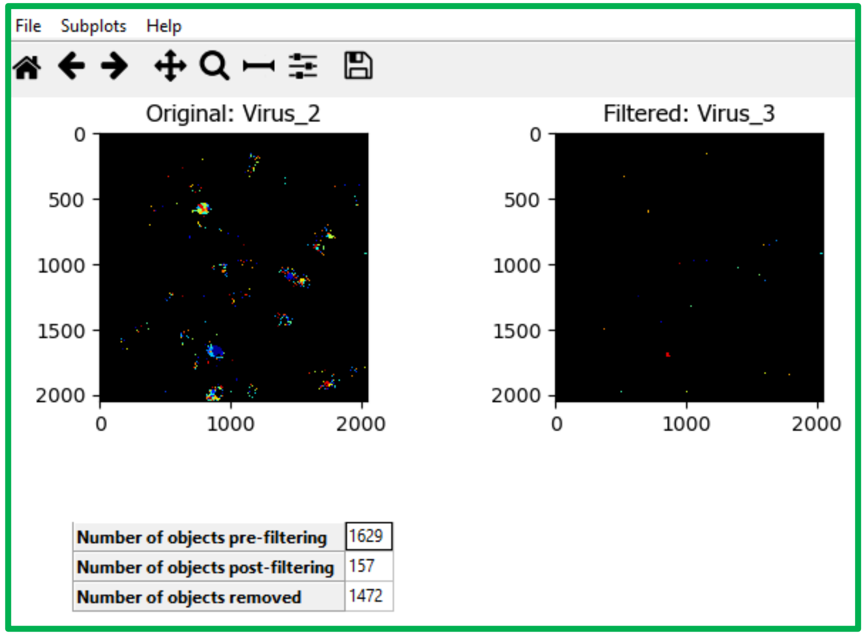


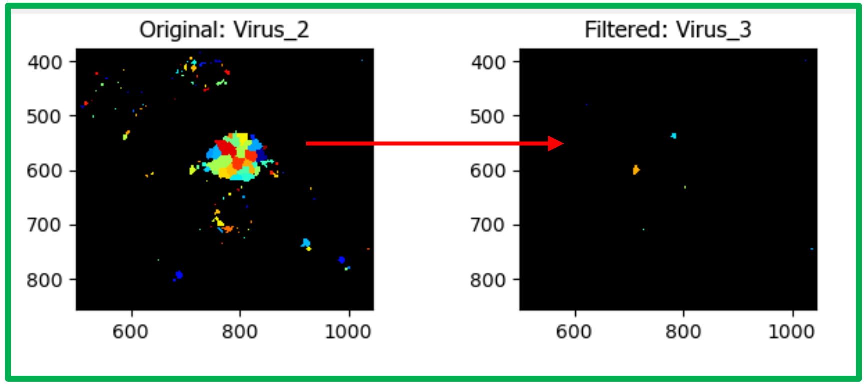


**Virus_2 = identified objects within one cell**

**Virus_3 = one virus object within one cell object = one infected cell**

< GrayToColor >

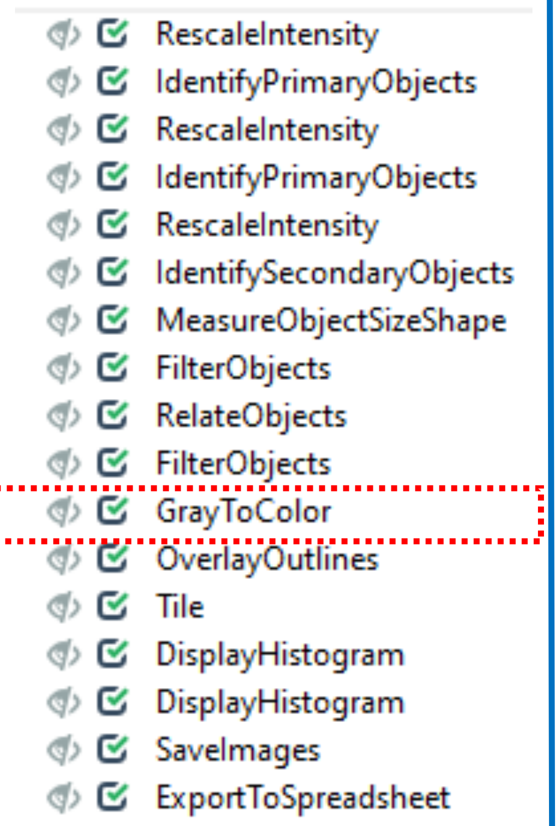


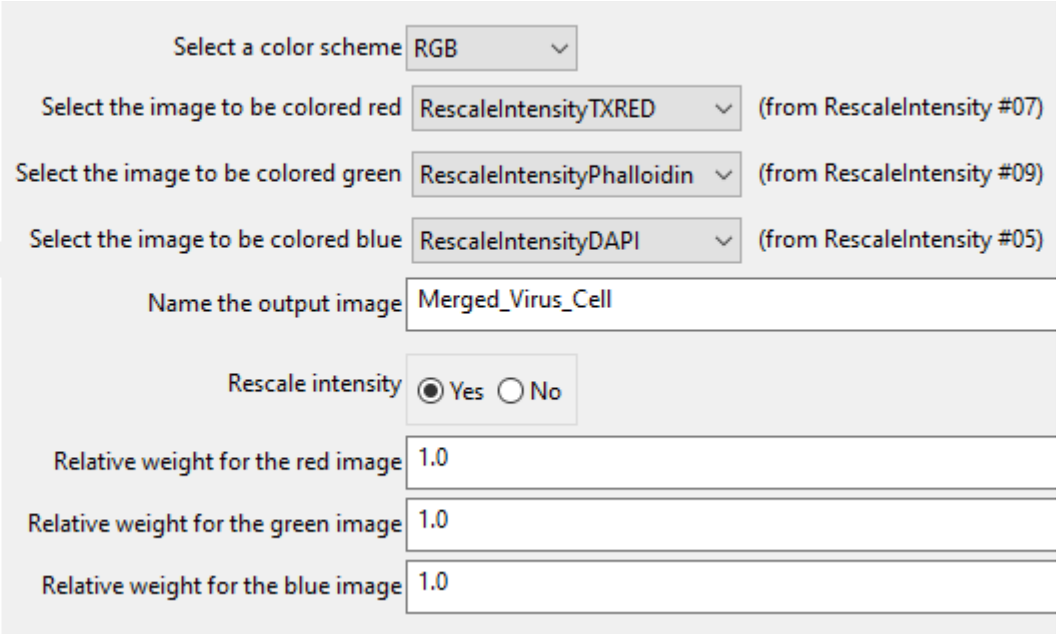


Overlay of channels and colored according to your settings

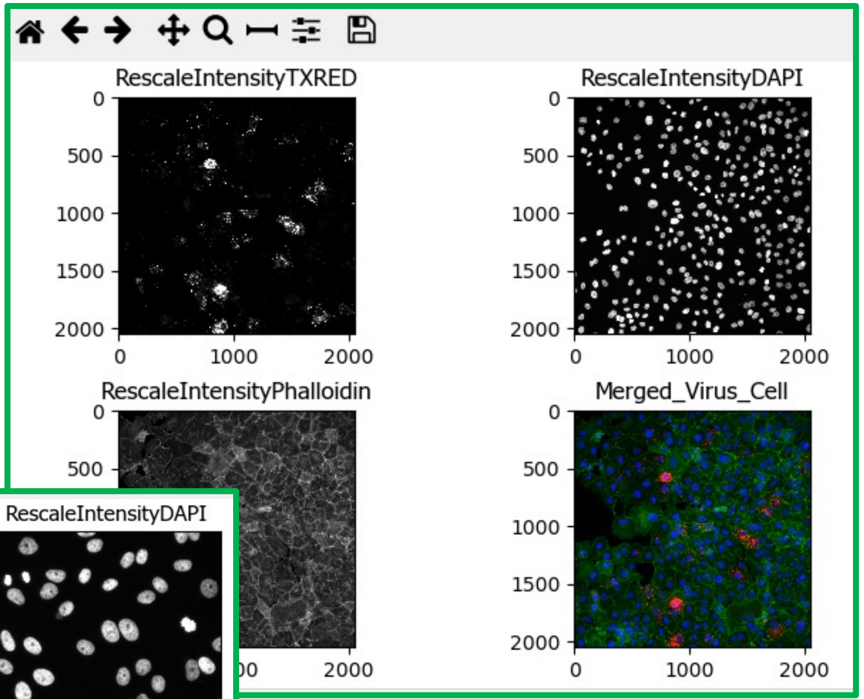


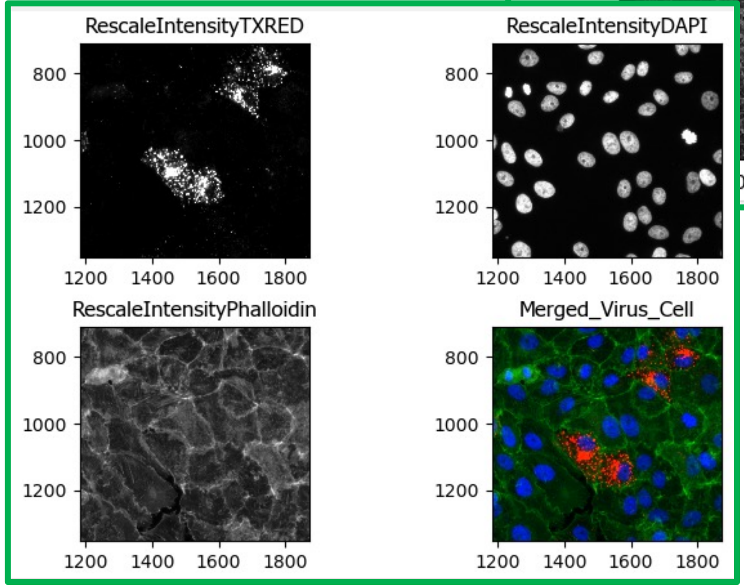

### < OverlayOutlines >

- RescaleIntensity
- IdentifyPrimaryObjects
- RescaleIntensity
- IdentifyPrimaryObjects
- RescaleIntensity
- IdentifySecondaryObjects
- MeasureObjectSizeShape
- FilterObjects
- RelateObjects
- FilterObjects
- GrayToColor
- **OverlayOutlines**
- Tile
- DisplayHistogram
- DisplayHistogram
- SaveImages
- ExportToSpreadsheet

Display outlines on a blank image? (•) Yes ( ) No

Name the output image: Outlines_on_Merge

Outline display mode: Color

How to outline: Inner

Select objects to display: Nucleus (from IdentifyPrimaryObjects #06)

Select outline color: [blue]

Select objects to display: Cell_2 (from FilterObjects #12)

Select outline color: [green]

Remove this outline

Select objects to display: Virus_1 (from IdentifyPrimaryObjects #08)

Select outline color: [red]

Remove this outline

Add another outline

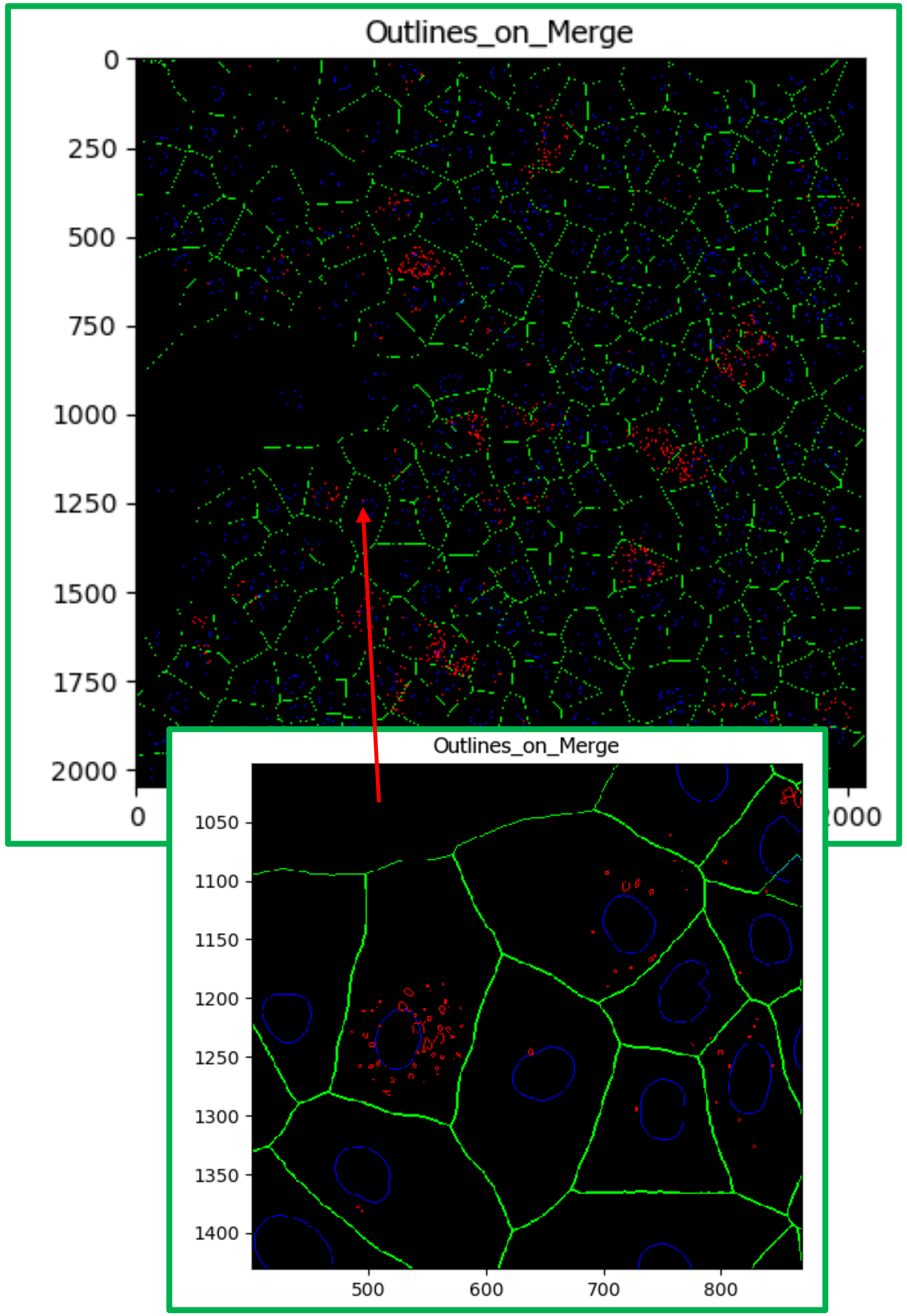

#### < Tile >

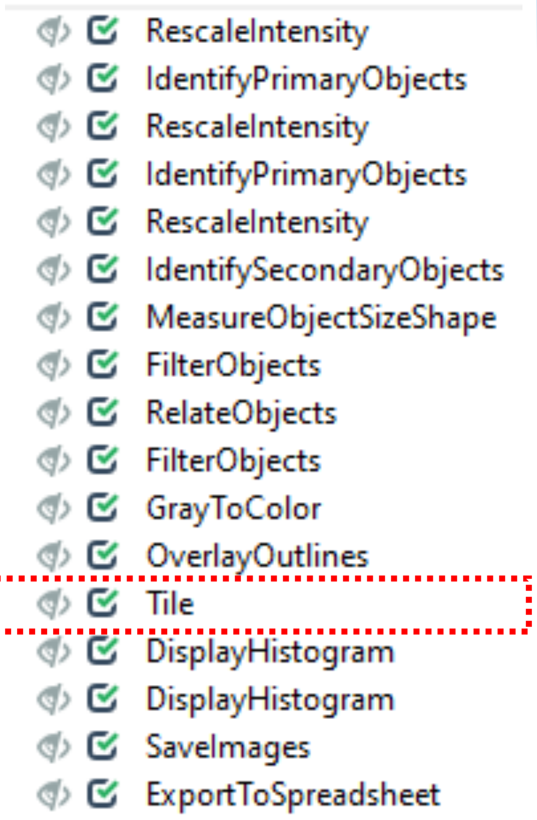
RescaleIntensity
IdentifyPrimaryObjects
RescaleIntensity
IdentifyPrimaryObjects
RescaleIntensity
IdentifySecondaryObjects
MeasureObjectSizeShape
FilterObjects
RelateObjects
FilterObjects
GrayToColor
OverlayOutlines
Tile
DisplayHistogram
DisplayHistogram
SaveImages
ExportToSpreadsheet

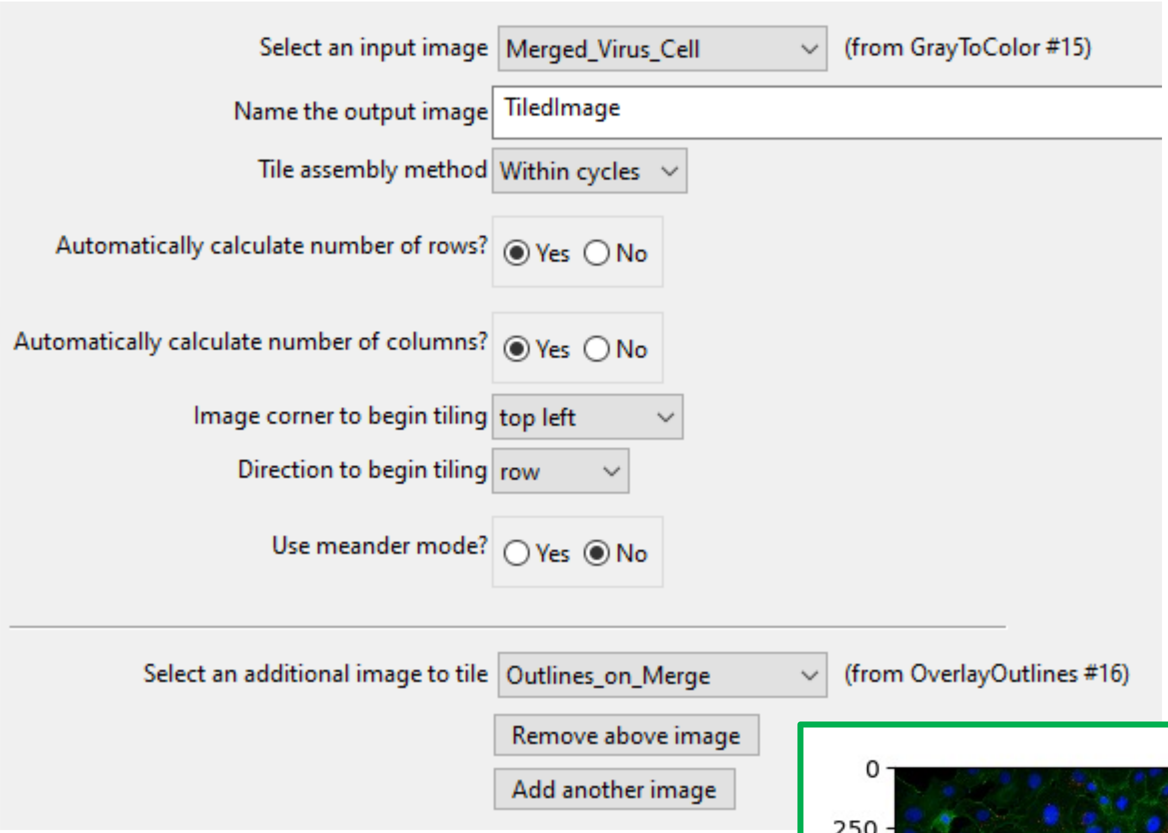
Select an input image
Merged_Virus_Cell
(from GrayToColor #15)
Name the output image
TiledImage
Tile assembly method
Within cycles
Automatically calculate number of rows?
Yes No
Automatically calculate number of columns?
Yes No
Image corner to begin tiling
top left
Direction to begin tiling
row
Use meander mode?
Yes No
Select an additional image to tile
Outlines_on_Merge
(from OverlayOutlines #16)
Remove above image
Add another image

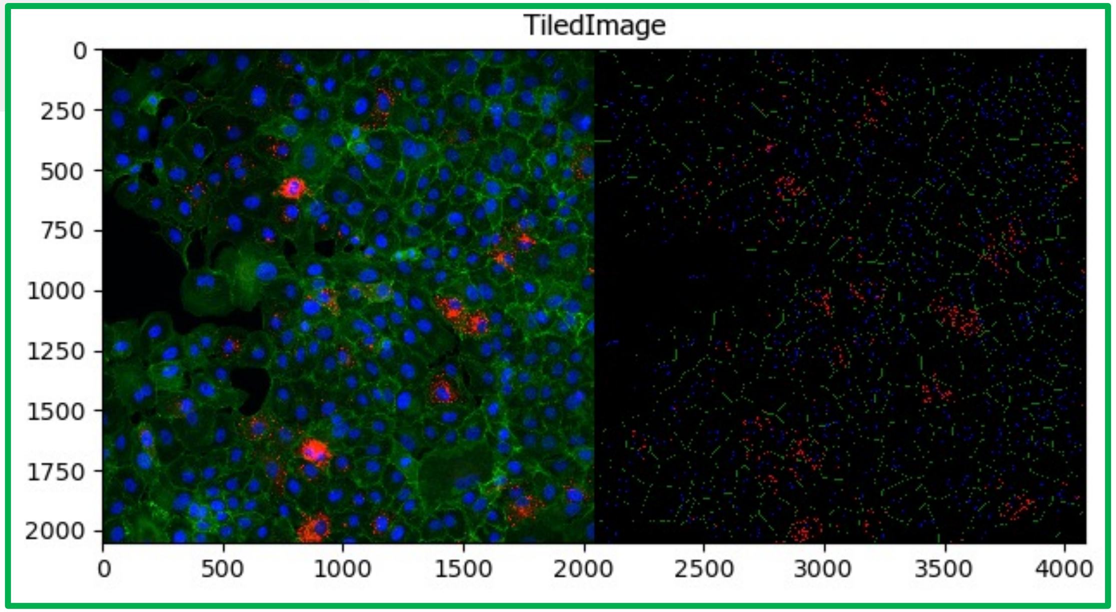
TiledImage
0
250
500
750
1000
1250
1500
1750
2000
0
500
1000
1500
2000
2500
3000
3500
4000

**Optional:** < DisplayHistogram >

- RescaleIntensity
- IdentifyPrimaryObjects
- RescaleIntensity
- IdentifyPrimaryObjects
- RescaleIntensity
- IdentifySecondaryObjects
- MeasureObjectSizeShape
- FilterObjects
- RelateObjects
- FilterObjects
- GrayToColor
- OverlayOutlines
- Tile
- DisplayHistogram
- DisplayHistogram
- SaveImages
- ExportToSpreadsheet

Select the object whose measurements will be displayed: Virus_2 (from RelateObjects #13)

Select the object measurement to plot: Category: Number; Measurement: Object_Number

Number of bins: 1

How should the X-axis be scaled? linear

How should the Y-axis be scaled? linear

Enter a title for the plot, if desired: all idetified viral objects [Nr.]

Specify min/max bounds for the X-axis? Yes / No (selected)

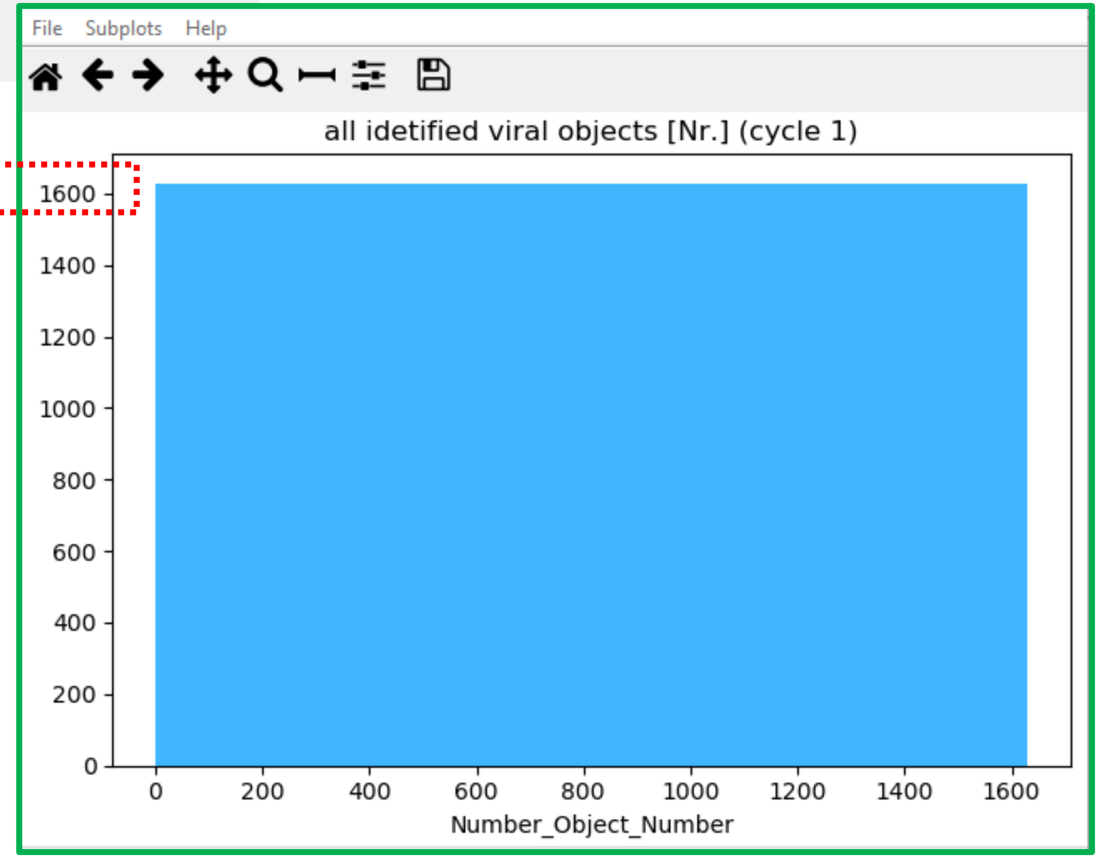

**Optional:** < DisplayHistogram >

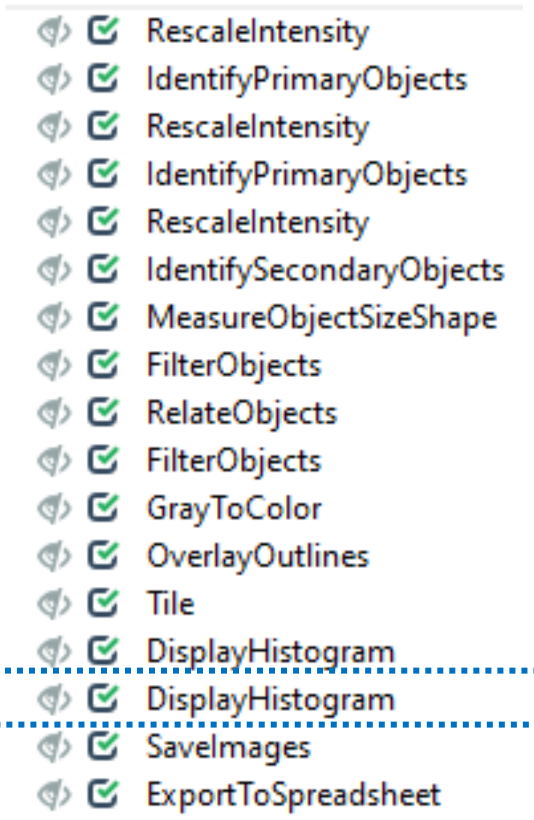


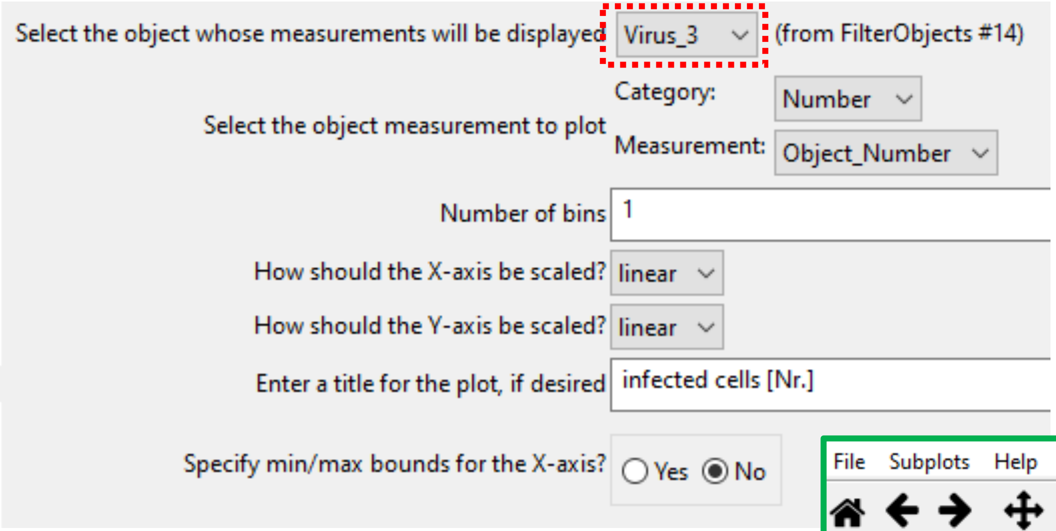


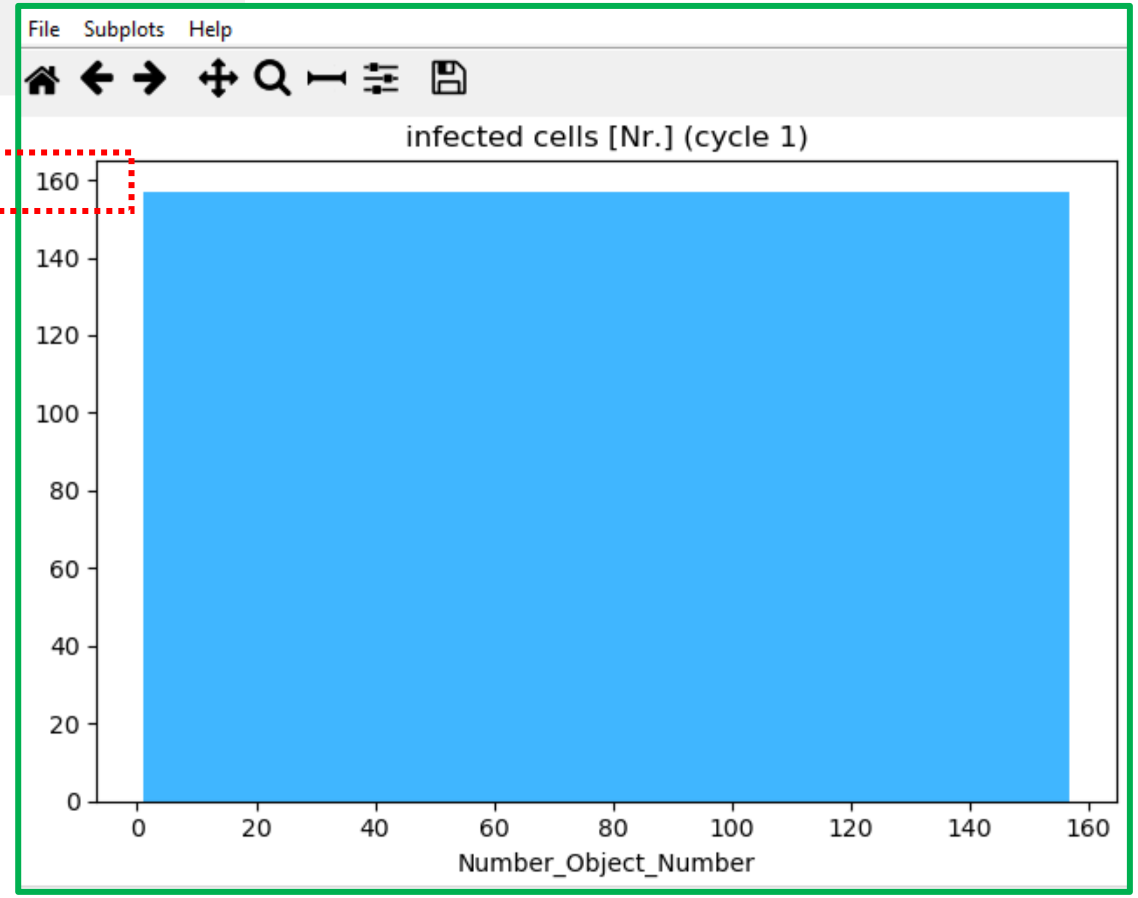

< SaveImages >

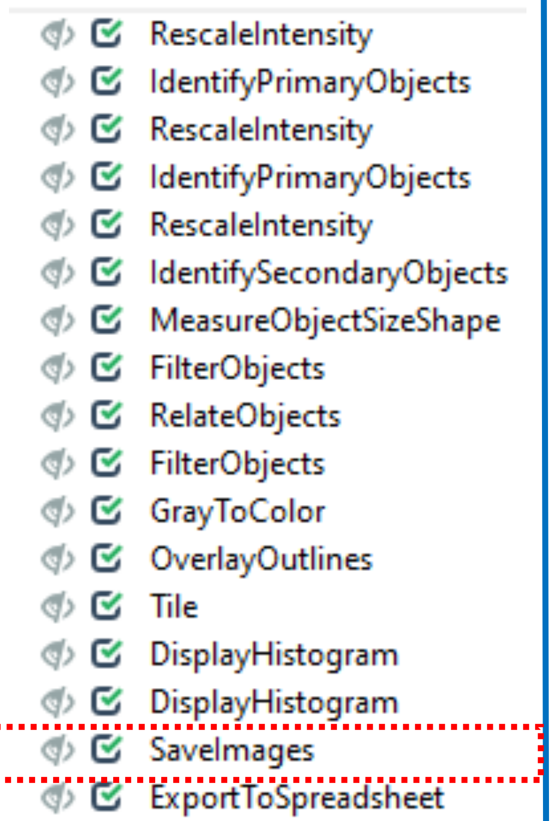
RescaleIntensity
IdentifyPrimaryObjects
RescaleIntensity
IdentifyPrimaryObjects
RescaleIntensity
IdentifySecondaryObjects
MeasureObjectSizeShape
FilterObjects
RelateObjects
FilterObjects
GrayToColor
OverlayOutlines
Tile
DisplayHistogram
DisplayHistogram
SaveImages
ExportToSpreadsheet

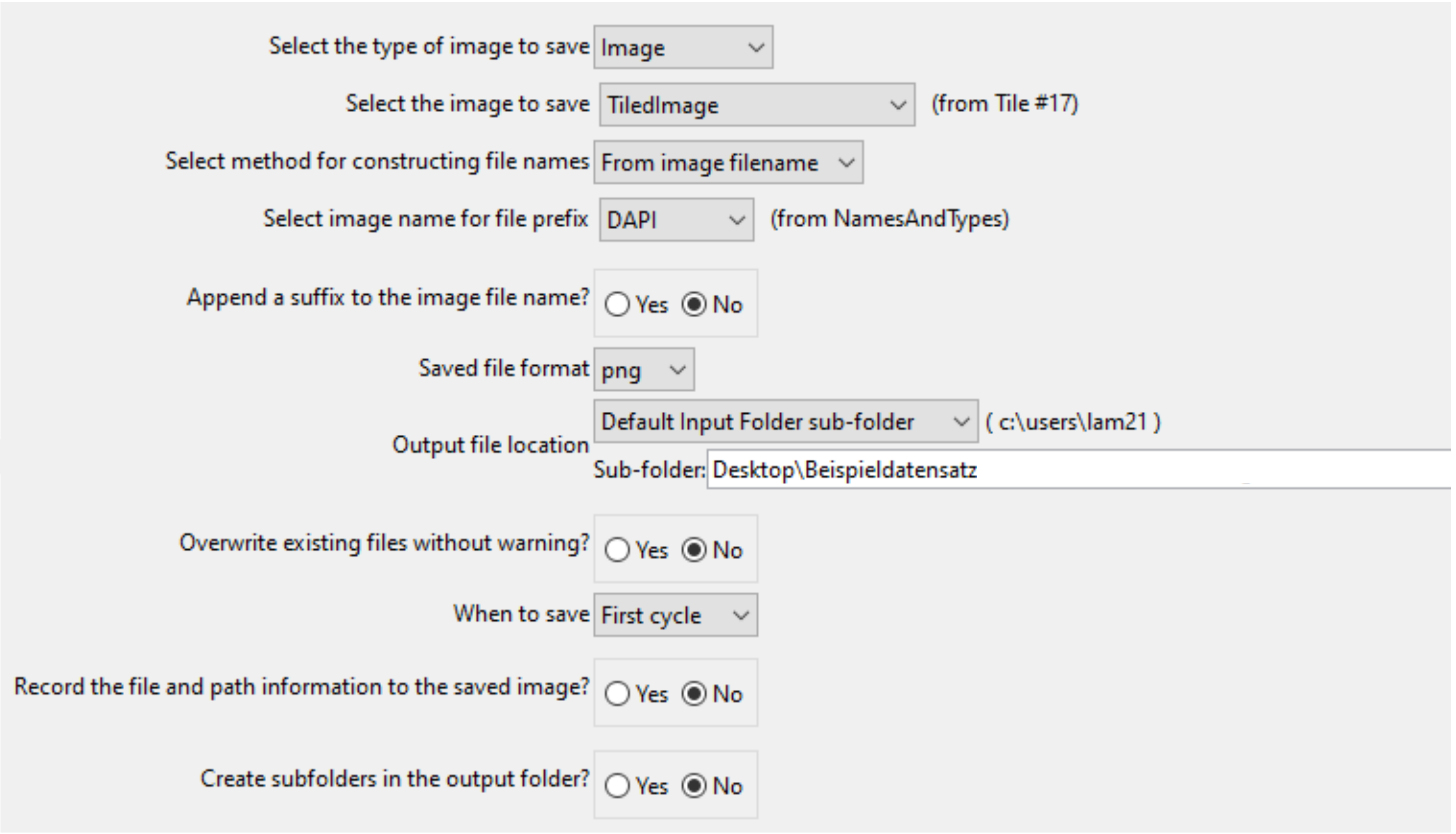
Select the type of image to save
Image
Select the image to save
TiledImage
(from Tile #17)
Select method for constructing file names
From image filename
Select image name for file prefix
DAPI
(from NamesAndTypes)
Append a suffix to the image file name?
Yes
No
Saved file format
png
Output file location
Default Input Folder sub-folder
( c:\users\lam21 )
Sub-folder: Desktop\Beispieldatensatz
Overwrite existing files without warning?
Yes
No
When to save
First cycle
Record the file and path information to the saved image?
Yes
No
Create subfolders in the output folder?
Yes
No

#### <ExportToSpreadsheet>

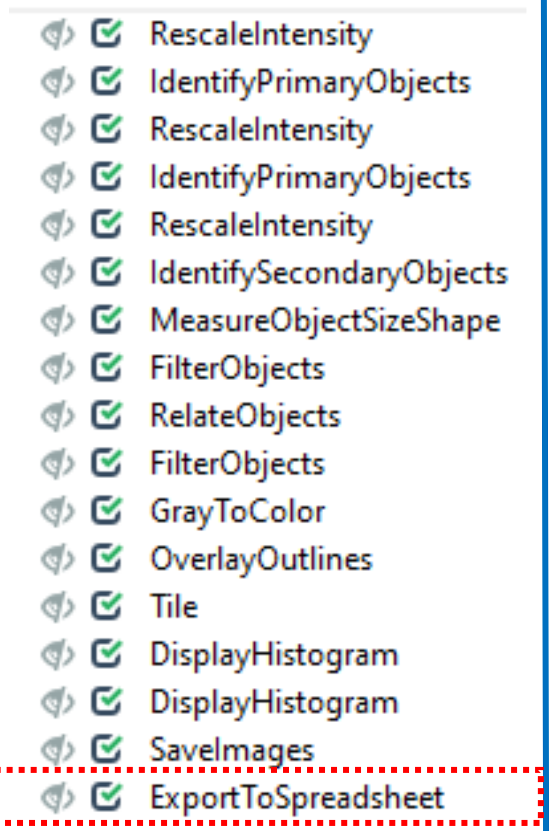


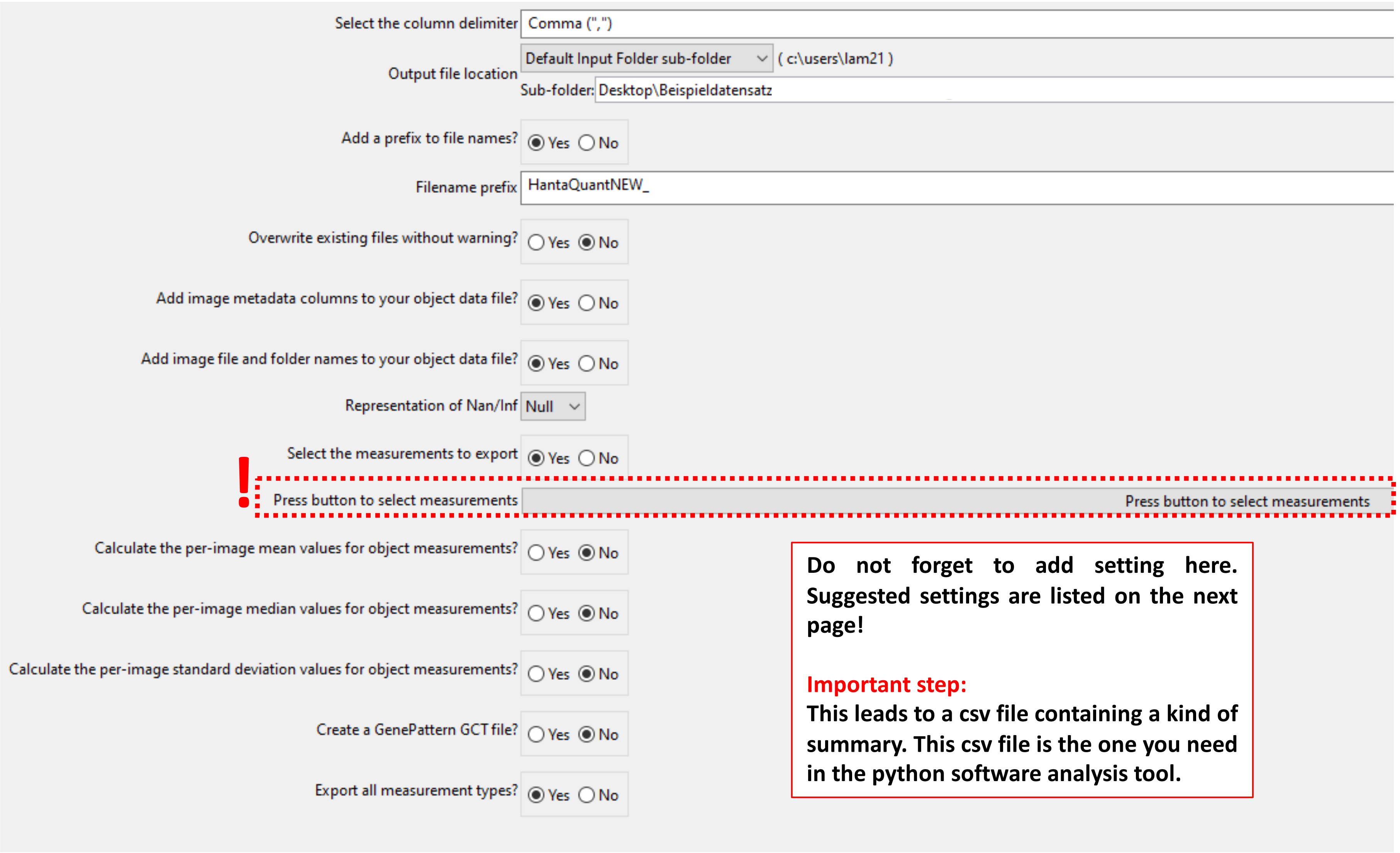


**Do not forget to add setting here. Suggested settings are listed on the next page!**

**Important step:**

**This leads to a csv file containing a kind of summary. This csv file is the one you need in the python software analysis tool.**

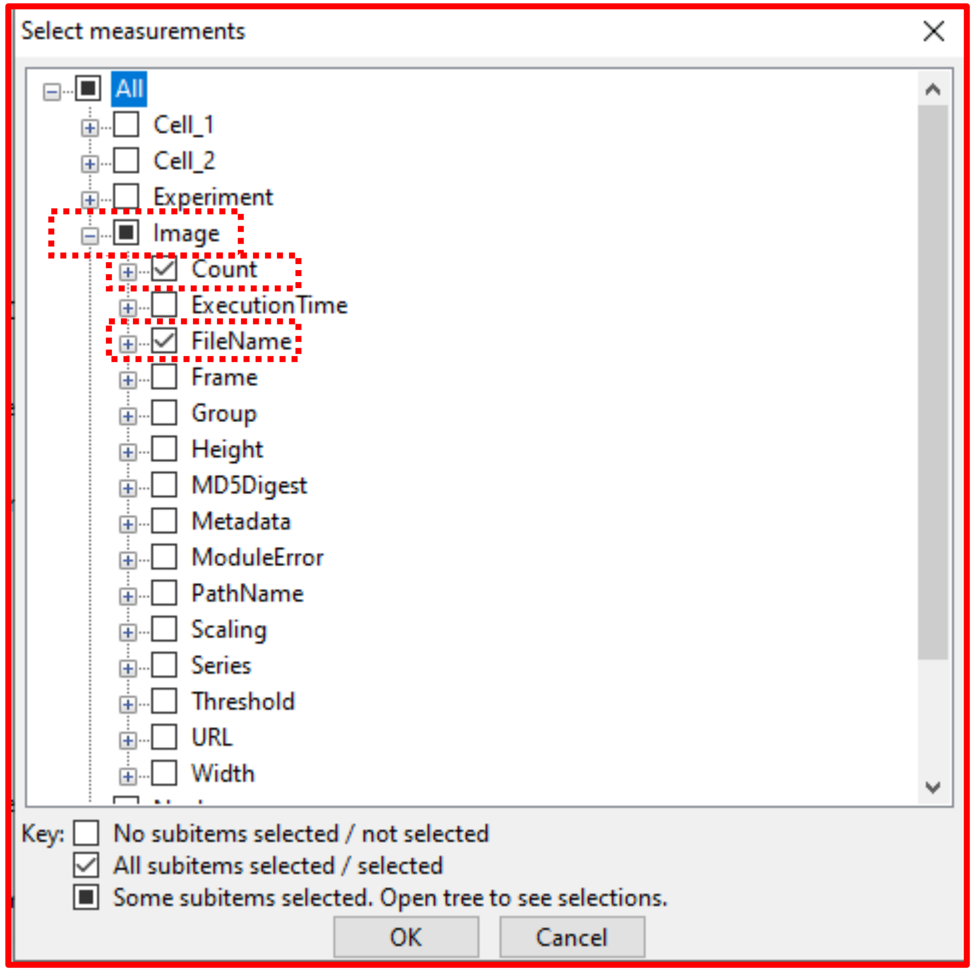


Export to spreadsheet

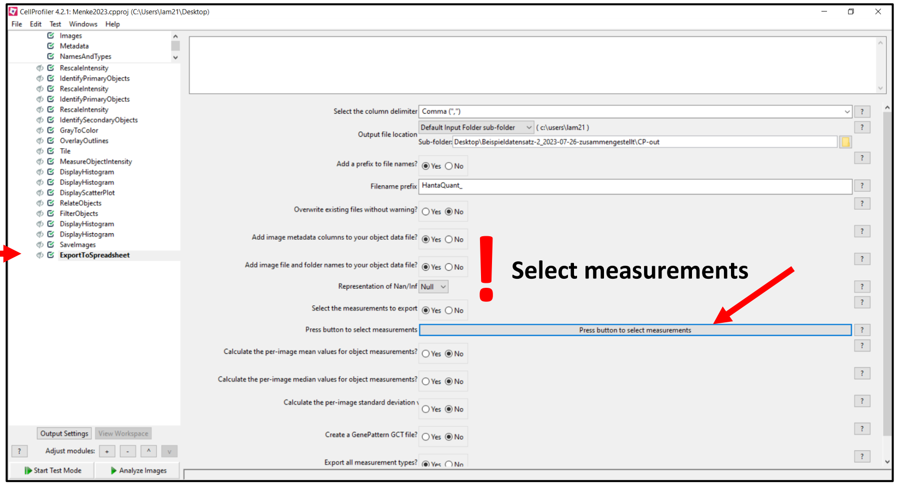


**Important step:**

**This lead to a csv file containing a kind of summary. This csv file is the one you need in the python software analysis tool.**

< Exit Test Mode > and start < Analyze Images >

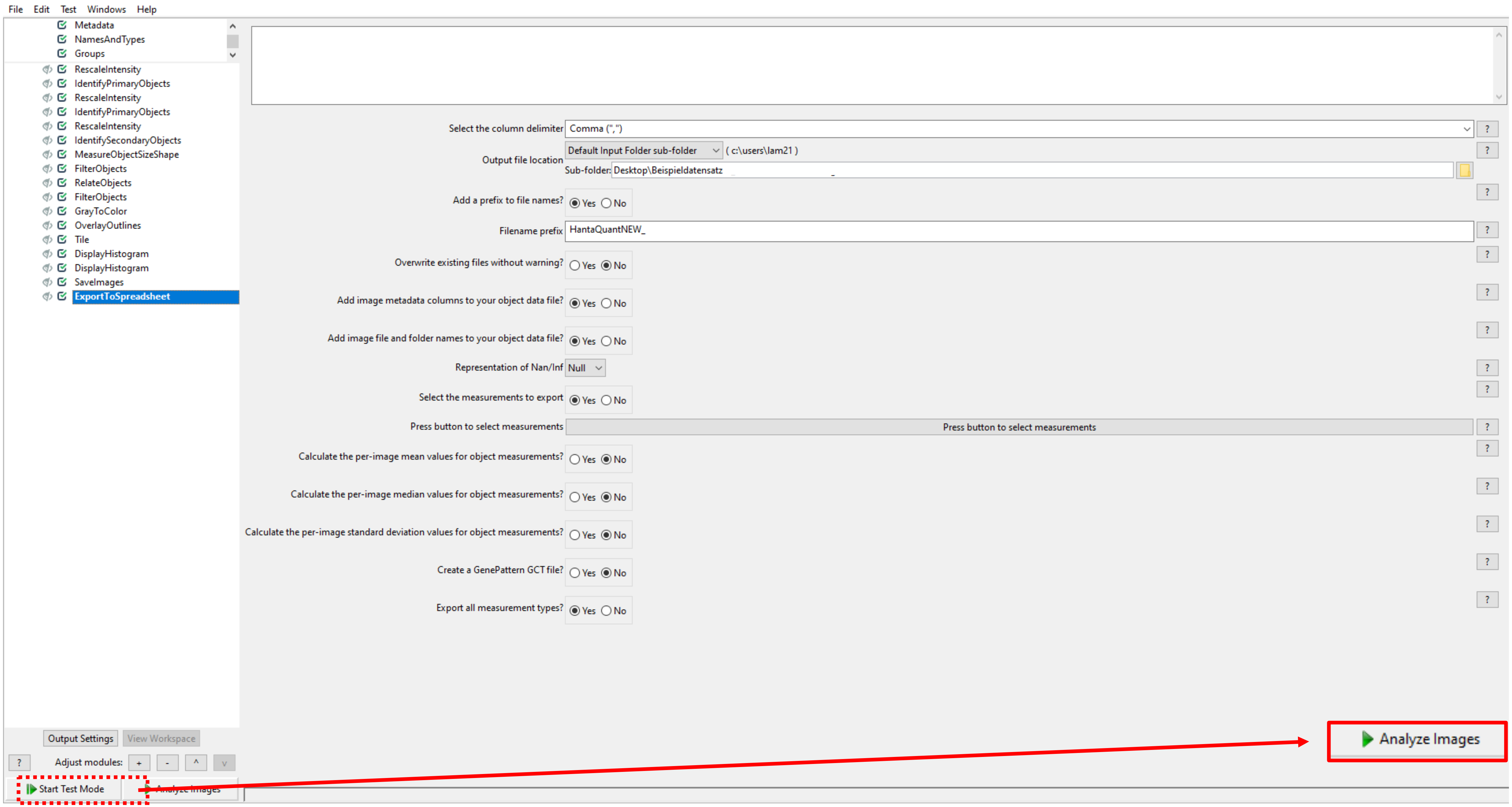

**In your output folder will be saved several csv files, as well as one png file containing the saved TiledImage**

| Name | Type |
|---|---|
| 2023-05-17_Vero_NEGATIV_48h_well-B_20x_000_C=2 | PNG File |
| HantaQuant_Cell | Microsoft Excel Comma... |
| HantaQuant_Experiment | Microsoft Excel Comma... |
| HantaQuant_FilterObjects_kept | Microsoft Excel Comma... |
| HantaQuant_Image | Microsoft Excel Comma... |
| HantaQuant_Nucleus | Microsoft Excel Comma... |
| HantaQuant_RelateObjects_VirusWithinCell | Microsoft Excel Comma... |
| HantaQuant_RemovedObjects | Microsoft Excel Comma... |
| HantaQuant_Virus | Microsoft Excel Comma... |

Xy_Image: would be the summary file, which could be used in the python analysis software tool

### Datalyze

**Software tool to analyze your data**

**Aim of this program:**

*sort necessary data for quantification to summarize and store data in a new file and show results graphically as a scatterplot and as a bar graph*

Raw data from CellProfiler:

```
Count_Cell,Count_FilterObjects_kept,Count_Nucleus,Count_RelateObjects_VirusWithinCell,Count_RemovedObjects,Count_Virus,FileName_DAPI,FileName_Phalloidin,FileName_TXRED,ImageNumber,Meta
335.0,11.0,335.0,15.0,4.0,15.0,2023-05-17_Vero_NEGATIV_48h_well-B_20x_020_C=2.tif,2023-05-17_Vero_NEGATIV_48h_well-B_20x_020_C=0.tif,2023-05-17_Vero_NEGATIV_48h_well-B_20x_020_C=1.
334.0,15.0,334.0,20.0,5.0,21.0,2023-05-17_Vero_NEGATIV_48h_well-B_20x_022_C=2.tif,2023-05-17_Vero_NEGATIV_48h_well-B_20x_022_C=0.tif,2023-05-17_Vero_NEGATIV_48h_well-B_20x_022_C=1.
327.0,212.0,327.0,2710.0,2498.0,2756.0,2023-05-17_Vero_TULV-M_48h_well-A_20x_000_C=2.tif,2023-05-17_Vero_TULV-M_48h_well-A_20x_000_C=0.tif,2023-05-17_Vero_TULV-M_48h_well-A_20x_00
218.0,138.0,218.0,1375.0,1237.0,1447.0,2023-05-17_Vero_TULV-M_48h_well-A_20x_001_C=2.tif,2023-05-17_Vero_TULV-M_48h_well-A_20x_001_C=0.tif,2023-05-17_Vero_TULV-M_48h_well-A_20x_00
365.0,244.0,365.0,2451.0,2207.0,2463.0,2023-05-17_Vero_TULV-M_48h_well-A_20x_002_C=2.tif,2023-05-17_Vero_TULV-M_48h_well-A_20x_002_C=0.tif,2023-05-17_Vero_TULV-M_48h_well-A_20x_00
328.0,204.0,328.0,2262.0,2058.0,2406.0,2023-05-17_Vero_TULV-M_48h_well-A_20x_003_C=2.tif,2023-05-17_Vero_TULV-M_48h_well-A_20x_003_C=0.tif,2023-05-17_Vero_TULV-M_48h_well-A_20x_00
394.0,252.0,394.0,2743.0,2491.0,2905.0,2023-05-17_Vero_TULV-M_48h_well-A_20x_004_C=2.tif,2023-05-17_Vero_TULV-M_48h_well-A_20x_004_C=0.tif,2023-05-17_Vero_TULV-M_48h_well-A_20x_00
663.0,330.0,663.0,2099.0,1769.0,2189.0,2023-05-19_Vero_PUUV-V_48h_well-A_20x_000_C=2.tif,2023-05-19_Vero_PUUV-V_48h_well-A_20x_000_C=0.tif,2023-05-19_Vero_PUUV-V_48h_well-A_20x_00
624.0,320.0,624.0,2007.0,1687.0,2087.0,2023-05-19_Vero_PUUV-V_48h_well-A_20x_001_C=2.tif,2023-05-19_Vero_PUUV-V_48h_well-A_20x_001_C=0.tif,2023-05-19_Vero_PUUV-V_48h_well-A_20x_00
648.0,336.0,648.0,2012.0,1676.0,2103.0,2023-05-19_Vero_PUUV-V_48h_well-A_20x_002_C=2.tif,2023-05-19_Vero_PUUV-V_48h_well-A_20x_002_C=0.tif,2023-05-19_Vero_PUUV-V_48h_well-A_20x_00
726.0,376.0,726.0,2123.0,1747.0,2199.0,2023-05-19_Vero_PUUV-V_48h_well-A_20x_003_C=2.tif,2023-05-19_Vero_PUUV-V_48h_well-A_20x_003_C=0.tif,2023-05-19_Vero_PUUV-V_48h_well-A_20x_00
767.0,380.0,767.0,2355.0,1975.0,2393.0,2023-05-19_Vero_PUUV-V_48h_well-A_20x_004_C=2.tif,2023-05-19_Vero_PUUV-V_48h_well-A_20x_004_C=0.tif,2023-05-19_Vero_PUUV-V_48h_well-A_20x_00
```

Keep important columns and process data

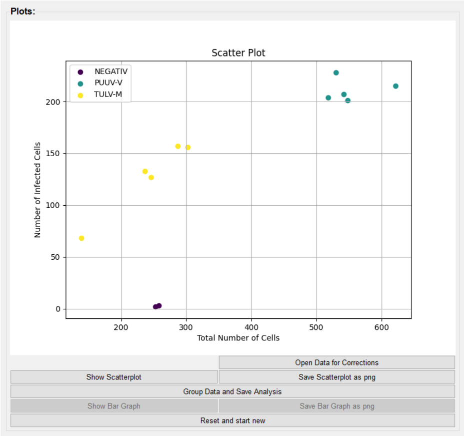


```
Total Number of Cells,Number of Infected Cells,Name,Percentage of Infected Cells [%]
335.0,11.0,NEGATIV,3.28
334.0,15.0,NEGATIV,4.49
327.0,212.0,TULV-M,64.83
218.0,138.0,TULV-M,63.3
365.0,244.0,TULV-M,66.85
328.0,204.0,TULV-M,62.2
394.0,252.0,TULV-M,63.96
663.0,330.0,PUUV-V,49.77
624.0,320.0,PUUV-V,51.28
648.0,336.0,PUUV-V,51.85
726.0,376.0,PUUV-V,51.79
767.0,380.0,PUUV-V,49.54
```

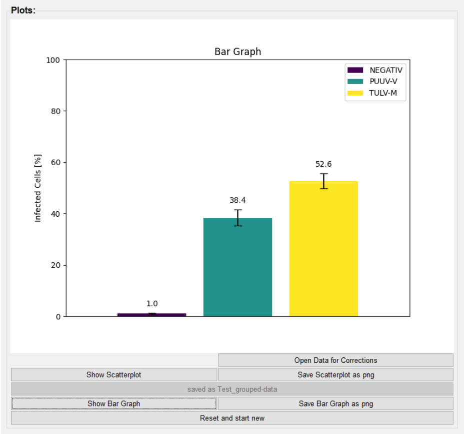


```
Name,Mean Percentage of Infected Cells [%],Standard Deviation (SD)
NEGATIV,3.88,0.86
PUUV-V,50.85,1.11
TULV-M,64.23,1.75
```

Path of file: ***Copy and Paste file path as text***

Name of file: ***Copy and Paste file name***

Name for new file: ***Enter the Name for the new document – it will be saved in the same folder as your loaded file***

Old Name from your Cell-Profiler-Pipeline will be renamed in:

- Name
- Number of Cells
- Number of infected Cells

- Percentage of infected cells will be calculated by (Number-of-infected-cells/number-of-cells) * 100

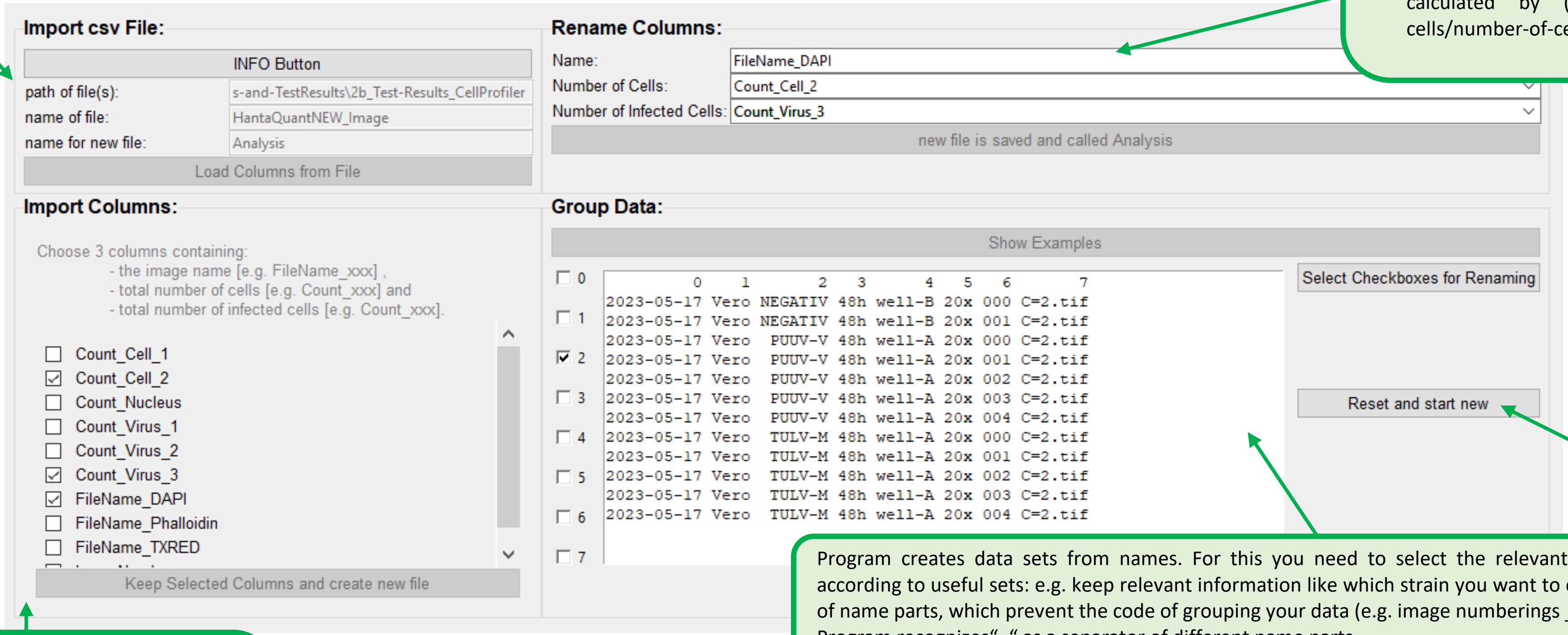


To Start again from the Top

- Enables to correct column choice
- Enables to adapt Renaming

Program creates data sets from names. For this you need to select the relevant information to group your data according to useful sets: e.g. keep relevant information like which strain you want to compare (PUUV/TULV), but get rid of name parts, which prevent the code of grouping your data (e.g. image numberings _022_C=0.tif ).
Program recognizes"_" as a separator of different name parts.

To proceed your data it is needed to set the basic parameters.

- The ***sample name*** e.g. „FileName_DAPI"
- The different ***counted values*** e.g. „Count_Cell_2" and "Count_Virus_3"

- All N/A will be changed to 0.

| Original | Delete letters: | Group data for MEAN and SD |
|---|---|---|
| 2023-05-17_Vero_PUUV-V_48h_well-A_20x_022_C=2.tif | ~~2023-05-17_Vero_~~PUUV-V~~_48h_well-A_20x_022_C=2.tif~~ | PUUV-V |
| 2023-05-17_Vero_PUUV-V_48h_well-A_20x_023_C=2.tif | ~~2023-05-17_Vero_~~PUUV-V~~_48h_well-A_20x_023_C=2.tif~~ | |
| 2023-05-17_Vero_PUUV-V_48h_well-A_20x_024_C=2.tif | ~~2023-05-17_Vero_~~PUUV-V~~_48h_well-A_20x_024_C=2.tif~~ | |
| 2023-05-17_Vero_TULV-M_48h_well-A_20x_025_C=2.tif | ~~2023-05-17_Vero_~~TULV-M~~_48h_well-A_20x_025_C=2.tif~~ | TULV-M |
| 2023-05-17_Vero_TULV-M_48h_well-A_20x_026_C=2.tif | ~~2023-05-17_Vero_~~TULV-M~~_48h_well-A_20x_026_C=2.tif~~ | |
| 2023-05-17_Vero_TULV-M_48h_well-A_20x_027_C=2.tif | ~~2023-05-17_Vero_~~TULV-M~~_48h_well-A_20x_027_C=2.tif~~ | |

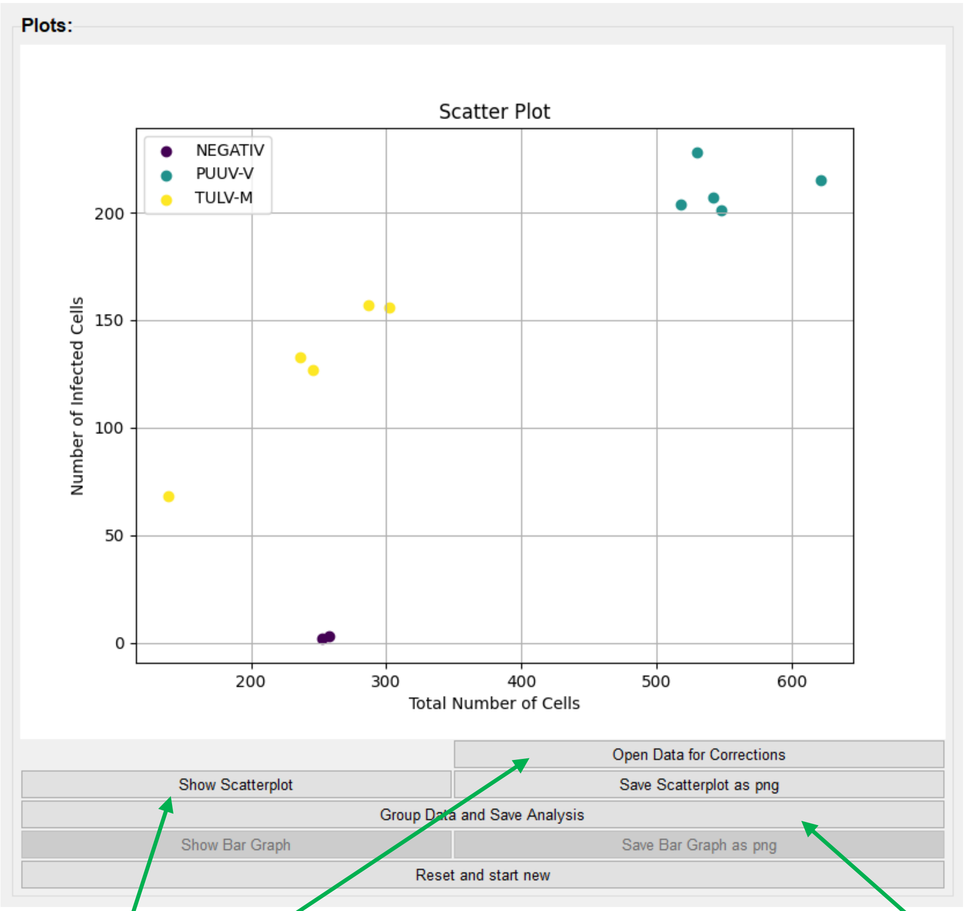


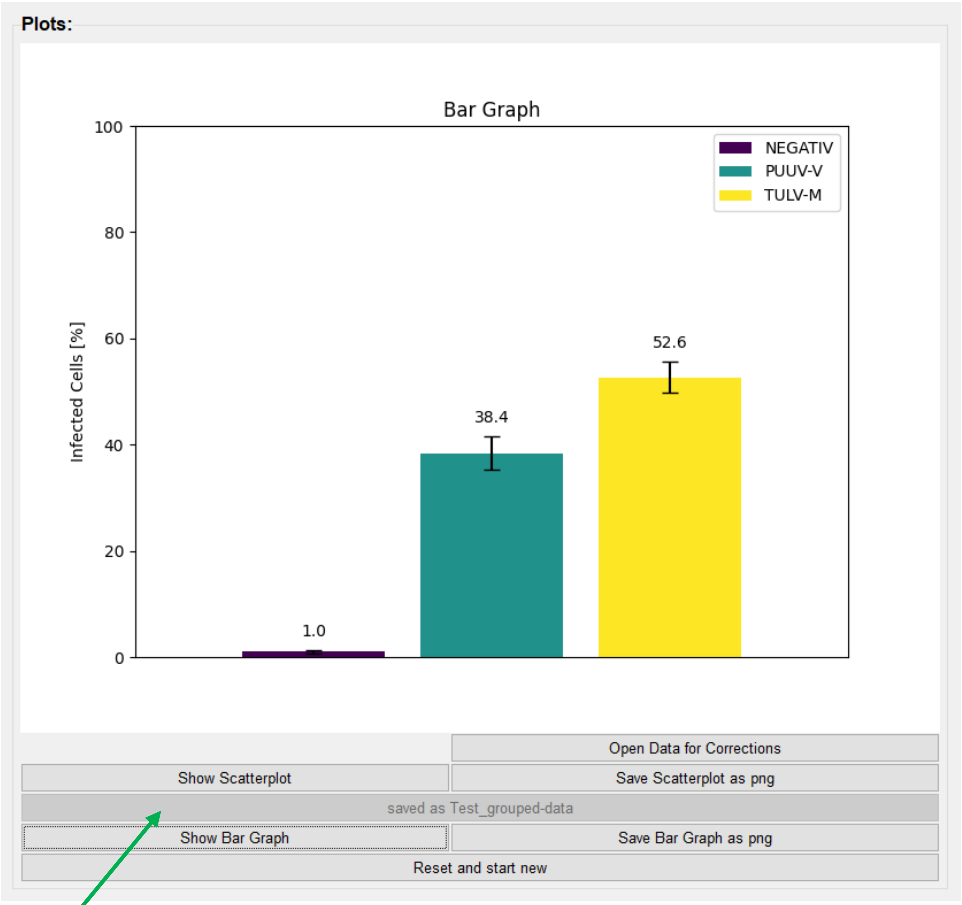


< Open Data for Corrections > **opens your new csv file, so that you could delete outliers. Make sure that you save the file without changing the file name and close it properly. After closing the document you can update your scatterplot by clicking again on** < Show Scatterplot >

< Group Data and Save Analysis > **will then convert and summarize all the grouped data and store it as a new sheet in the document.**

< Reset and start new >
To Start again from former window/ start new
- Enables to process another file
- Enables to correct column choice
- Enables to adapt Renaming and therefore grouping
