## Supplementary Software for "An improved workflow for the quantification of orthohantavirus infection using automated imaging and flow cytometry": Tutorial-NisElements_Datalyze.pdf

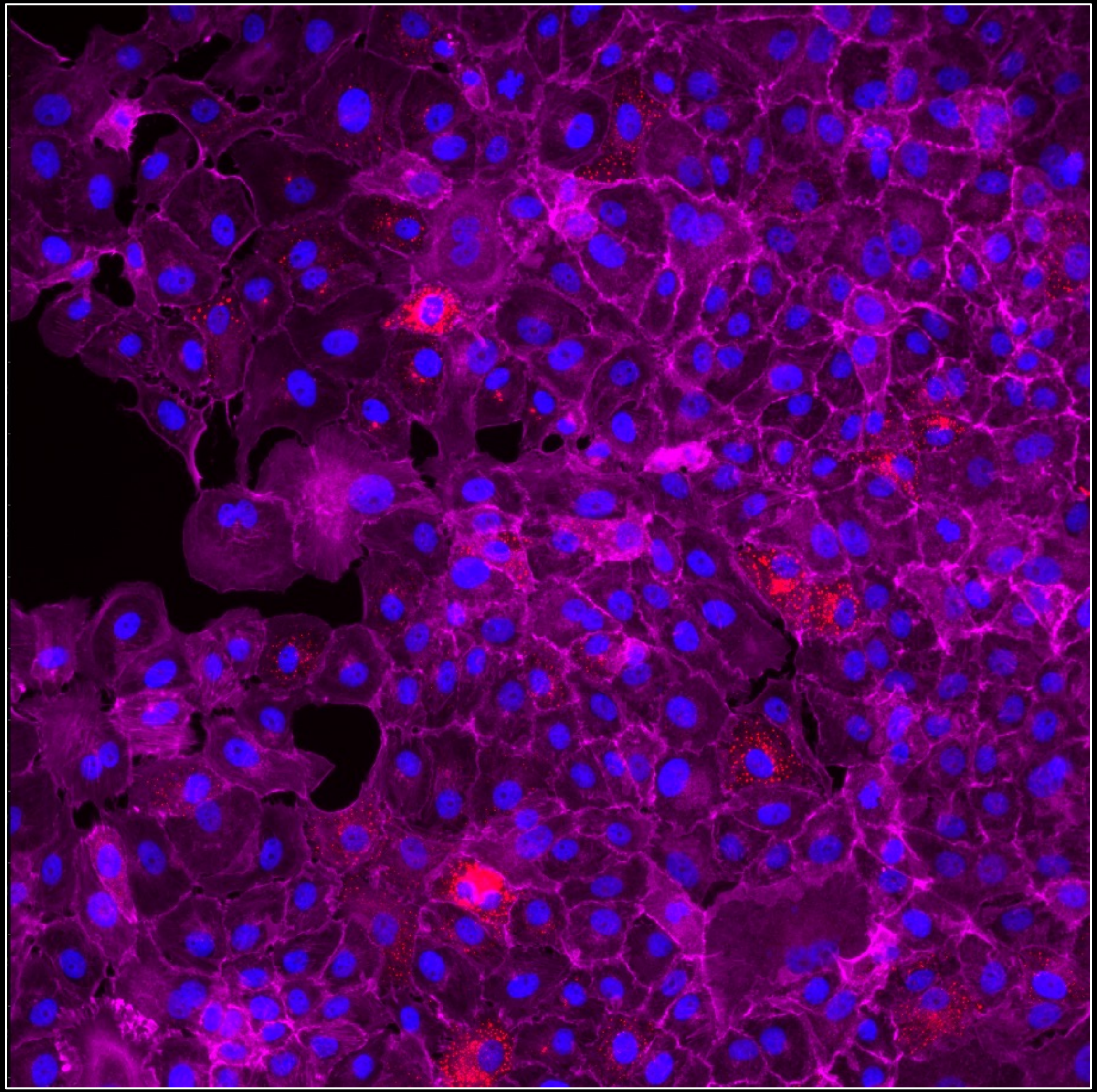

2023-05-17_Vero_TULV-M_48h_well-A_20x_002

### Tutorial for image analysis and virus quantification using CellProfiler

< minimal requirements >

NIS-Elements AR
„Advanced Research“
Imaging Software

### Necessary staining:

Infobox:
- Tips for using the published software Datalyze:
  - Choose your image name by using "_" as a separator
  - e.g. "2023-08-20_VeroE6_PUUV"
  - By recognizing "_" the script will give you the opportunity to get rid of different name parts

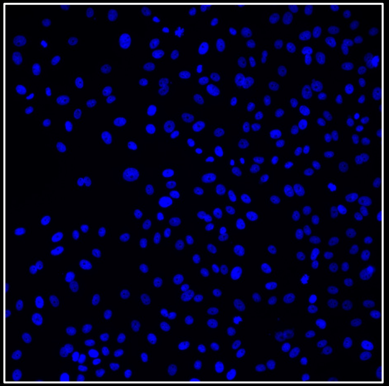

#### Nucleus staining

e.g Hoechst or DAPI staining

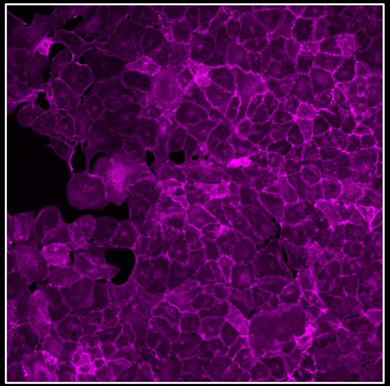

#### Cell borders

e.g F-Actin staining with Phalloidin

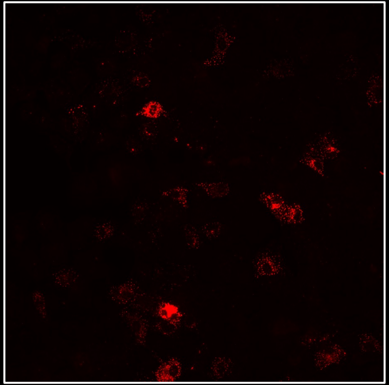

#### Virus staining

e.g. staining of viral nucleocapsid protein (N)

### First Steps:

- Load an image into NisElements
- Open > Image > Analysis Explorer
- Start > Create New > General Analysis

- Helpful to open also LUTs
  > View > Visualization Controls > LUTs

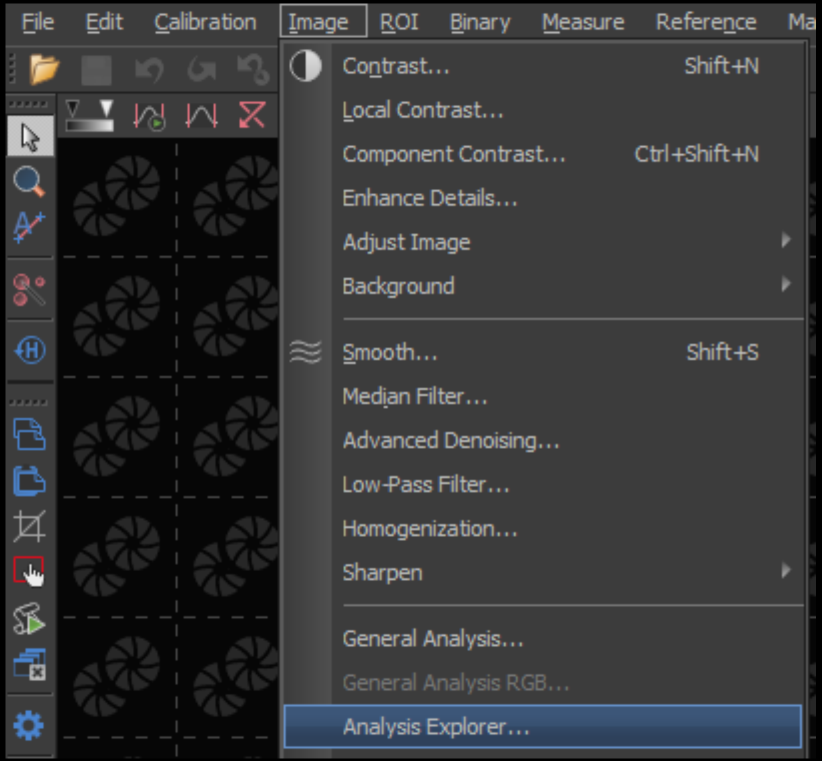


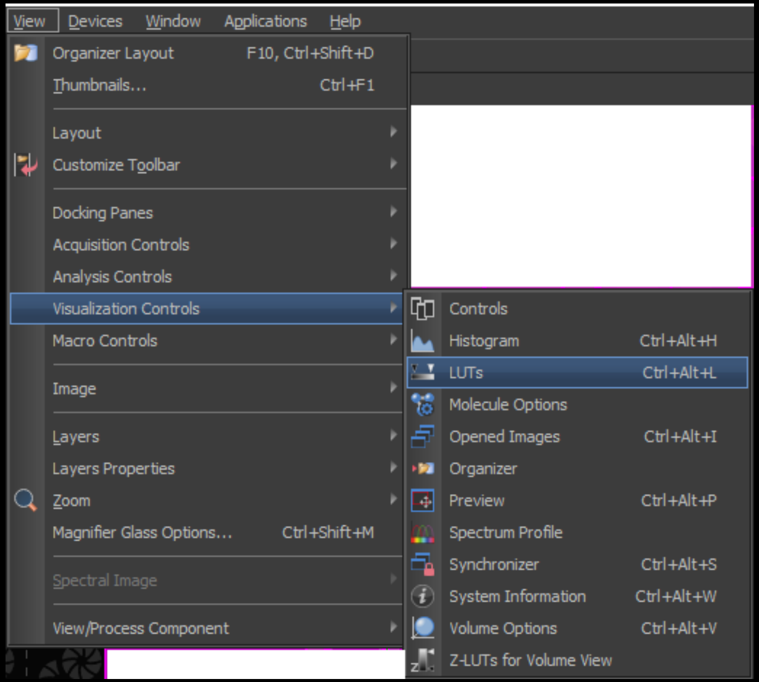


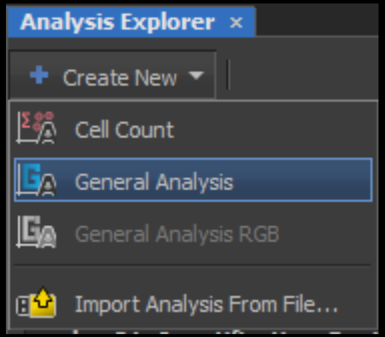

### First Steps:

- Helpful to change transparency

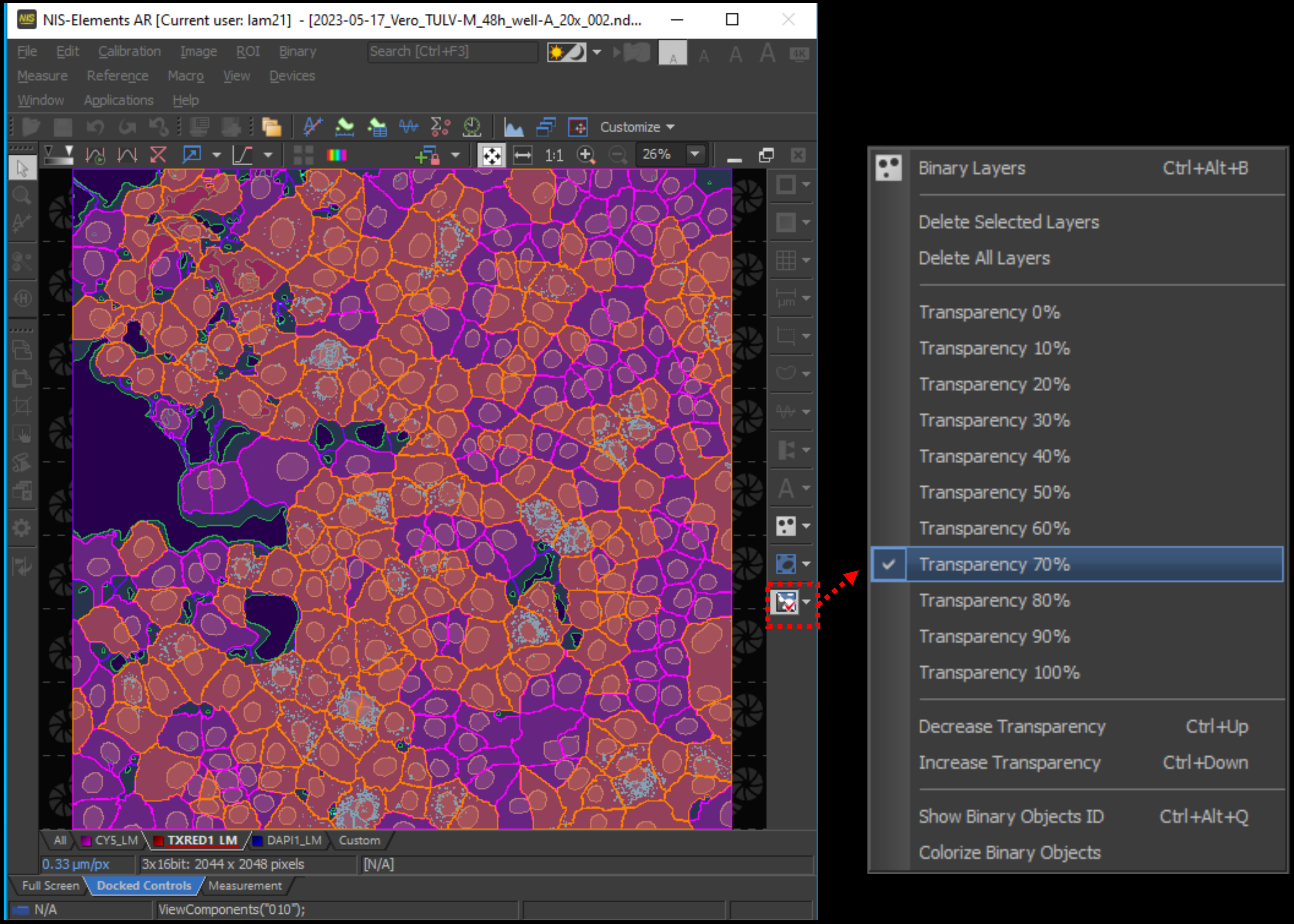

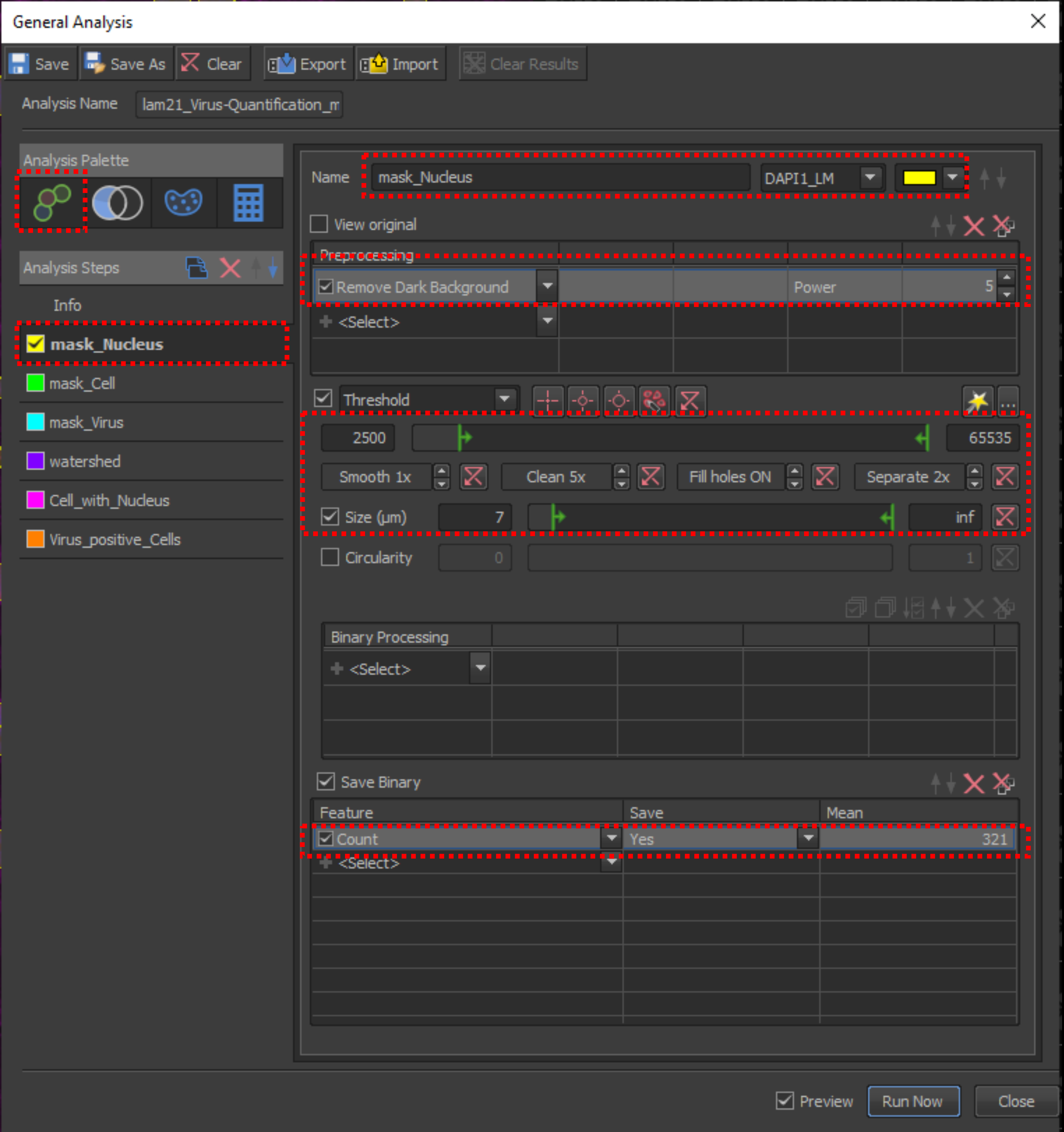
General Analysis
Save
Save As
Clear
Export
Import
Clear Results
Analysis Name
lam21_Virus-Quantification_m
Analysis Palette
Analysis Steps
Info
mask_Nucleus
mask_Cell
mask_Virus
watershed
Cell_with_Nucleus
Virus_positive_Cells
Name
mask_Nucleus
DAPI1_LM
View original
Preprocessing
Remove Dark Background
Power
5
<Select>
Threshold
2500
65535
Smooth 1x
Clean 5x
Fill holes ON
Separate 2x
Size (µm)
7
inf
Circularity
0
1
Binary Processing
<Select>
Save Binary
Feature
Save
Mean
Count
Yes
321
<Select>
Preview
Run Now
Close

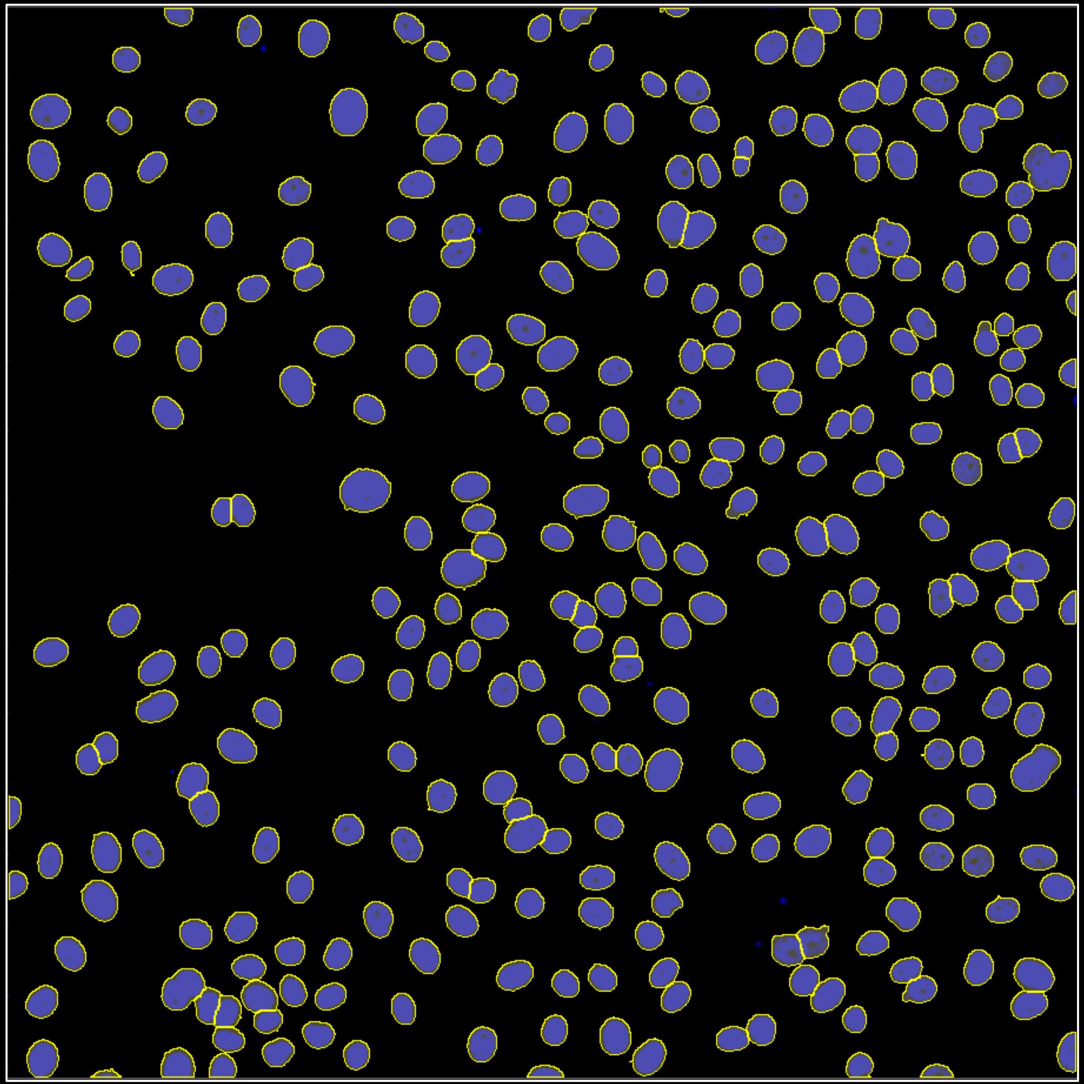

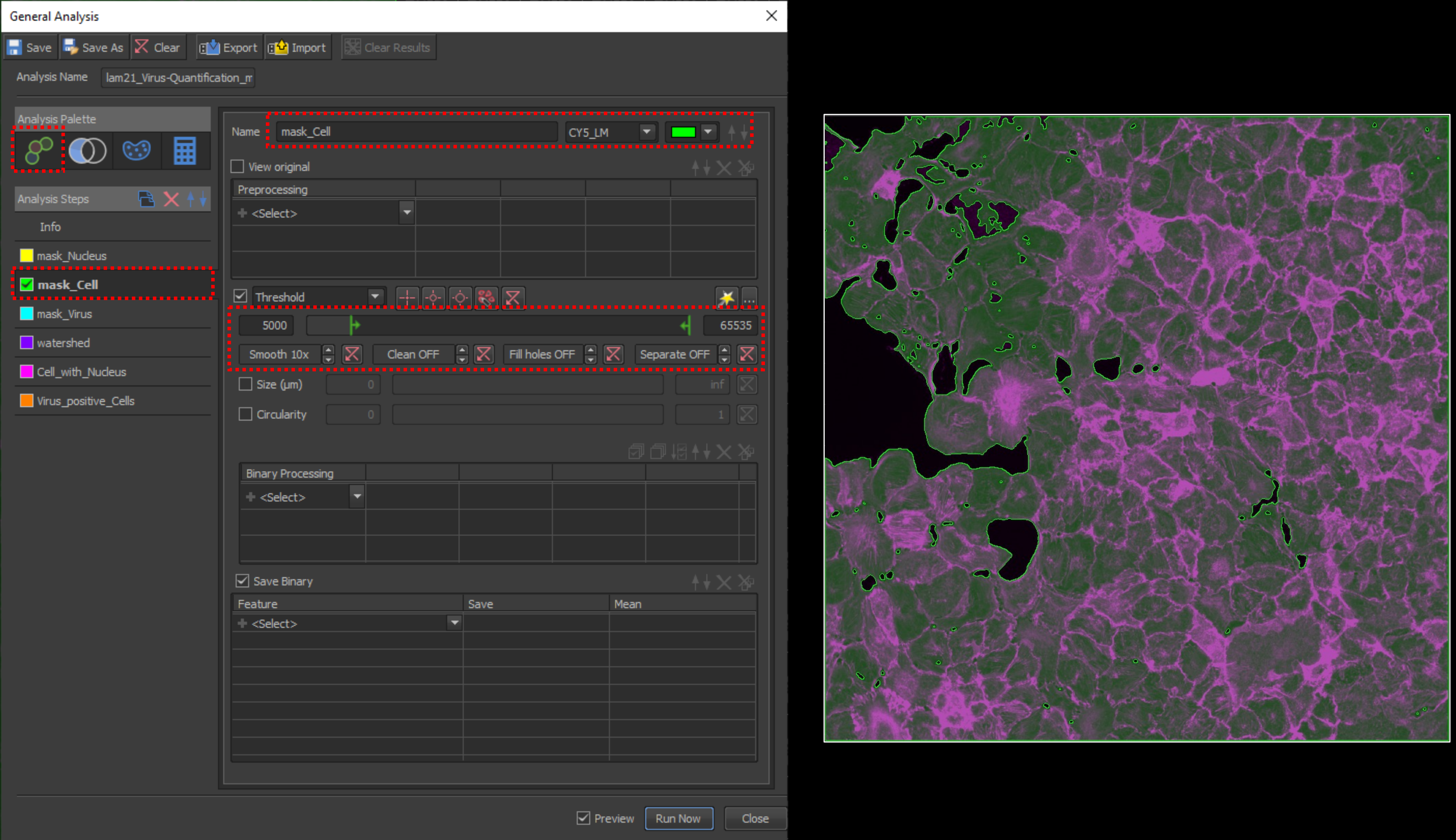
General Analysis
Save
Save As
Clear
Export
Import
Clear Results
Analysis Name
lam21_Virus-Quantification_m
Analysis Palette
Analysis Steps
Info
mask_Nucleus
mask_Cell
mask_Virus
watershed
Cell_with_Nucleus
Virus_positive_Cells
Name
mask_Cell
CY5_LM
View original
Preprocessing
<Select>
Threshold
5000
65535
Smooth 10x
Clean OFF
Fill holes OFF
Separate OFF
Size (µm)
0
inf
Circularity
0
1
Binary Processing
<Select>
Save Binary
Feature
Save
Mean
<Select>
Preview
Run Now
Close

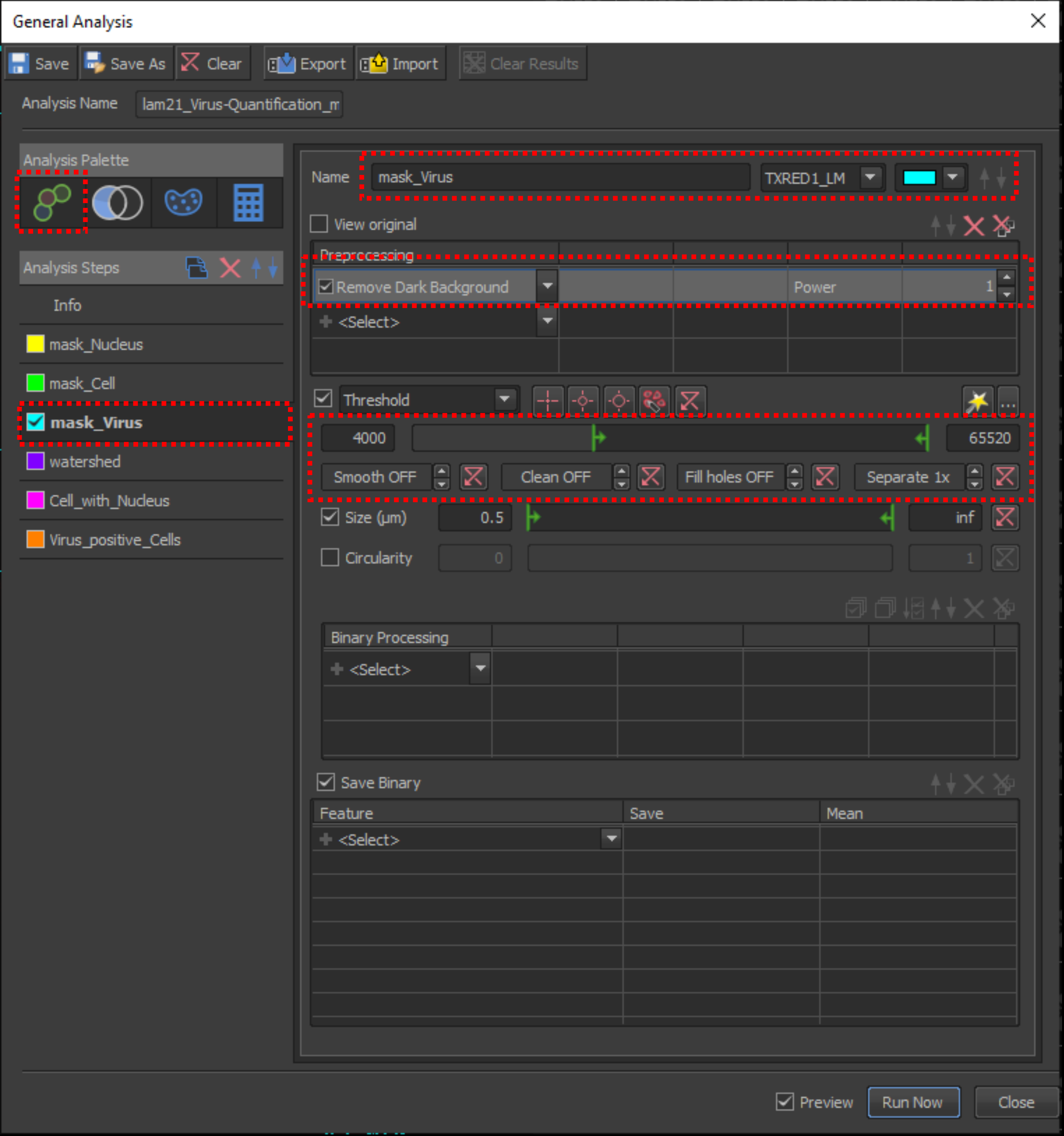
General Analysis
Save
Save As
Clear
Export
Import
Clear Results
Analysis Name
lam21_Virus-Quantification_m
Analysis Palette
Analysis Steps
Info
mask_Nucleus
mask_Cell
mask_Virus
watershed
Cell_with_Nucleus
Virus_positive_Cells
Name
mask_Virus
TXRED1_LM
View original
Preprocessing
Remove Dark Background
Power
1
<Select>
Threshold
4000
65520
Smooth OFF
Clean OFF
Fill holes OFF
Separate 1x
Size (µm)
0.5
inf
Circularity
0
1
Binary Processing
<Select>
Save Binary
Feature
Save
Mean
<Select>
Preview
Run Now
Close
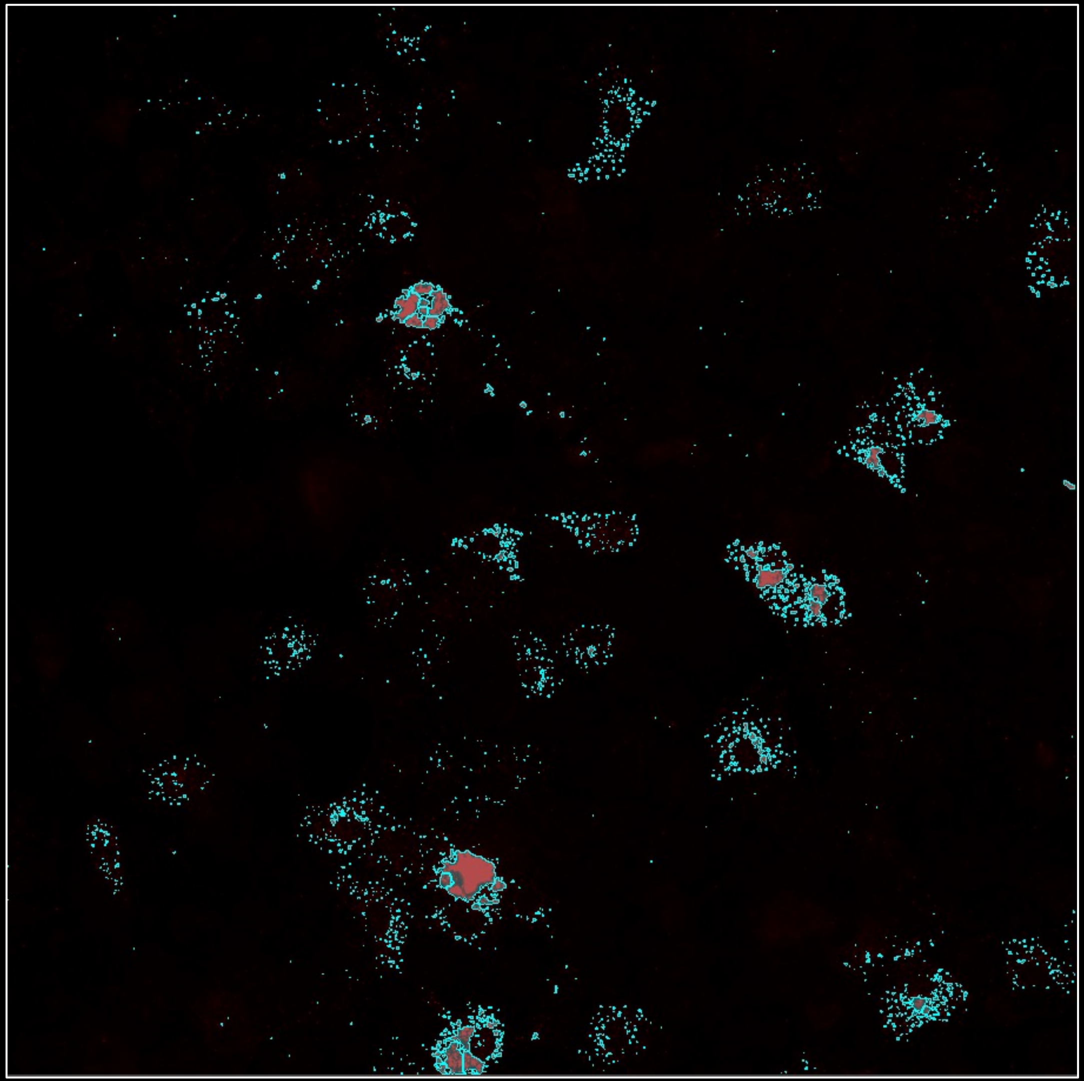

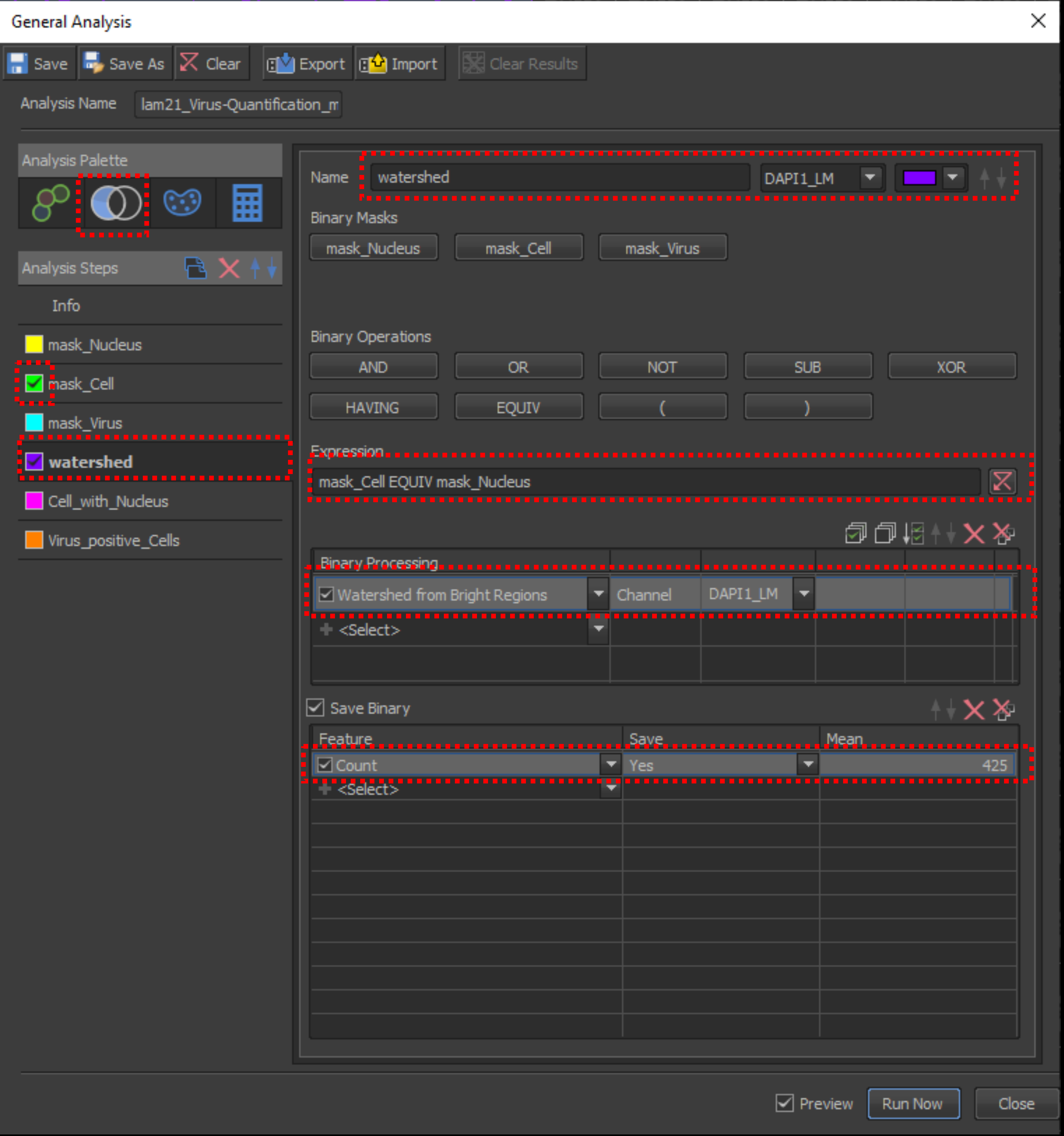
General Analysis
Save
Save As
Clear
Export
Import
Clear Results
Analysis Name
lam21_Virus-Quantification_m
Analysis Palette
Analysis Steps
Info
mask_Nucleus
mask_Cell
mask_Virus
watershed
Cell_with_Nucleus
Virus_positive_Cells
Name
watershed
DAPI1_LM
Binary Masks
mask_Nucleus
mask_Cell
mask_Virus
Binary Operations
AND
OR
NOT
SUB
XOR
HAVING
EQUIV
(
)
Expression
mask_Cell EQUIV mask_Nucleus
Binary Processing
Watershed from Bright Regions
Channel
DAPI1_LM
<Select>
Save Binary
Feature
Save
Mean
Count
Yes
425
<Select>
Preview
Run Now
Close

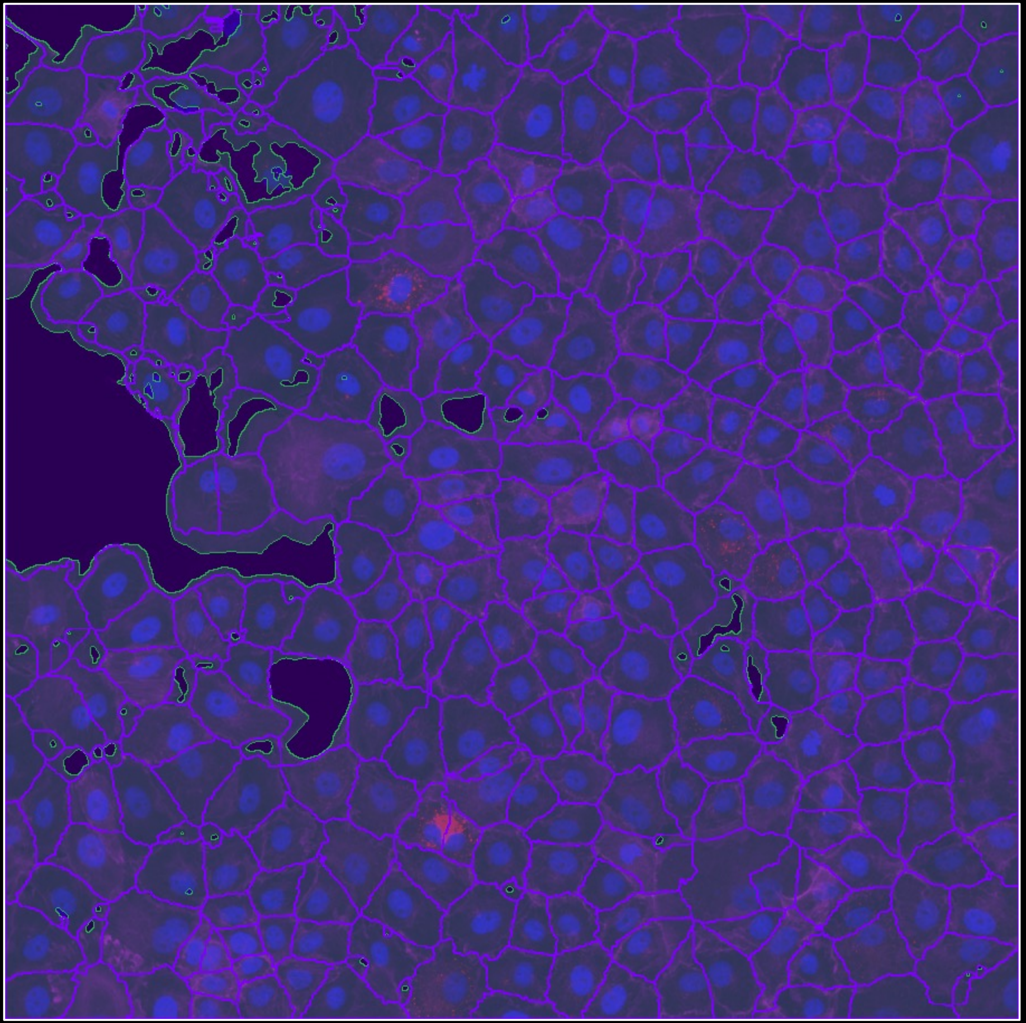

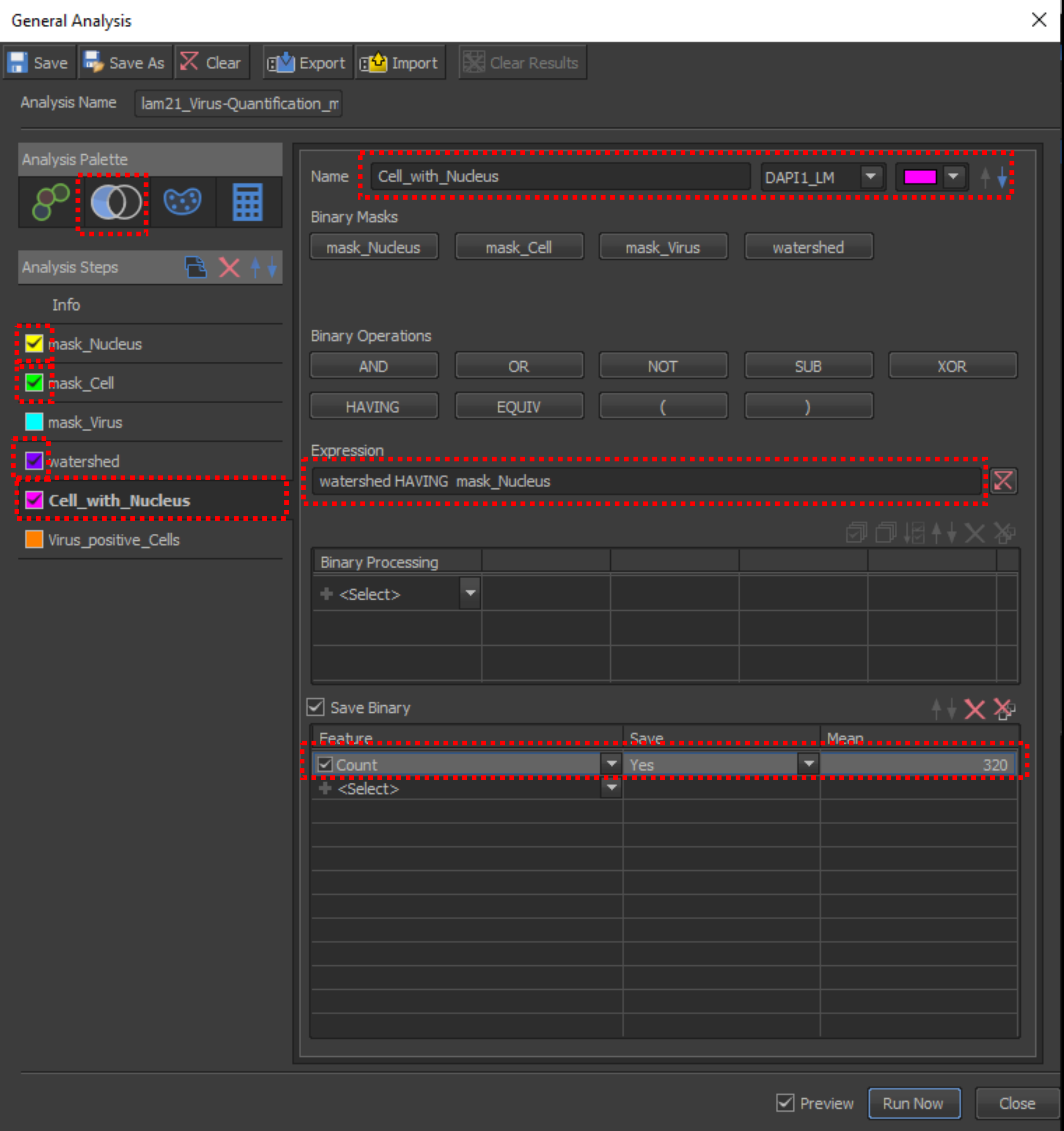
General Analysis
Save
Save As
Clear
Export
Import
Clear Results
Analysis Name
lam21_Virus-Quantification_m
Analysis Palette
Analysis Steps
Info
mask_Nucleus
mask_Cell
mask_Virus
watershed
Cell_with_Nucleus
Virus_positive_Cells
Name
Cell_with_Nucleus
DAPI1_LM
Binary Masks
mask_Nucleus
mask_Cell
mask_Virus
watershed
Binary Operations
AND
OR
NOT
SUB
XOR
HAVING
EQUIV
(
)
Expression
watershed HAVING mask_Nucleus
Binary Processing
<Select>
Save Binary
Feature
Save
Mean
Count
Yes
320
<Select>
Preview
Run Now
Close

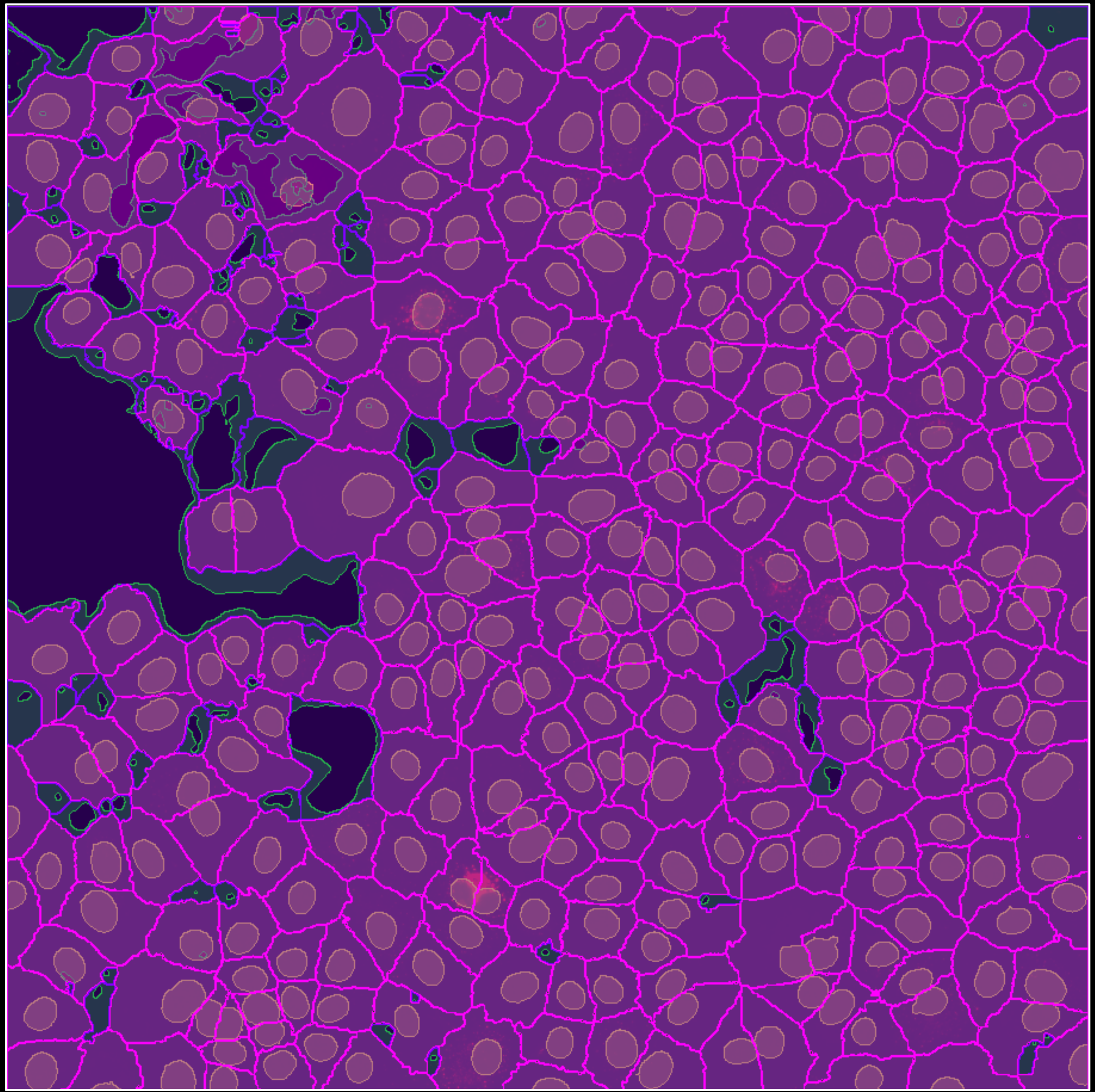

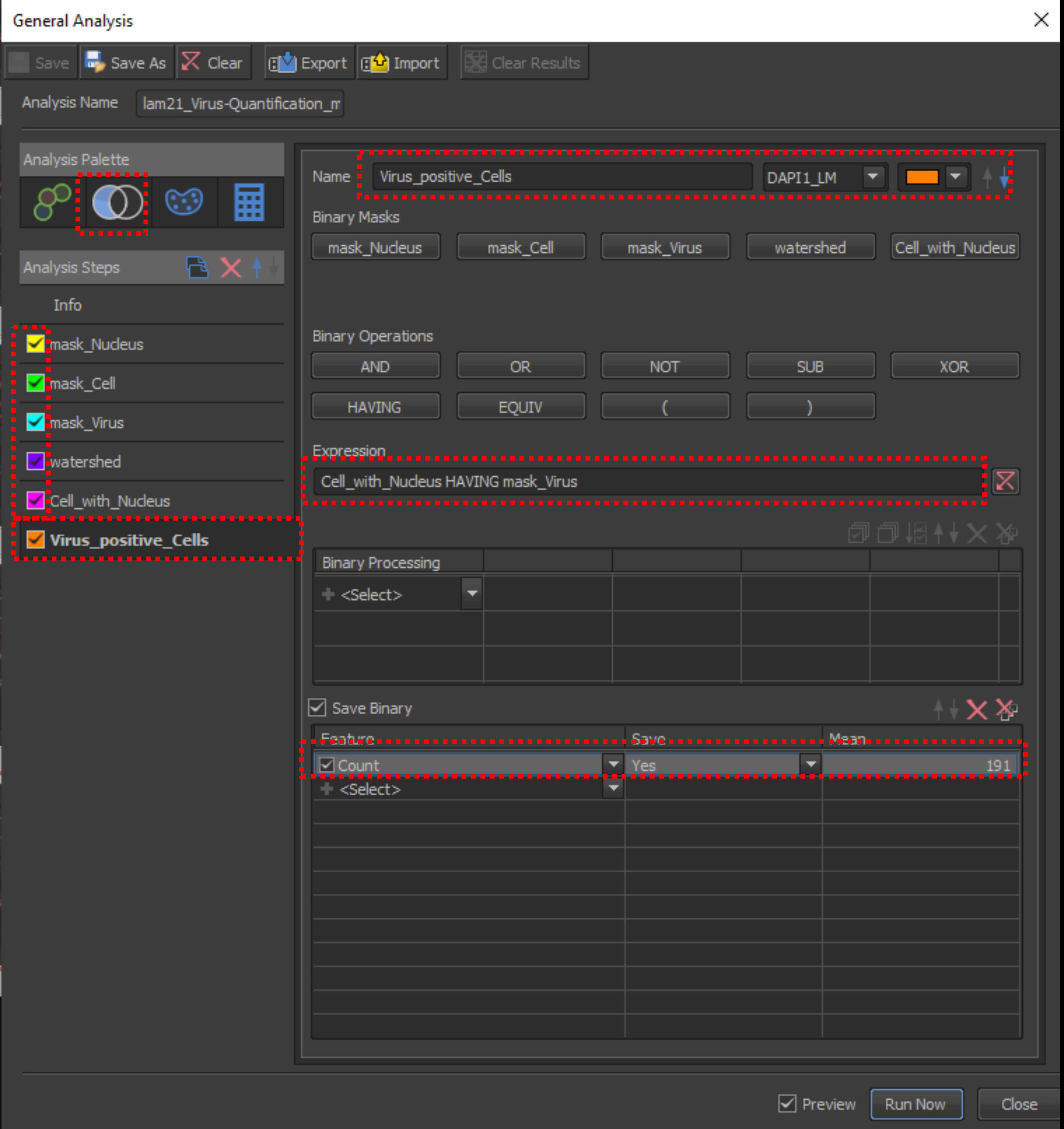
General Analysis
Save
Save As
Clear
Export
Import
Clear Results
Analysis Name
lam21_Virus-Quantification_m
Analysis Palette
Analysis Steps
Info
mask_Nucleus
mask_Cell
mask_Virus
watershed
Cell_with_Nucleus
Virus_positive_Cells
Name
Virus_positive_Cells
DAPI1_LM
Binary Masks
mask_Nucleus
mask_Cell
mask_Virus
watershed
Cell_with_Nucleus
Binary Operations
AND
OR
NOT
SUB
XOR
HAVING
EQUIV
(
)
Expression
Cell_with_Nucleus HAVING mask_Virus
Binary Processing
<Select>
Save Binary
Feature
Save
Mean
Count
Yes
191
<Select>
Preview
Run Now
Close

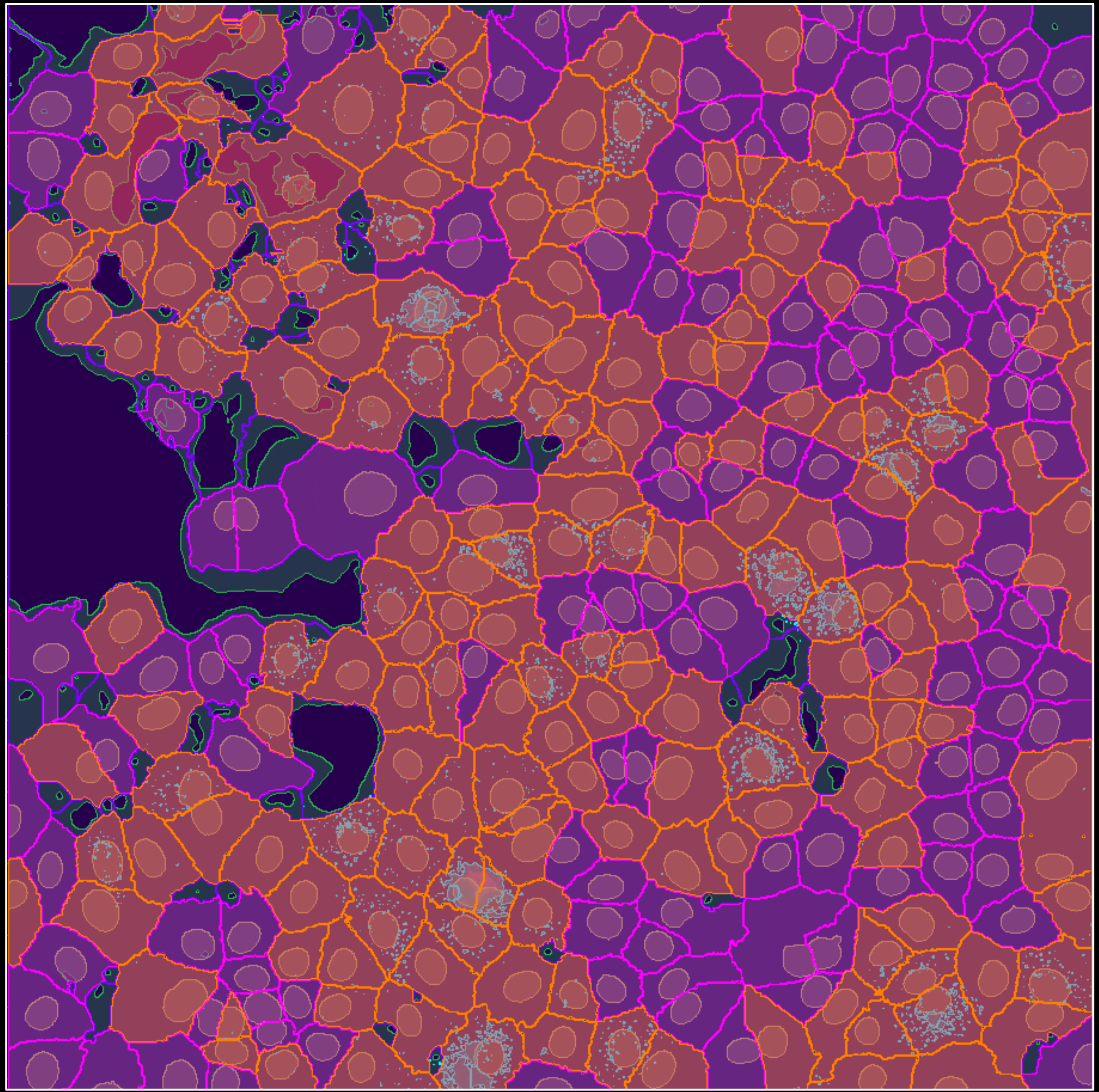

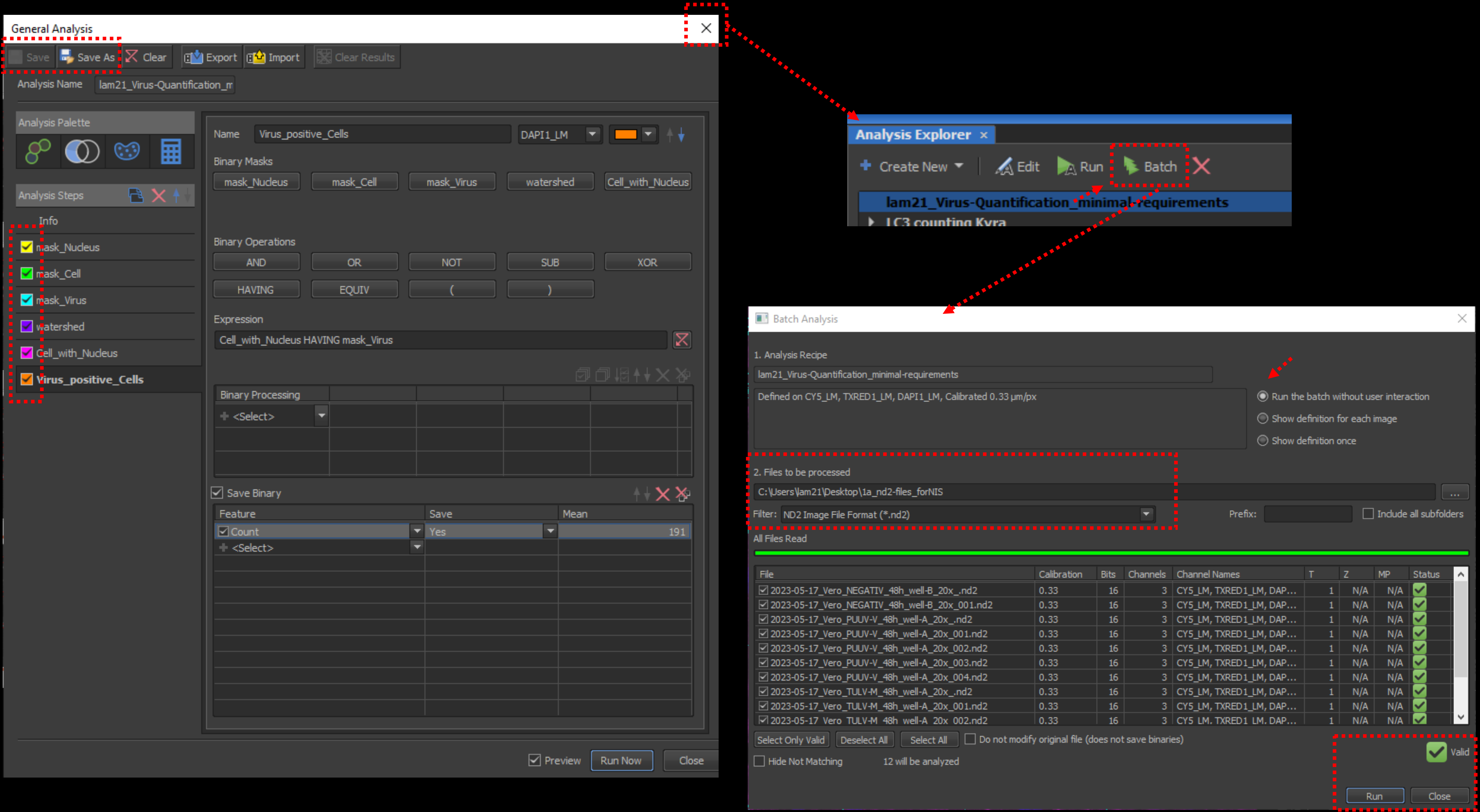
General Analysis
Save
Save As
Clear
Export
Import
Clear Results
Analysis Name
lam21_Virus-Quantification_m
Analysis Palette
Analysis Steps
Info
mask_Nucleus
mask_Cell
mask_Virus
watershed
Cell_with_Nucleus
Virus_positive_Cells
Name
Virus_positive_Cells
DAPI1_LM
Binary Masks
mask_Nucleus
mask_Cell
mask_Virus
watershed
Cell_with_Nucleus
Binary Operations
AND
OR
NOT
SUB
XOR
HAVING
EQUIV
(
)
Expression
Cell_with_Nucleus HAVING mask_Virus
Binary Processing
<Select>
Save Binary
Feature
Save
Mean
Count
Yes
191
<Select>
Preview
Run Now
Close
Analysis Explorer
Create New
Edit
Run
Batch
lam21_Virus-Quantification_minimal-requirements
LC3 counting Kyra
Batch Analysis
1. Analysis Recipe
lam21_Virus-Quantification_minimal-requirements
Defined on CY5_LM, TXRED1_LM, DAPI1_LM, Calibrated 0.33 µm/px
Run the batch without user interaction
Show definition for each image
Show definition once
2. Files to be processed
C:\Users\lam21\Desktop\1a_nd2-files_forNIS
Filter:
ND2 Image File Format (*.nd2)
Prefix:
Include all subfolders
All Files Read
File
Calibration
Bits
Channels
Channel Names
T
Z
MP
Status
2023-05-17_Vero_NEGATIV_48h_well-B_20x_.nd2
2023-05-17_Vero_NEGATIV_48h_well-B_20x_001.nd2
2023-05-17_Vero_PUUV-V_48h_well-A_20x_.nd2
2023-05-17_Vero_PUUV-V_48h_well-A_20x_001.nd2
2023-05-17_Vero_PUUV-V_48h_well-A_20x_002.nd2
2023-05-17_Vero_PUUV-V_48h_well-A_20x_003.nd2
2023-05-17_Vero_PUUV-V_48h_well-A_20x_004.nd2
2023-05-17_Vero_TULV-M_48h_well-A_20x_.nd2
2023-05-17_Vero_TULV-M_48h_well-A_20x_001.nd2
2023-05-17_Vero_TULV-M_48h_well-A_20x_002.nd2
0.33
16
3
CY5_LM, TXRED1_LM, DAP...
1
N/A
Select Only Valid
Deselect All
Select All
Do not modify original file (does not save binaries)
Valid
Hide Not Matching
12 will be analyzed
Run
Close

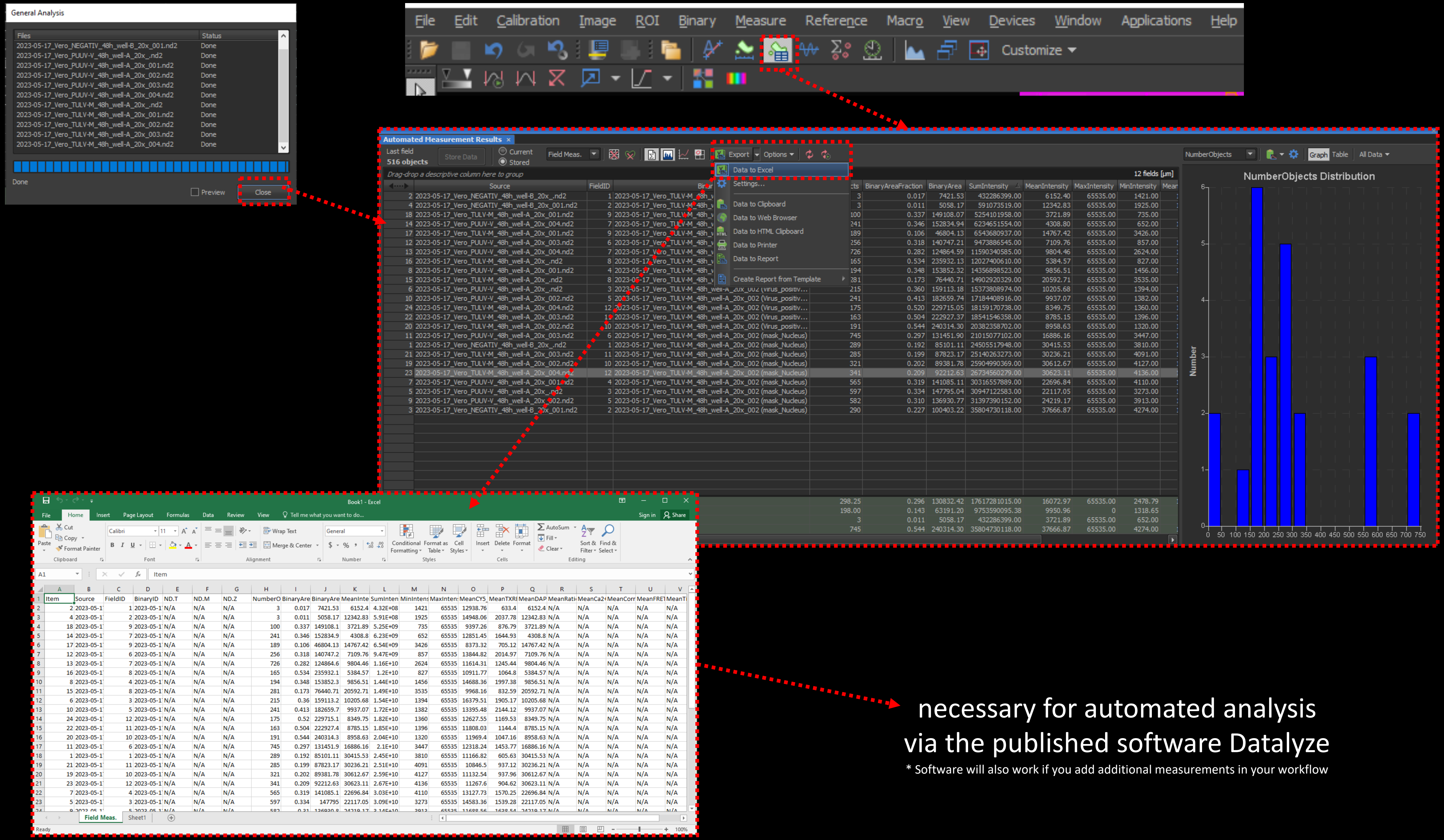


necessary for automated analysis
via the published software Datalyze

* Software will also work if you add additional measurements in your workflow

### Datalyze

**Software tool to analyze your data**

**Aim of this program:**

*sort necessary data for quantification to summarize and store data in a new file and show results graphically as a scatterplot and as a bar graph*

Raw data from NisElements:

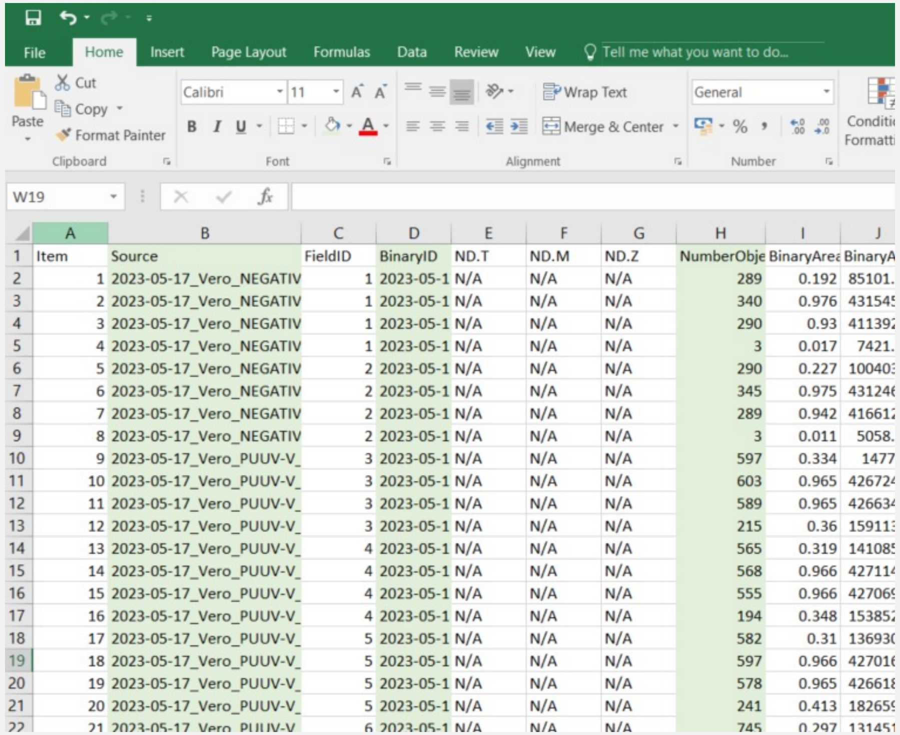

| Item | Source | FieldID | BinaryID | ND.T | ND.M | ND.Z | NumberObje | BinaryArea | BinaryA |
|---|---|---|---|---|---|---|---|---|---|
| 1 | 2023-05-17_Vero_NEGATIV | 1 | 2023-05-1 | N/A | N/A | N/A | 289 | 0.192 | 85101. |
| 2 | 2023-05-17_Vero_NEGATIV | 1 | 2023-05-1 | N/A | N/A | N/A | 340 | 0.976 | 431545 |
| 3 | 2023-05-17_Vero_NEGATIV | 1 | 2023-05-1 | N/A | N/A | N/A | 290 | 0.93 | 411392 |
| 4 | 2023-05-17_Vero_NEGATIV | 1 | 2023-05-1 | N/A | N/A | N/A | 3 | 0.017 | 7421. |
| 5 | 2023-05-17_Vero_NEGATIV | 2 | 2023-05-1 | N/A | N/A | N/A | 290 | 0.227 | 100403 |
| 6 | 2023-05-17_Vero_NEGATIV | 2 | 2023-05-1 | N/A | N/A | N/A | 345 | 0.975 | 431246 |
| 7 | 2023-05-17_Vero_NEGATIV | 2 | 2023-05-1 | N/A | N/A | N/A | 289 | 0.942 | 416612 |
| 8 | 2023-05-17_Vero_NEGATIV | 2 | 2023-05-1 | N/A | N/A | N/A | 3 | 0.011 | 5058. |
| 9 | 2023-05-17_Vero_PUUV-V_ | 3 | 2023-05-1 | N/A | N/A | N/A | 597 | 0.334 | 1477 |
| 10 | 2023-05-17_Vero_PUUV-V_ | 3 | 2023-05-1 | N/A | N/A | N/A | 603 | 0.965 | 426724 |
| 11 | 2023-05-17_Vero_PUUV-V_ | 3 | 2023-05-1 | N/A | N/A | N/A | 589 | 0.965 | 426634 |
| 12 | 2023-05-17_Vero_PUUV-V_ | 3 | 2023-05-1 | N/A | N/A | N/A | 215 | 0.36 | 159113 |
| 13 | 2023-05-17_Vero_PUUV-V_ | 4 | 2023-05-1 | N/A | N/A | N/A | 565 | 0.319 | 141085 |
| 14 | 2023-05-17_Vero_PUUV-V_ | 4 | 2023-05-1 | N/A | N/A | N/A | 568 | 0.966 | 427114 |
| 15 | 2023-05-17_Vero_PUUV-V_ | 4 | 2023-05-1 | N/A | N/A | N/A | 555 | 0.966 | 427069 |
| 16 | 2023-05-17_Vero_PUUV-V_ | 4 | 2023-05-1 | N/A | N/A | N/A | 194 | 0.348 | 153852 |
| 17 | 2023-05-17_Vero_PUUV-V_ | 5 | 2023-05-1 | N/A | N/A | N/A | 582 | 0.31 | 136930 |
| 18 | 2023-05-17_Vero_PUUV-V_ | 5 | 2023-05-1 | N/A | N/A | N/A | 597 | 0.966 | 427016 |
| 19 | 2023-05-17_Vero_PUUV-V_ | 5 | 2023-05-1 | N/A | N/A | N/A | 578 | 0.965 | 426618 |
| 20 | 2023-05-17_Vero_PUUV-V_ | 5 | 2023-05-1 | N/A | N/A | N/A | 241 | 0.413 | 182659 |
| 21 | 2023-05-17_Vero_PUUV-V_ | 6 | 2023-05-1 | N/A | N/A | N/A | 745 | 0.297 | 131451 |

Keep important columns and process data

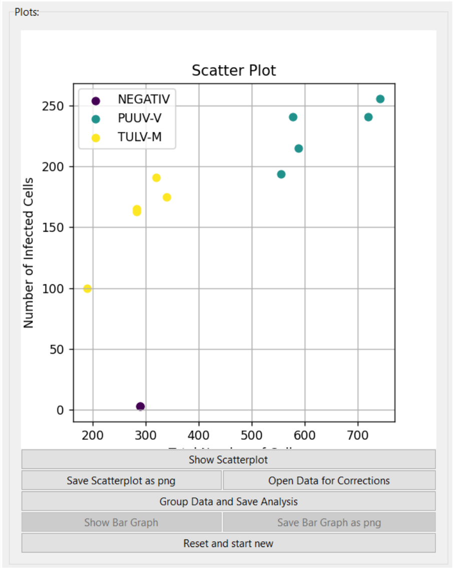


| Source | Percentage of Infected Cells [%] | Total Number of Cells | Number of Infected Cells |
|---|---|---|---|
| NEGATIV | 1.03 | 290 | 3 |
| NEGATIV | 1.04 | 289 | 3 |
| PUUV-V | 36.5 | 589 | 215 |
| PUUV-V | 34.95 | 555 | 194 |
| PUUV-V | 41.7 | 578 | 241 |
| PUUV-V | 34.45 | 743 | 256 |
| PUUV-V | 33.47 | 720 | 241 |
| TULV-M | 58.3 | 283 | 165 |
| TULV-M | 52.63 | 190 | 100 |
| TULV-M | 59.69 | 320 | 191 |
| TULV-M | 57.6 | 283 | 163 |
| TULV-M | 51.62 | 339 | 175 |

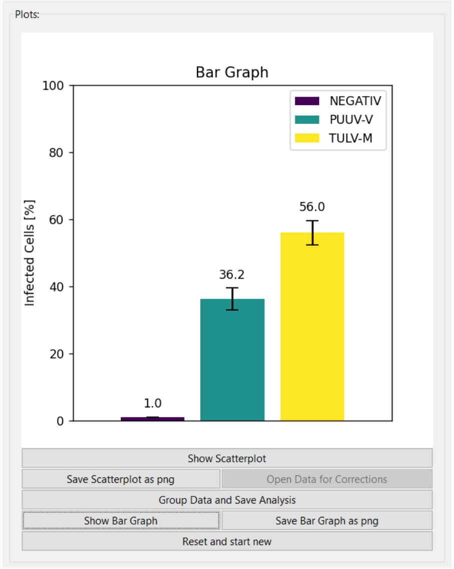


| Source | Mean Percentage of Infected Cells [%] | Standard Deviation (SD) |
|---|---|---|
| NEGATIV | 1.04 | 0.01 |
| PUUV-V | 36.21 | 3.26 |
| TULV-M | 55.97 | 3.61 |

Path of file: ***Copy and Paste file path as text***

Name of file: ***Copy and Paste file name***

Name for new file: ***Enter the Name for the next document – it will be saved in the same folder as your loaded file***

- The "Load Columns from File" button also deletes everything after "Feature"
- software recognizes the columns "Source", "BinaryID" and "NumberObjects" automatically by names/strings

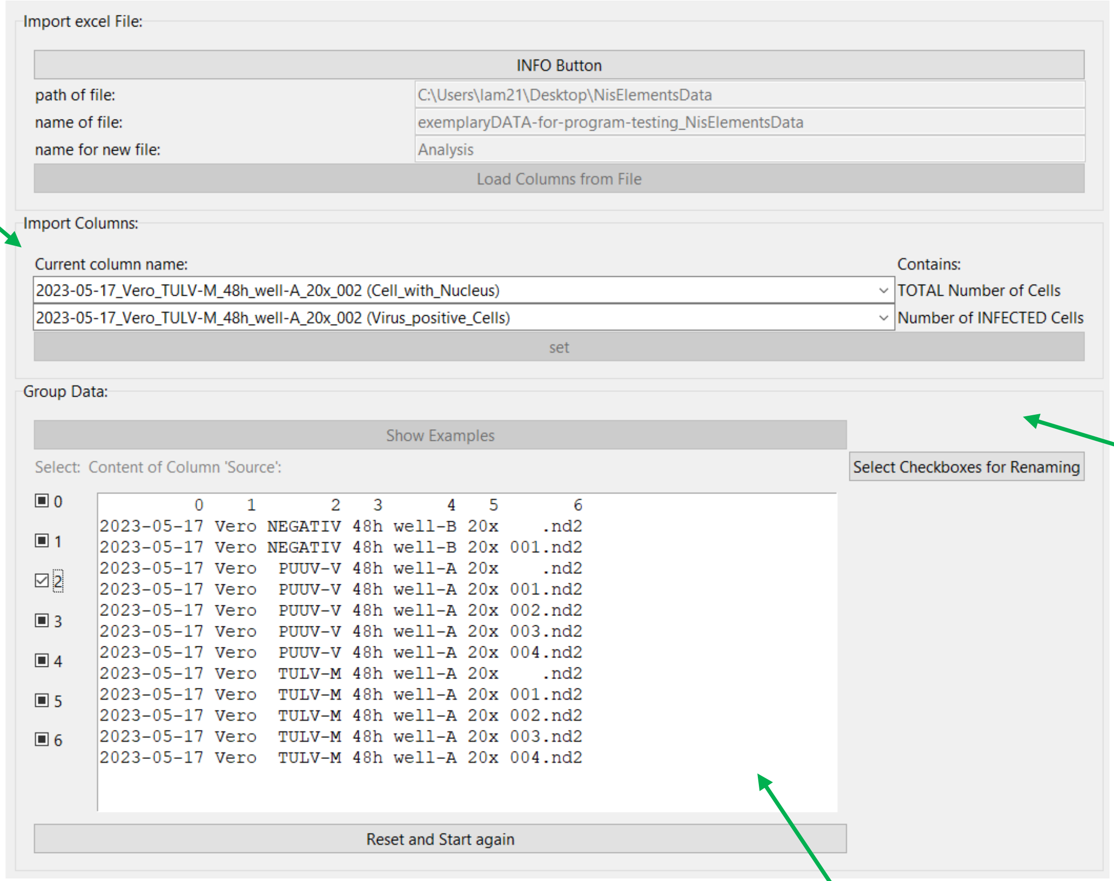


| Source | BinaryID | NumberObjects |
|---|---|---|
| 2023-05-17_Vero_PUUV-V_48h_well-A_20x_001.nd2 | 2023-05-17_Vero_PUUV-V_48h_well-A_20x_003 (Virus_positive_Cells) | 194 |
| 2023-05-17_Vero_PUUV-V_48h_well-A_20x_001.nd2 | 2023-05-17_Vero_PUUV-V_48h_well-A_20x_003 (Cell_with_Nucleus) | 568 |
| 2023-05-17_Vero_PUUV-V_48h_well-A_20x_002.nd2 | 2023-05-17_Vero_PUUV-V_48h_well-A_20x_003 (Virus_positive_Cells) | 241 |
| 2023-05-17_Vero_PUUV-V_48h_well-A_20x_002.nd2 | 2023-05-17_Vero_PUUV-V_48h_well-A_20x_003 (Cell_with_Nucleus) | 597 |

To proceed the data you got from NisElements, this software pivots the excel table.
Please enter the correct BinaryID / mask. Only these two selected will be considered and processed.

| Name | Number of INFECTED Cells | TOTAL Number of Cells |
|---|---|---|
| 2023-05-17_Vero_PUUV-V_48h_well-A_20x_001.nd2 | 194 | 568 |
| 2023-05-17_Vero_PUUV-V_48h_well-A_20x_002.nd2 | 241 | 597 |

Program creates data sets from names. For this you need to select the relevant information to group your data according to useful sets: e.g. keep relevant information like which strain you want to compare (PUUV/TULV), but get rid of name parts, which prevent the code of grouping your data (e.g. image numberings -001.nd2)

| Original | Delete letters: | Group data for MEAN and SD |
|---|---|---|
| 2023-05-17_Vero_PUUV-V_48h_well-A_20x_.nd2 | 2023-05-17_Vero_PUUV-V_48h_well-A_20x_000.nd2 | PUUV-V |
| 2023-05-17_Vero_PUUV-V_48h_well-A_20x_001.nd2 | 2023-05-17_Vero_PUUV-V_48h_well-A_20x_001.nd2 | |
| 2023-05-17_Vero_PUUV-V_48h_well-A_20x_002.nd2 | 2023-05-17_Vero_PUUV-V_48h_well-A_20x_002.nd2 | |
| 2023-05-17_Vero_TULV-M_48h_well-A_20x_.nd2 | 2023-05-17_Vero_TULV-M_48h_well-A_20x_000.nd2 | TULV-M |
| 2023-05-17_Vero_TULV-M_48h_well-A_20x_001.nd2 | 2023-05-17_Vero_TULV-M_48h_well-A_20x_001.nd2 | |
| 2023-05-17_Vero_TULV-M_48h_well-A_20x_002.nd2 | 2023-05-17_Vero_TULV-M_48h_well-A_20x_002.nd2 | |

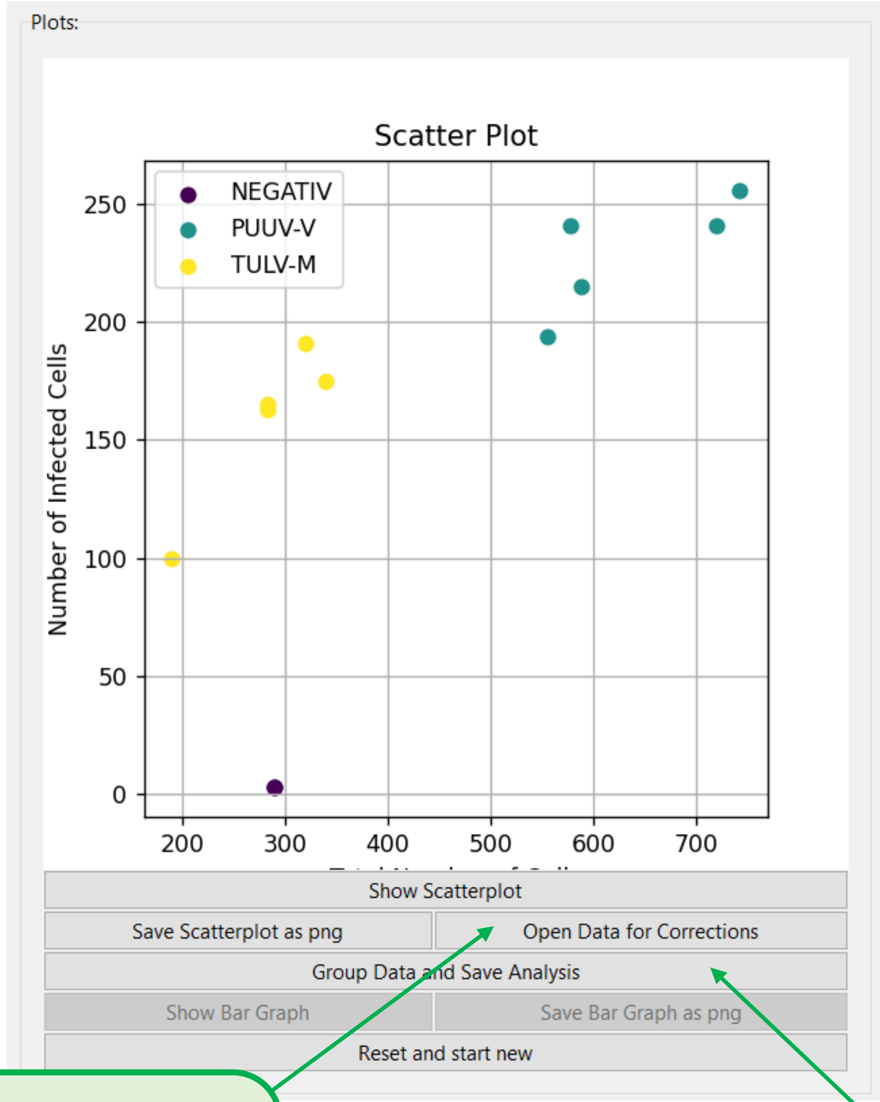


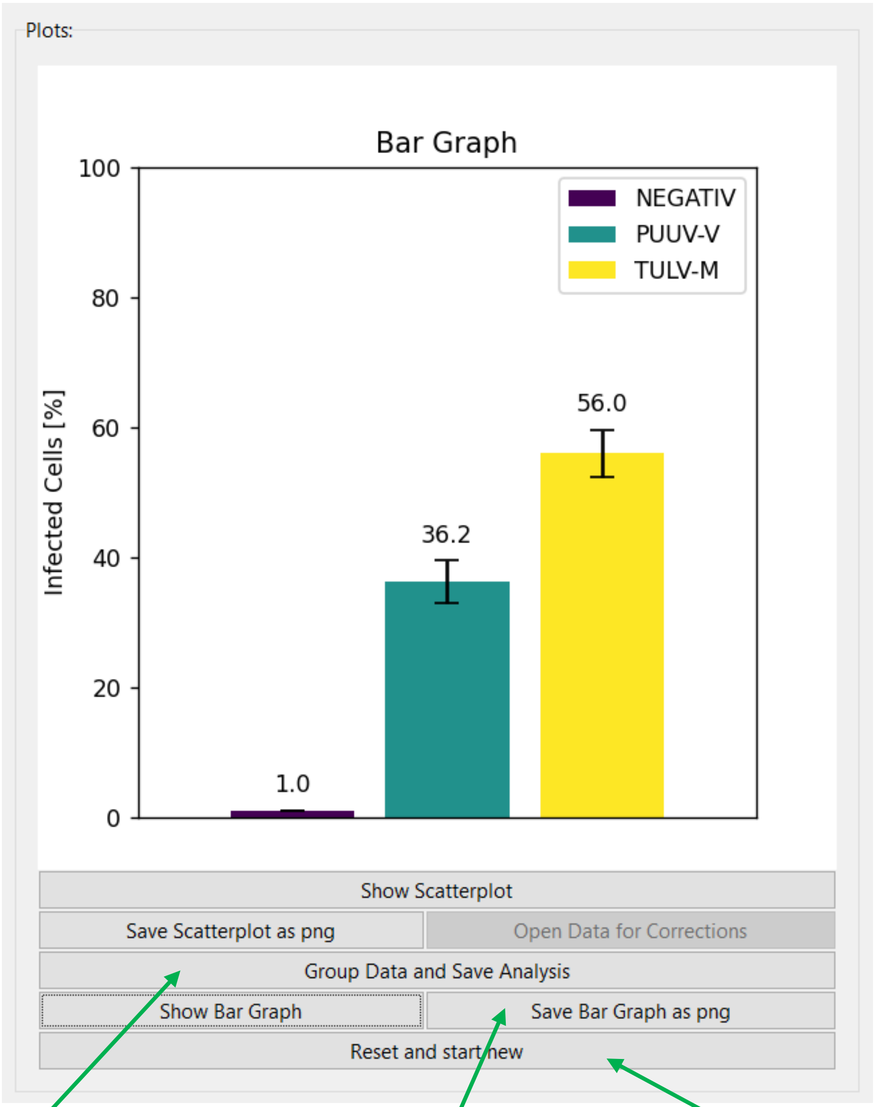


**< Open Data for Corrections > opens your new excel sheet, so that you could delete outliers. Make sure that you save the excel sheet without changing the file name and close it properly. After closing the document you can update your scatterplot by clicking again on < Show Scatterplot >**

**< Group Data and Save Analysis > will then convert and summarize all the grouped data and store it as a new sheet in the document.**

**< Save … as png > PNG will be saved in the same folder, which currently stores your loaded/saved file.**

**< Reset and start new > Software will switch back to the first window and enables the start of a new analysis**
